# GraviKit: an easy-to-implement microscope add-on for observation of gravitation dependent processes

**DOI:** 10.1101/2021.08.30.458259

**Authors:** Christian Feldhaus, Martina Kolb, Michelle Küppers, Steffen Hardy, Ralph Palmisano

## Abstract

One of the most important environmental cues for living organisms is gravity and many developmental processes depend on it. However, when it comes to light microscopy, a majority of studies on these processes work with their objects of interest placed perpendicular to their natural orientation. One reason for that is probably that light microscopes with the required horizontal beampath are either costly or require advanced technical skills. To circumvent these obstacles and make imaging of gravity-dependent processes with a horizontal beampath possible for any lab we developed GraviKit. It converts a standard inverted research microscope into an imaging device with a horizontal beampath with a stage that rotates the sample around the optical axis. Like this, the direction of gravity can be freely chosen during an imaging experiment. The system is easy to implement and suitable for multi-user environments.

## Introduction

Life on earth has adapted to all sorts of environmental conditions, most of which have been changing during the course of evolution. However, one major environmental cue has remained constant during the history of life, which is gravity (Herranz & Medina 2013). Sophisticated gravity sensing mechanisms have evolved which help living organisms developing and maintaining their physical shape (Takahashi et al. 2021), enabling them to use available resources in the most efficient way to ensure survival and reproduction.

Especially sessile land plants heavily rely during their development on gravitational cues to correctly build up their supporting tissue so they can reach their resources down in the soil (water, nutrients) and up above the ground (sunlight) (Volkmann & Baluska 2006). They are able to even sense minor changes gravitational forces (Herranz & Medina 2013). One of the molecular key players in mediating gravitational cues is Auxin signalling. For this small molecule there is emerging evidence, that important signalling events already take place within less than a minute after altering the direction of gravity (Herud-Sikimic et al. 2020). Therefore it would be of importance to take this into account when designing future experiments dealing with gravitation dependent processes in plants, e.g. concerning the orientation of plants during those experiments.

Light microscopy has been established as the most important tool to visualize a plethora of processes in living plants under defined environmental conditions in laboratories worldwide. One common interest is in understanding the physiological changes and responses of plants on numerous processes that depend on gravitational forces. As said, observation of such gravitation dependent processes in living specimen, e.g. plants, on the levels of organs or cells usually involves the use of a light microscope. However, standard upright or inverted microscope designs do not allow for such observations under ideal circumstances as the sample needs to be oriented perpendicular to the gravitational field of our planet (Barnes 1896, Ovecka et al. 2018). While this bears the risk of observing artefacts when dealing with gravitation dependent processes, still more than 90% of all studies observing intact living roots on a microscope use standard setups without the possibility of putting the plant in its natural orientation (see literature search in suppl. data).

One probable reason is that there is no solution around which could be easily implemented for microscopes in any plant lab or in a multi-user environment. Current solutions for dedicated vertical imaging - and especially gravitational imaging - are either costly (von Wangenheim et al. 2017, Hemmersbach et al. 2006) or, if you aim for a home-built optical system, require quite some technical skills to implement (Maizel et.al. 2011). Solutions for converting existing inverted setups e.g. with a periscope (Monshausen et al. 2011) also require technical skills and time to set up. One further runs the risk of damaging the system (periscope can crash into stage) and it’s also not necessarily affordable for every lab.

Our aim was to build a microscope add-on that

- would allow to place the sample in a vertical position and rotate it around the horizontal axis (= optical axis of the microscope)
- would fit on most commercially available inverted research grade microscopes
- could be installed and de-installed by any person operating the microscope (= no advanced technical skills necessary) in less than 30 minutes
- is below 1000€ (manual rotation) or 2000€ (motorised rotation)
- is built with off-the-shelf components with world-wide availability

Here we present the GraviKit, an easy to implement solution that meets the above mentioned requirements. It consists of an objective holder replacing the condenser of an inverted microscope stand and a stage adaptor for vertical mounting of the sample on a standard multiwell plate holder.

## Methods

### Construction of the objective holder and alignment tool

The objective holder was constructed from Thorlabs parts as depicted in Figure 1. A complete parts list for inverted microscopes from several manufacturers you can find in Table 1. The objective holder replaces the condenser of the inverted microscope. The condenser focussing drive is repurposed as lateral displacement drive for the GraviKit objective.

**Figure 1.**
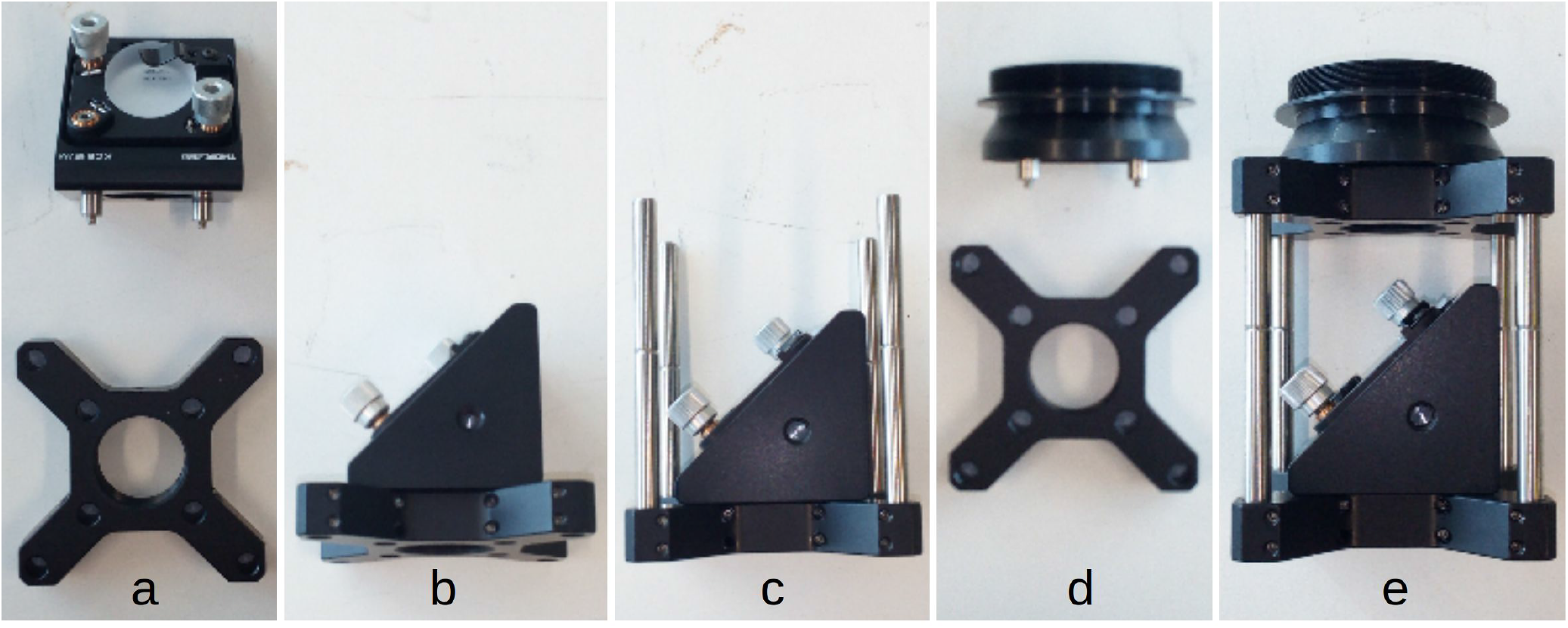
shows the assembly of the objective holder from the parts listed in Table 1. The example given is for the assembly of an objective holder for an Olympus IX-83. Please see text for further mounting instructions.

**Table 1.**
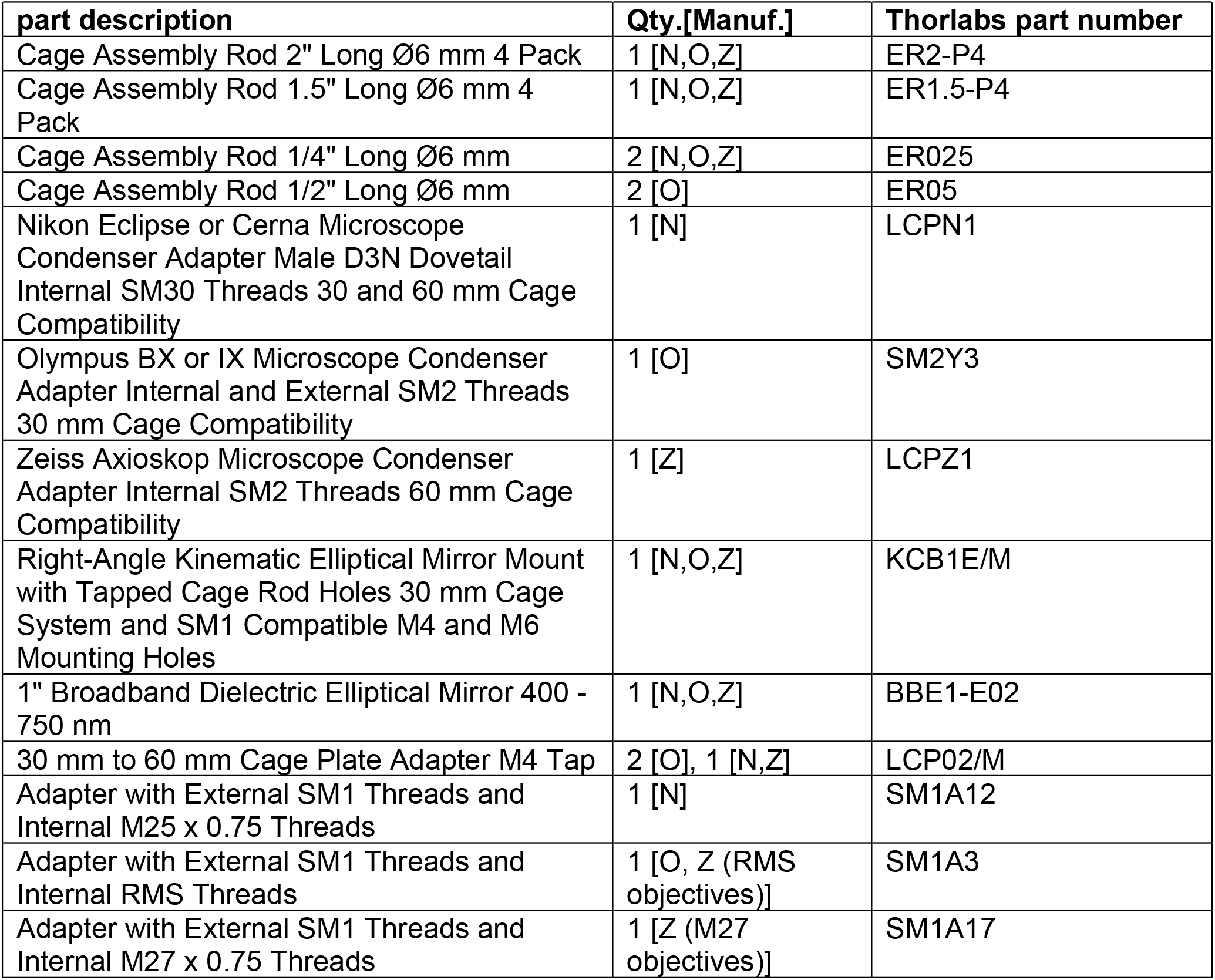
Thorlabs parts list for building the objective holder: 1st column shows the parts description, 2nd column tells the quantity of parts you need for a setup from a given manufacturer (N - Nikon, O - Olympus, Z - Zeiss), 3rd column holds the part number.

| part description | Qty.[Manuf.] | Thorlabs part number |
| --- | --- | --- |
| Cage Assembly Rod 2" Long Ø6 mm 4 Pack | 1 [N,O,Z] | ER2-P4 |
| Cage Assembly Rod 1.5" Long Ø6 mm 4 Pack | 1 [N,O,Z] | ER1.5-P4 |
| Cage Assembly Rod 1/4" Long Ø6 mm | 2 [N,O,Z] | ER025 |
| Cage Assembly Rod 1/2" Long Ø6 mm | 2 [O] | ER05 |
| Nikon Eclipse or Cerna Microscope Condenser Adapter Male D3N Dovetail Internal SM30 Threads 30 and 60 mm Cage Compatibility | 1 [N] | LCPN1 |
| Olympus BX or IX Microscope Condenser Adapter Internal and External SM2 Threads 30 mm Cage Compatibility | 1 [O] | SM2Y3 |
| Zeiss Axioskop Microscope Condenser Adapter Internal SM2 Threads 60 mm Cage Compatibility | 1 [Z] | LCPZ1 |
| Right-Angle Kinematic Elliptical Mirror Mount with Tapped Cage Rod Holes 30 mm Cage System and SM1 Compatible M4 and M6 Mounting Holes | 1 [N,O,Z] | KCB1E/M |
| 1" Broadband Dielectric Elliptical Mirror 400 - 750 nm | 1 [N,O,Z] | BBE1-E02 |
| 30 mm to 60 mm Cage Plate Adapter M4 Tap | 2 [O], 1 [N,Z] | LCP02/M |
| Adapter with External SM1 Threads and Internal M25 x 0.75 Threads | 1 [N] | SM1A12 |
| Adapter with External SM1 Threads and Internal RMS Threads | 1 [O, Z (RMS objectives)] | SM1A3 |
| Adapter with External SM1 Threads and Internal M27 x 0.75 Threads | 1 [Z (M27 objectives)] | SM1A17 |

How to build the objective holder:

1. Place the broadband dielectric mirror in the kinematic mirror mount (Fig. 1 a).
2. Fix the kinematic mirror mount on a cage plate adaptor using 2 of the 1/4” cage assembly rods (Fig. 1 a & b).
3. Connect each of the 1.5” cage assembly rods with one of the 2” cage assembly rods.
4. Place each of the built 3.5” rods in one of the corner holes of the cage plate adaptor holding the kinematic mirror mount (Fig. 1 c).
5. Olympus only: connect the microscope condenser adaptor with the unused cage plate adaptor using the 1/2” cage assembly rods (Fig. 1 d).
6. Place the microscope condenser adaptor (on Olympus connected to the cage plate adaptor) on the free ends of the 3.5” rods (Fig. 1 e).
7. Replace the condenser of your inverted microscope with the objective holder (!!! please make sure you don’t just unplug a motorised condenser, this might damage the electronics of your microscope, ask the microscope manufacturer how to safely remove a motorised condenser).
8. For calibration: please see below
9. For imaging: mount your objective on the free opening of the kinematic mirror holder using the appropriate thread adaptor

The alignment tool for the objective holder was constructed from Thorlabs and Fischertechnik parts as depicted in Figure 2. A parts list is contained in Table 2.

**Figure 2.**
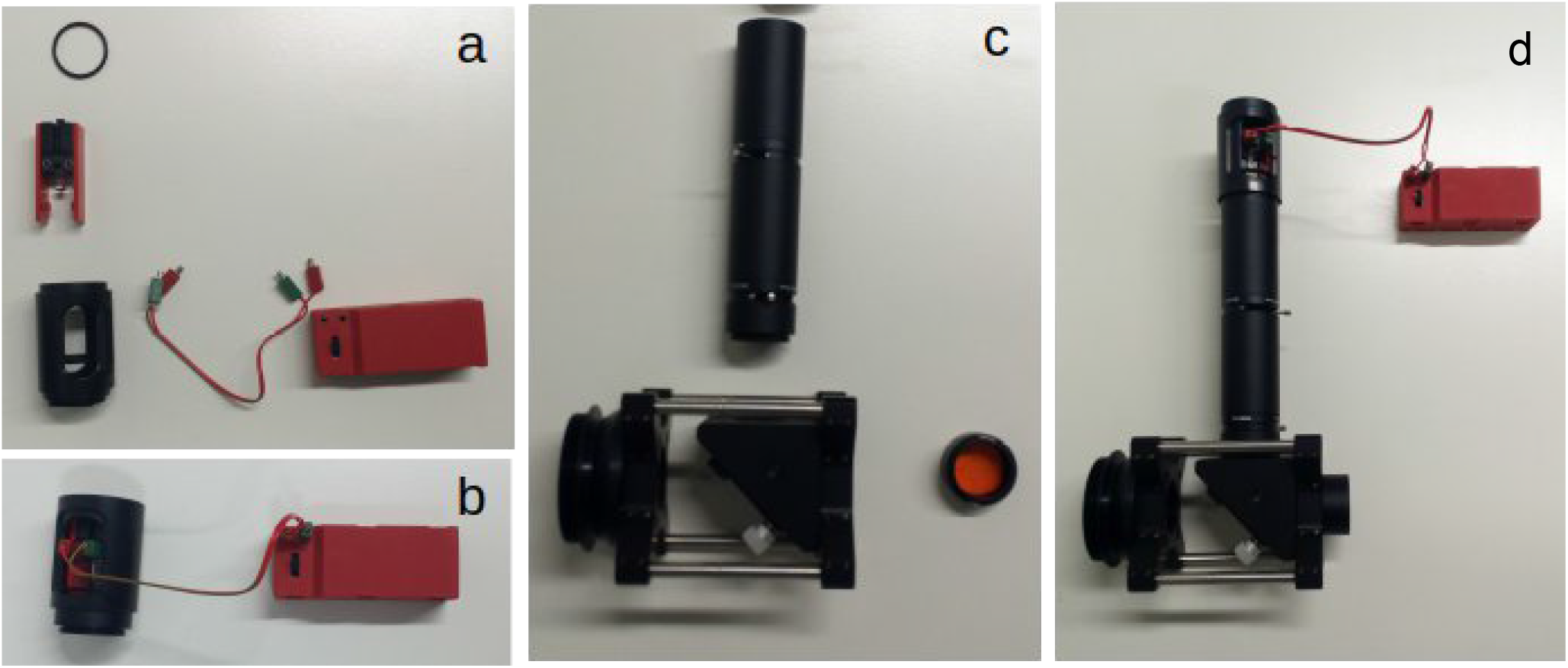
(a) Disassembled and (b) assembled view of the illumination unit of the calibration tool built from a Thorlabs slotted tube with retainer ring and Fischertechnik battery powered LED and construction parts. (c) Assembled view of the other parts of the calibration tool (tube and alignment disc) next to the corresponding mounting holes of the objective holder. (d) Calibration tool mounted on the objective holder.

**Table 2.** Parts list for building the calibration tool: 1st column shows the parts description, 2nd column gives the quantity of parts you need, 3rd column holds the parts number. Upper part of the table contains the Thorlabs parts, lower part the Fischertechnik parts.

| Thorlabs part description | Qty. | Thorlabs part number |
| --- | --- | --- |
| SM1 Lever-Actuated Iris Diaphragm (Ø0.8 - Ø12 mm) | 2 | SM1D12 |
| SM1 Slotted Lens Tube, 2" Thread Depth, 2 Retaining Rings Included | 1 | SM1L20C |
| SM1 Lens Tube, 2.00" Thread Depth, One Retaining Ring Included | 2 | SM1L20 |
| Fluorescent Alignment Disk Green | 1 | ADF2 |
| SM1 Lens Tube 0.50" Thread Depth One Retaining Ring Included | 2 | SM1L05 |
| **Fischertechnik part description** | **Qty.** | **Fischertechnik part number** |
| building block 15, black | 1 | 32881 |
| mounting plate 15x45, black or red | 2 | 38261 or 38242 |
| lead 250, red/green | 1 | 36210 |
| battery holder (9V), red | 1 | 135719 |
| flat plug, green | 2 | 31336 |
| flat plug, red | 2 | 31337 |
| LED white assembled 9v, 0.01A | 1 | 162134 |

How to build the alignment tool:

1. Put together Fischertechnik (Ft) building block 15, the two mounting plates and the LED as depicted in Fig. 2a.
2. Fix the four flat plugs at the ends of the lead and put a 9V battery in the battery holder.
3. Take one of the Thorlabs retainer rings and move it down the slotted lens tube as shown in Fig. 2a.
4. Slide the Ft LED holder into the slotted lens tube and clamp the mounting plates in the retainer ring (Fig. 2b, LED should be facing towards the end of the slotted lens tube with the external thread).
5. Fix the LED holder from the other side with another retainer ring.
6. Connect the battery power pack with the LED via the lead through the slot on the lens tube (Fig. 2b).
7. The tube to connect the LED holder with the objective holder is made from Thorlabs parts in the following order: 0.5” lens tube (external thread facing objective holder), iris diaphragm, 2” lens tube, 2” lens tube, iris diaphragm (internal thread facing LED holder) (Fig. 2c).
8. Fix the fluorescent alignment disc in the remaining 0.5” lens tube using retainer rings (Fig. 2c).
9. Put together the objective holder, tube and LED holder (the tube goes into the thread which will later hold the objective adaptor, Fig. 2d).
10. Screw the lens tube holding the fluorescent disc in the bottom opening of the objective holder which will later face towards the opening in the objective turret of the microscope (Fig. 2d).

### Mounting and alignment of the objective holder

1. Remove the microscope condenser according to the manufacturer’s manual (for motorised condensers: microscope has to be switched off during this procedure).
2. Remove the objective in the beam path.
3. Place the objective holder with the condenser adaptor ring in the condenser holder with the opening for the objective facing sidewards (tighten all the screws properly, but not to strong!).
4. Screw the alignment tool in the objective opening (as shown in Fig. 2d).
5. Switch on the LED and close the shutter of the calibration tool.
6. Remove ocular on one side, you will now see the alignment disc.
7. Iteratively center the alignment disc with the adjustment screws on the objective holder and the Köhler screws of the microscope while opening and closing the shutters (Fig. 3).
8. When done, remove calibration tool (tube and alignment disc).
9. Screw in the desired objective with the appropriate adaptor.
10. Configure the objective you are using in the microscope software.

**Figure 3.**
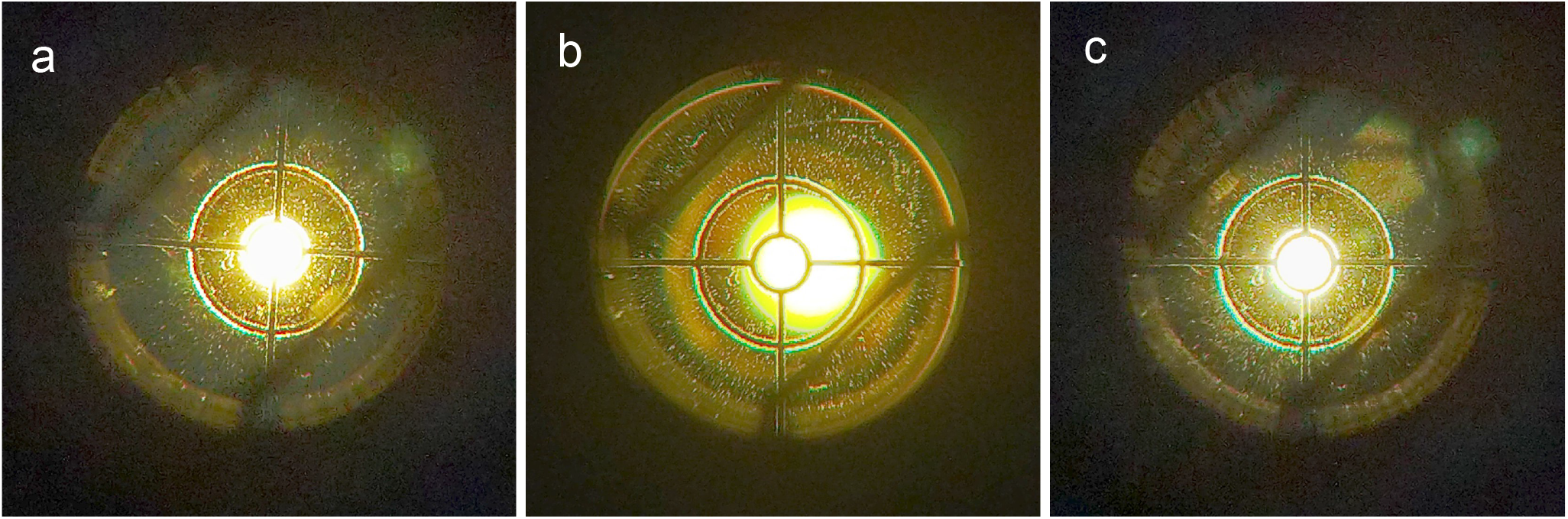
View into the empty ocular tube while aligning the objective holder: (a) Iris image (both irises closed) centred on crosshair, crosshair of the aligment disc not centred onto the microscope beampath - correct with Köhler screws; (b) Iris image (only 1 iris closed here) not centred, crosshair centred on microscope beampath - correct with alignment screws on objective holder; (c) Iris image, crosshair and microscope beampath centred. Images were taken with a Cat S62 Pro mobile phone (Bullitt Mobile, Reading, UK) using an Omegon Easypic Universal smartphone Adapter (nimax, Landsberg, Germany) on the ocular tube of an Zeiss AxioImager Z.1 with the oculars removed.

### Construction of the stage adaptor and sample holder

A construction manual for a simple stage adaptor with manual rotation can be found in the supplementary pdf “ftConstruction_manualStage”. All parts are included in the “Profi Optics” Set offered by Fischertechnik.

For a detailed construction plan of the sample holder with motorised rotation please see supplementary pdf “ftConstruction_motorisedStage”. The document also contains a Fischertechnik parts list.

All construction plans for Fischertechnik can easily be modified and adapted to different stages, incubator housings and experimental requirements due to the highly modular design.

### Mounting of the stage adaptor and sample holder

Click the stage adaptor into the multiwell plate holder of your microscope stage like a multiwell plate. The side where the sample holder gets attached should face in the same direction as the objective on the objective holder. Move the stage far out in that direction without covering the space above the empty objective position of the of the objective turret (should still be and remain in the beam path). After mounting and centering the sample on the sample holder slide it into its designated position on the stage adaptor and focus onto the sample using the x-drive of your microscope stage. For moving in y-direction you use the y-drive of the stage, for moving the field of view vertically (= x-direction at the oculars/ on the camera image) use the condenser focus of your microscope stand.

### Sample preparation

In the following, we will give a description for preparing young A. thaliana seedlings. For other types of samples this protocol needs to be adapted accordingly.

1. Prepare the sample storage container
  a. Get a closed container for storing the slides upright, e.g. a staining container from histology supplies does a good job (Fig. 4c).
  b. Place a wet paper inside the container to keep up a humid environment.
2. Prepare the slide
  a. Prepare a slide with double-sided adhesive tape (Tesa, type 05338, Beiersdorf, Hamburg, Germany) to build a buffer reservoir (green polygon in Fig. 4b).
  b. Add growth medium or buffer to buffer zone 1 (purple box in Fig. 4b).
  c. Put a seedling on the slide (as strait as possible).
  d. Add the coverslip (red square in Fig. 4b) that only the root is covered (so the seedling doesn’t get injured.
  e. Build a bigger buffer reservoir with medium or buffer containing LM agarose (Agarose, Low Melting Point, Promega Corporation V2111) in zone 2 (blue box in Fig. 4b) to keep the seedling alive during microscopy.
    i. Wait till the agarose has 37°C before you add it to the slide.
    ii. This is to keep the sample humid, leaves use up lots of water during the scans.
    iii. ∼60 min imaging can be done, during longer sessions the sample may get dry.
3. Slide storage
  a. Put in the slides in the prepared storage container as soon as the LM agarose is solid and immediately close the box.
  b. Leave the slides like this until just before you start the imaging experiment.
4. Mounting the slide
  a. Always keep the slide in a way that the plant is vertically oriented.
  b. Fix the slide with the sample located in the centre of the holder (Fig. 4d).
  c. If the part of the plant to be imaged is correctly centred you don’t need much correction during the experiment.
    i. Fixing a previously calibrated crosshair behind the sample holder facilitates orienting the sample for imaging (Fig. 4e).

A description for building an alternative sample chamber suitable for observations of thicker specimen can be found in the supplementary pdf “chamberPrepVerticalImaging”.

**Figure 4:**
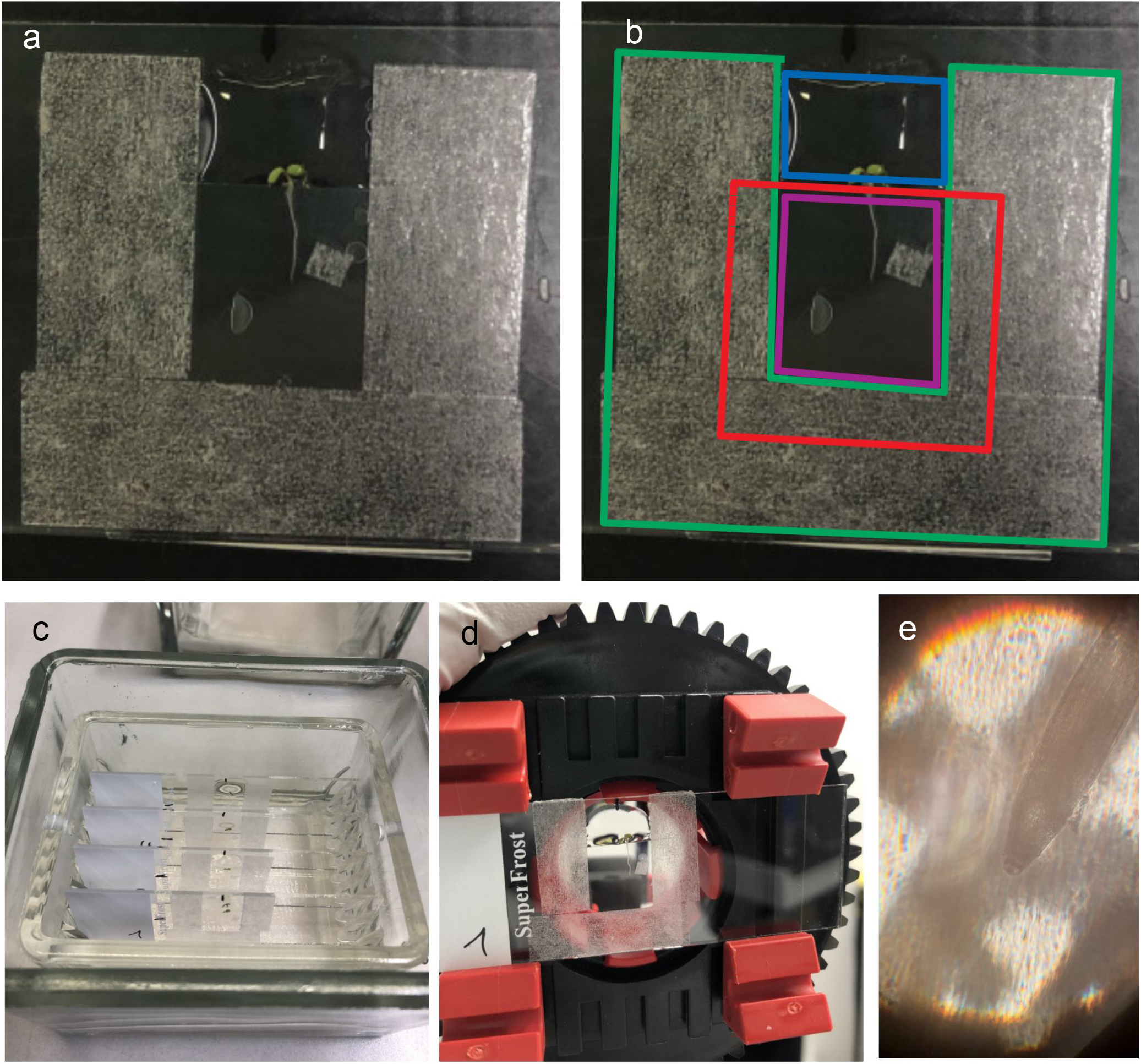
Example of sample preparation for young A. thaliana seedlings; (a) and (b) sample chamber on a slide (for coloured boxes in b) please see text); (c) Sample storage; (d) Mounted slide on rotation stage; (e) Root tip of A. thaliana seedling with the crosshair for sample orientation in the background.

### Measurement of optical quality

Optical quality measurements were done on an Olympus IX-83 (Olympus, Hamburg, Germany) equipped with a FLIR Flea 3 camera (Point Grey/ FLIR FL3-U3-13S2M-CS, FLIR Systems, Wilsonville, USA, mounted on a 1x camera adaptor) or with a Basler Ace camera (Basler Ace, acA2440-75uc, Basler, Ahrensburg, Germany, mounted on a 0.63x camera adaptor) and controlled by Micro-Manager 1.4 (Edelstein et al. 2014). We also used the IX-83 with an attached FluoView 1200 (FV 1200) scanhead to test the performance for confocal microscopy.

Additional PSF measurements were done on a Zeiss AxioObserver with Colibri 7 illumination unit and an AxioCam 702 running on ZEN blue 2.6 (Zeiss, Oberkochen, Germany). For the PSF measurements on the IX-83 the Flea 3 camera was used with a 20x/0.75 Plan Apochromat objective (Olympus, Hamburg, Germany, UPlanSApo 20x). For the PSF measurements on the AxioObserver we used a 20x/0.8 Plan Apochromat objective (Zeiss, Oberkochen, Germany, P/N: 440640-9903-000). VID_20210404_215528rev As sample we used subresolution beads derived from Zebra Mildliner Markers (orange marker from WKT7-5C, Zebra Co., Tokyo, Japan). We distinguished between the edge of the field of view (FOV) and the centre of the field of view. The centre was defined as the central square of the image with a side length of 1/3 of the image width. Edge measurements were done in squares of the same size positioned in the corners of the image.

For PSF visualisation (see Fig. 5) the 40/0.9 UPlanSApo objective and orange PS Speck beads (Thermo Fisher Scientific, Waltham US) were used on the IX-83 wit the Flea 3 camera..

**Figure 5.**
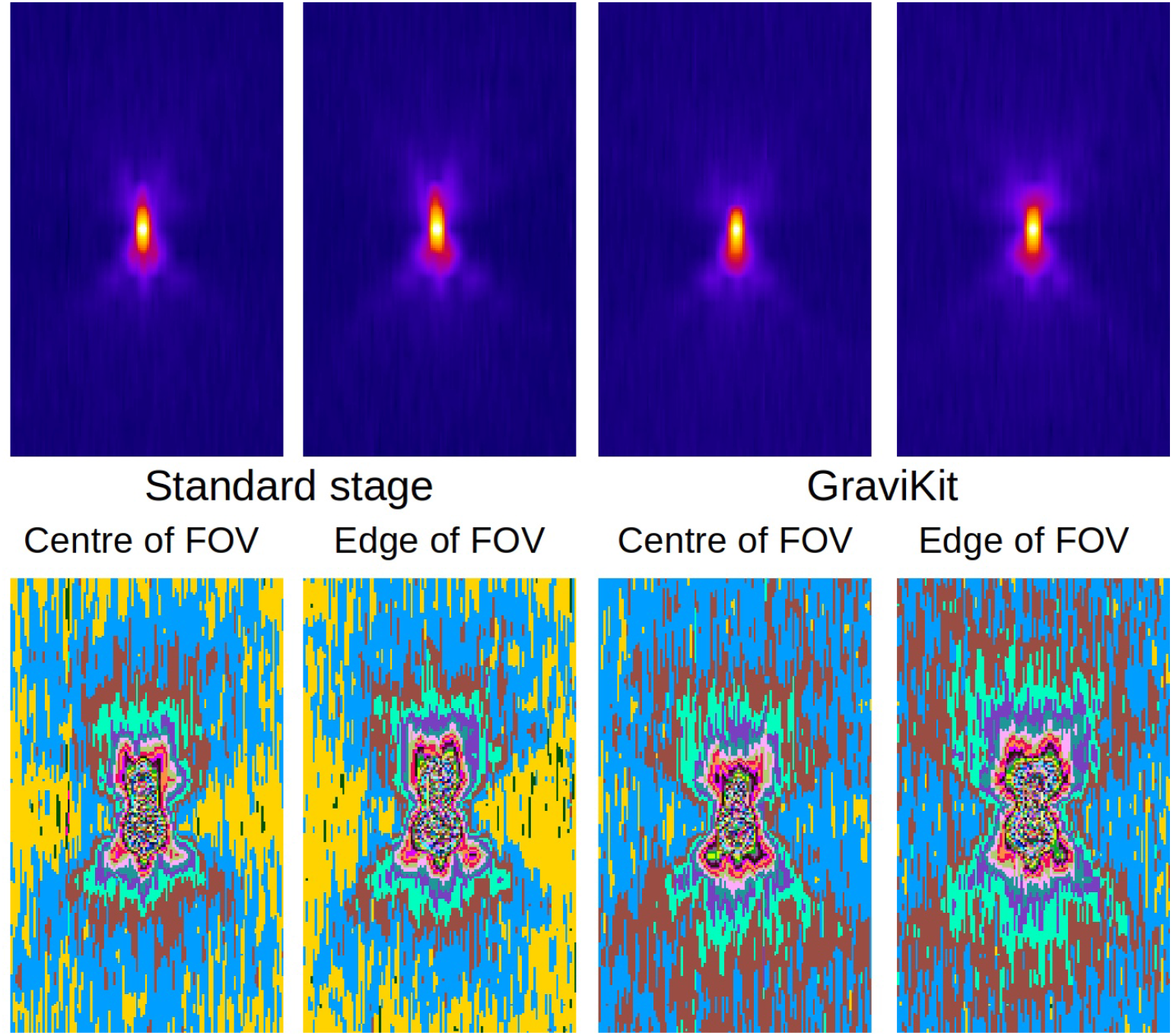
shows an example PSF taken with a 40/0.9 objective. The PSF shape with the sample mounted on a standard stage and on the GraviKit is visualised with ImageJ’s “Fire” LUT (upper panel) and “Glasbey” LUT (lower panel), both in the centre and on the edge of the field of View. Image width of all images is 10µm.

For the field homogeneity measurements on the IX-83 we used both cameras representing 2 different sizes of FOV and the FV 1200 scanhead with a 20x/0.75 Plan Apochromat objective. We used a yellow Chroma slide as sample, focussing on the brightest plane. As a measure for field homogeneity we used the coefficient of variation over a line profile along the diagonal of the FOV (Zucker 2006).

### Estimation of horizontal beampath usage in recent plant literature

Databases of 5 plant journals have been searched with the term “fluorescence microscopy roots”. The 10 most recent of those articles which utilised microscopic imaging of intact living roots were assessed whether they used a vertical beampath (standard upright or inverted microscope) or a horizontal beampath (e.g. light sheet, tilted standard stand).

## Results

### Optical quality

As the GraviKit elongates the optical path a reduced optical performance might occur. While the elongation is taking place in infinity space, in theory no such loss should occur. However, in commercial microscope systems the diameter of the tube lens is designed for a certain distance from the objective for a given microscope setup. Further, the interplay between objective and tube lens (e.g. for aberration correction) might also be compromised. Therefore field homogeneity and point spread function (PSF) were measured to assess the imaging quality using the conversion kit.

From the PSF measurements we achieved the full width half maximum (FWHM) values shown in Table 3. Some example PSFs are shown in Fig. 5, taken with a higher NA to better highlight changes in the PSF shape when using the GraviKit.

**Table 3.**
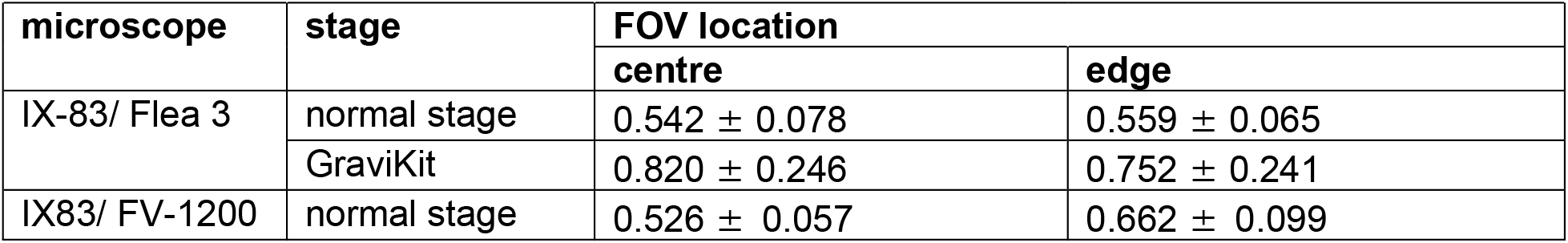
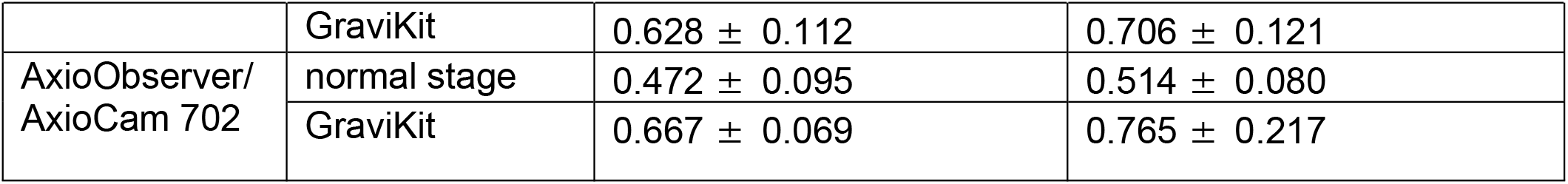
shows the FWHM measurements in µm for subresolution beads on different setups. The IX-83 was equipped with a 20x/0.75 objective, the AxioObserver was equipped with a 20x/0.8 objective. Measurements were done in the centre and on the edges of the field of view.

| microscope | stage | FOV location |  |
| --- | --- | --- | --- |
|  |  | centre | edge |
| IX-83/ Flea 3 | normal stage | $0.542 \pm 0.078$ | $0.559 \pm 0.065$ |
| | GraviKit | $0.820 \pm 0.246$ | $0.752 \pm 0.241$ |
| IX83/ FV-1200 | normal stage | $0.526 \pm 0.057$ | $0.662 \pm 0.099$ |
| | GraviKit | $0.628 \pm 0.112$ | $0.706 \pm 0.121$ |
| AxioObserver/<br>AxioCam 702 | normal stage | $0.472 \pm 0.095$ | $0.514 \pm 0.080$ |
| | GraviKit | $0.667 \pm 0.069$ | $0.765 \pm 0.217$ |

For the field homogeneity we got the results shown in Table 4 and Figure 6.

**Table 4.** shows the coefficient of variation for diagonal profile plots over different field of view (FOV) sizes as a measure for field homogeneity.

| FOV diameter (detector) | normal stage | GraviKit |
| --- | --- | --- |
| $583\mu\text{m}$ (Basler Ace) | 0.131 | 0.389 |
| $307\mu\text{m}$ (FLIR Flea3) | 0.058 | 0.168 |
| $899\mu\text{m}$ (Olympus FV1200) | 0.054 | 0.599 |

**Figure 6.**
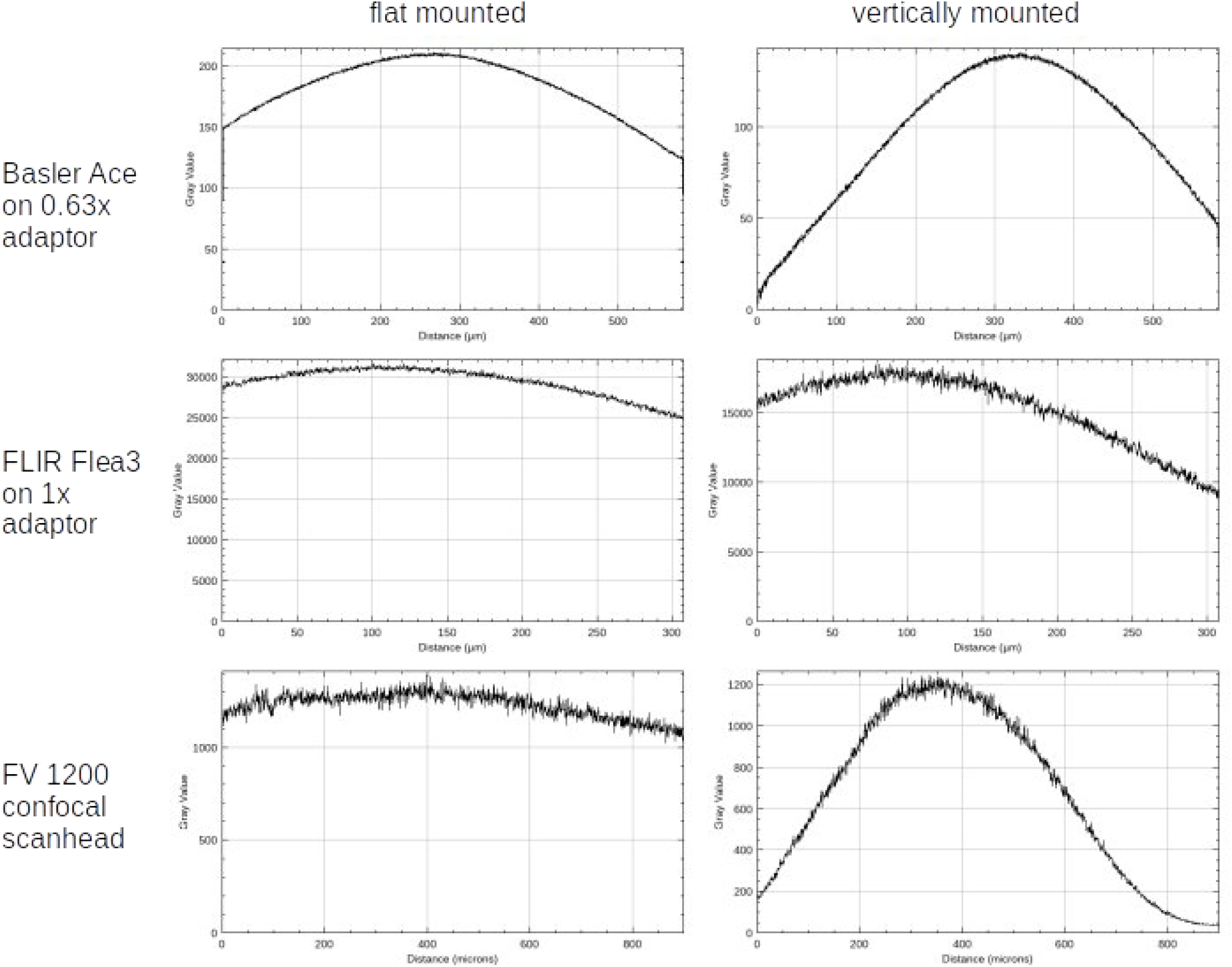
shows line profiles along the diagonal of the field of view on different setups. Left column shows the line profiles on a normal setup, right column shows the profiles using the GraviKit.

### Precision of positioning

For use in scientific experiments, especially with potential time constraints, a precise positioning of the rotating stage is essential. Therefore we tested the repositioning accuracy of the GraviKit. All measurements were done with the motorised setup to rule out bias by the manual rotation accuracy (please see a movie of the stage with motorised rotation in the supplementary files: GraviKitRotates1 & 2).

We always did an initial rotation by 20° forward (or arbitrary number of degrees backward and then 20° forward) for settling the sample holder right after loading the sample before we take any image. Without this your first rotation may have an error of several degrees, making reliable positioning impossible.

With this as a prerequisite we tested under 3 different scenarios: forward rotation only (2x 180°, FF), simple forward and backward rotation (180° each, FB), forward and backward rotation with backlash compensation (+180° > -200° > +20°, FBc) (please see examples for all conditions in Fig. 7). Then the displacement was measured (see Table 5, 5 repeats with 2 full rotations for each condition, sample remounted after each repeat).

**Figure 7.**
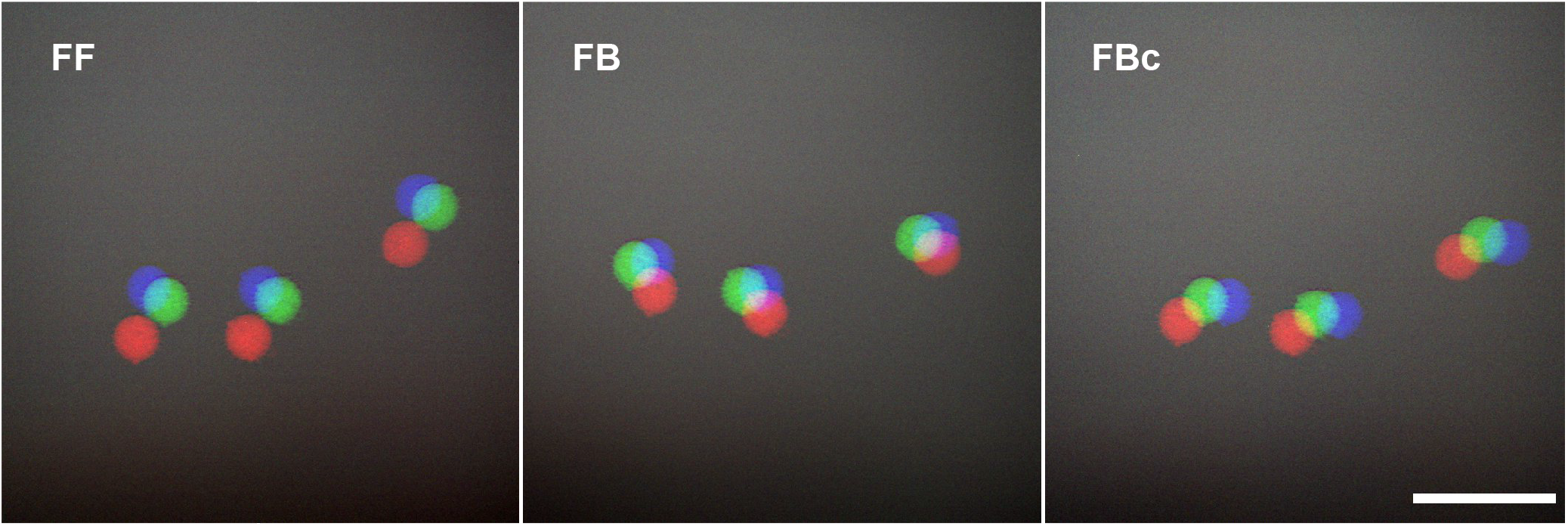
shows representative examples for the displacement by rotation for each of the conditions (FF, FB, FBc, see text). Start position in red, positions after 1 rotation (green) and 2 rotations (blue) are shown. Scale bar: 100µm.

**Table 5.** shows the position displacement after rotation with the GraviKit in µm.

| Condition | Average [μm] | STD [μm] |
| --- | --- | --- |
| FF | 16.6 | 14.0 |
| FB | 11.5 | 4.8 |
| FBc | 16.0 | 7.5 |

### Application of the GraviKit

In addition to the measured specifications we also have a range of application related experiences which are summed up below. Those experiences are derived from a study on gravitation dependent Auxin monitoring (Herud-Sikimic et al. 2021):

- After few rounds of practising the time for rotation and recentering can be reduced to less than one minute on a routine basis.
- There are no engineering skills needed for building and using the kit, the device is used on a routine basis by people with no background in engineering or (bio)-physics.
- The device is lightweight (objective holder: 463 g, stage adaptor/sample holder: 253 g (318 g incl. battery pack)) and has a small footprint, it can be easily fitted into an incubator box (see Fig. 8).
- Make sure the spectral range you’re imaging fits the specifications of the mirror built into the objective holder, when imaging outside of the specs of the specified mirror you need to fit in a suitable mirror.
- As there are manufacturing tolerances in multiwell plate holders the size of the stage adaptor may need to be adjusted by putting small stripes of tape on its corners to make it fit tightly into the multiwell plate holder.

**Figure 8:**
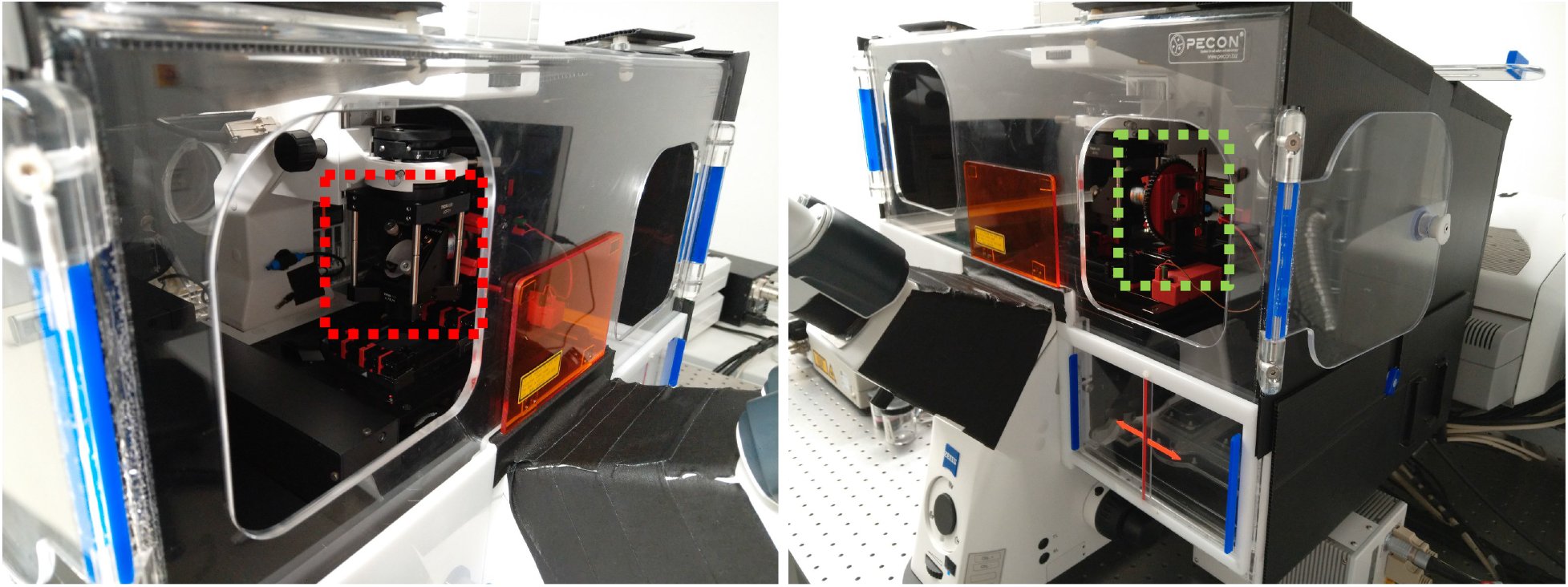
2 different views of the GraviKit mounted inside an incubator on a Zeiss AxioImager Z.1. Red dashed box: objective holder. Green dashed box: rotational stage.

## Discussion

Our aim was to create a system for observing vertically mounted specimen that is easy to implement in any laboratory. Therefore we decided to go for low price, off-the-shelf components and use as many features of existing inverted microscope stands as possible. The reason for using the condenser holder for mounting the objective was to be able to use the condenser focus as “x” drive. Using the native y-drive of the stage as y-drive and the x-drive of the stage for focussing we could use drives that were already on the microscope stand for all translational axes. This enabled us to reduce the cost and the footprint of the GraviKit substantially as commercially available drives are quite expensive, heavy and would take additional space.The only additional motorised drive used is the one for rotation (if the motorised version is desired). However, an axis for sample rotation is usually not available at all on standard microscope setups in combination with an xy-stage. So, even without motorisation, you gain an additional degree of freedom for sample movement.

As expected the optical resolution is negatively impacted which needs to be considered when using the GraviKit. The impact on optical resolution is slightly different on systems from different manufacturers, also concerning the location within the field of view. This may be due to different optical designs regarding the interplay between objective and tube lens. Moreover, the homogeneity of the field illumination is compromised. It should also be mentioned that alignment of the GraviKit gets more demanding with increasing NA. Therefore we recommend to use the GraviKit only with air objectives with an NA of up to 0.8 and objective magnification up to 40x. Using just a cropped field of view (e.g. for confocal with a setting like 2x zoom or higher) or a camera adaptor with a magnification higher than 1 will improve the experience. Mind that with these constraints it is still possible to image at subcellular resolution (Herud-Sikimic et al. 2021).

Rotation, even if manual repositioning is needed, was, with about 1 minute for any given angle, substantially faster than other published solutions for vertical sample mounting (von Wangenheim 2017, 3min). The repositioning accuracy was between 10µm and 20µm, which keeps the sample well within the field of view for the optical setups recommended above. Unexpectedly the backlash compensated rotation (FBc) turned out to be less precise than the non-compensated one (FB). This is probably due to the Ft stepper motor, as the step size of the motor is probably larger than the backlash induced error. Therefore the compensation (using 3 different numbers of steps for going forward - backward - compensate) might induce larger error than the option without compensation (using the same number of steps forward - backward).

The step size of the stepper motor is probably also responsible for the larger displacement when going 2 times in the forward direction (FF), as it doesn’t get to precisely 360° when going two times more or less precisely to 180°. The comparably large standard deviation can be explained by the fact that the strength of this effect depends on how accurately your sample was centred. With an overall cost for the manual version of less than 800 € for Thorlabs parts (e.g.753 € for IX 83 kit, inquired May 2021) plus ∼80 € for the Fischertechnik Optics Pro kit, our construction stays safely below the 1.000 € mark and is substantially cheaper than all other solutions around: for the tilted microscope solution (von Wangenheim 2017) costs can easily become a 6-digit number of Euros and a price inquiry at a company offering the periscope solution gave an estimate of around 10.000 €. For motorised rotation, one needs to purchase the Fischertechnik Robotics TXT Discovery kit in addition, which comes at another ∼400 €, so this variation considerably stays below our self-set 2.000 € limit.

In brief:

Pros:

– Fast rotation
– Low price
– Easy to implement
– Quick mounting and unmounting on a microscope, therefore suitable for multi-user environments
– Possibility to have 3 translational and 1 rotational axis in the same system

Cons:

– Only useful with air objectives with NA up to about 0.8
– Loss of resolution and signal intensity compared to standard mounted objectives
– (Currently) no option for elaborate transmitted light contrasting

## Supporting information

ftConstruction_motorisedStage (construction manual)

ftConstruction_manualStage (construction manual)

GraviKitRotates2 (movie)

chamberPrepVerticalImaging (sample preparation)

GraviKitRotates1 (movie)

LiteratureRootImaging (suppl. table)

## Acknowledgements

Marina Ortega-Perez & Max Kraner (provision of plants), Sara Mendes (help with IX-83 configuration), teams of OICE and Light Microscopy Facility at MPI for Developmental Biology (general assistance)

## Author Contributions

CF: original idea and construction of GraviKit, manuscript writing, tests on Olympus and Zeiss setups

MKo: sample preparation, tests and real-life application on Zeiss setup, technical feedback for CF and SH

MKu: design of calibration tool, tests on Nikon setup

SH: design, implementation and QC of motorised version of the stage, QC of manual version of the stage, tests on Olympus setup

RP: critical revision of GraviKit designs, sample preparation, tests on Nikon setup, manuscript writing

## References

Barnes CR (1896) A horizontal microscope. Botanical Gazette Vol. 22, No. 1, Jul 1896, p. 55–56; https://doi.org/10.1086/327378

Edelstein AD, Tsuchida MA, Amodaj N, Pinkard H, Vale RD, and Stuurman N (2014) Advanced methods of microscope control using μManager software. Journal of Biological Methods 2014 1(2):e11 doi:10.14440/jbm.2014.36

Hemmersbach R, von der Wiesche M, Seibt D (2006) Ground-based experimental platforms in gravitational biology and human physiology. Signal Transduction, 6:381–387. doi:10.1002/sita.200600105.

Herranz R, Medina FJ (2013) Cell proliferation and plant development under novel altered gravity environments. Plant Biol J, 16: 23–30. https://doi.org/10.1111/plb.12103

Herud-Sikimic O, Stiel AC, Kolb M, Shanmugaratnam S, Berendzen KW, Feldhaus C, Höcker B, Jürgens G (2021) A biosensor for the direct visualisation of Auxin. Nature 592(7856): 768–772. doi: 10.1038/s41586-021-03425-2

Maizel A, von Wangenheim D, Federici F, Haseloff J, Stelzer EH (2011) High-resolution live imaging of plant growth in near physiological bright conditions using light sheet fluorescence microscopy. The Plant Journal 68: 377–385. doi: 10.1111/j.1365-313X.2011.04692.x, PMID: 21711399

Monshausen GB, Miller ND, Murphy AS, Gilroy S (2011) Dynamics of auxin-dependent Ca2+ and pH signaling in root growth revealed by integrating high-resolution imaging with automated computer vision-based analysis. The Plant Journal 65:309–318. doi: 10.1111/j.1365-313X.2010.04423.x, PMID: 21223394

Ovecka M, von Wangenheim D, Tomancak P, Samajova O, Komis G, Samaj J (2018) Multiscale imaging of plant development by light sheet fluorescence microscopy. Nature Plants 4, 639–650 (2018). https://doi.org/10.1038/s41477-018-0238-2

Takahashi K, Takahashi H, Toyota M, Furutani-Seiki M, Kobayashi T, Watanabe-Takano H, Shinohara M, Numaga-Tomita T, Sakaue-Sawano A, Miyawaki A, Naruse K (2021) Gravity sensing in plant and animal cells. npj Microgravity 7, 2. https://doi.org/10.1038/s41526-020-00130-8

Volkmann D, Baluska F (2006) Gravity: one of the driving forces for evolution. Protoplasma 229, 143–148. https://doi.org/10.1007/s00709-006-0200-4

von Wangenheim D, Hauschild R, Fendrych M, Barone V, Benková E, Friml J (2017) Live tracking of moving samples in confocal microscopy for vertically grown roots. eLife 2017; 6:e26792 doi: 10.7554/eLife.26792

Zucker RM (2006), Quality assessment of confocal microscopy slide based systems: Performance. Cytometry, 69A: 659–676. https://doi.org/10.1002/cyto.a.20314

