## Supplementary material for "GraviKit: an easy-to-implement microscope add-on for observation of gravitation dependent processes": ftConstruction_motorisedStage (construction manual)

### GraviMicKit
### -motorised version-
### *Fischertechnik constructional manual*

### Pictures from different angles

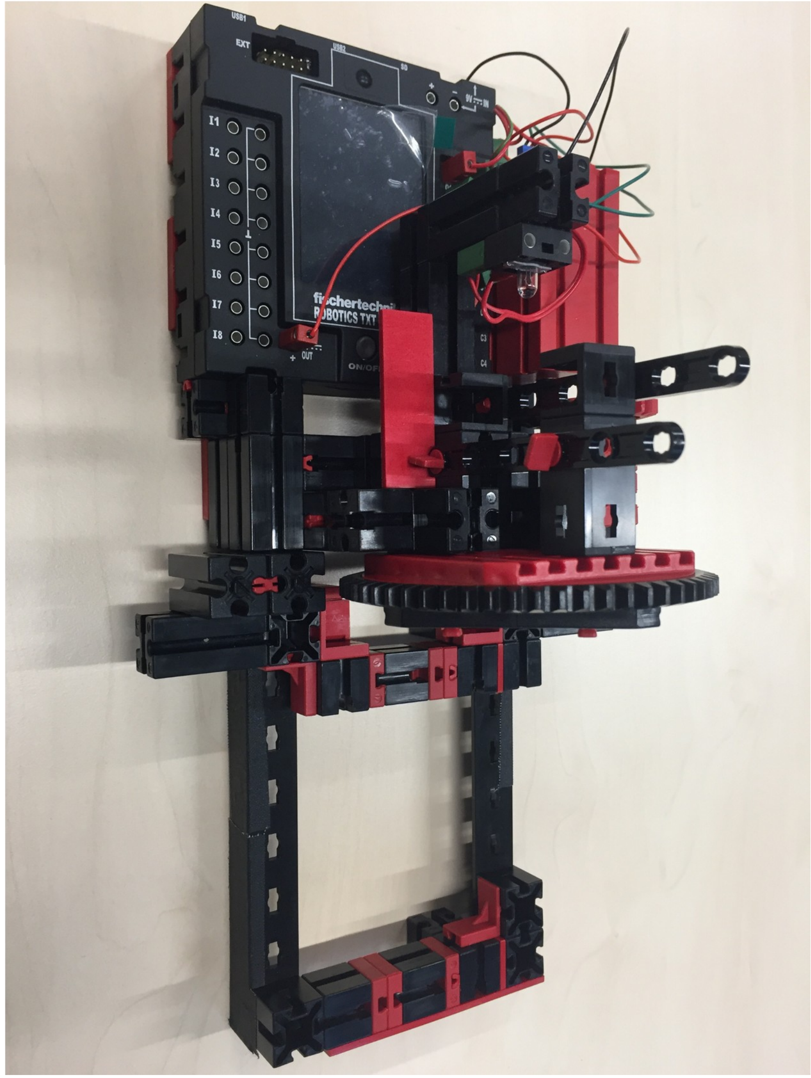

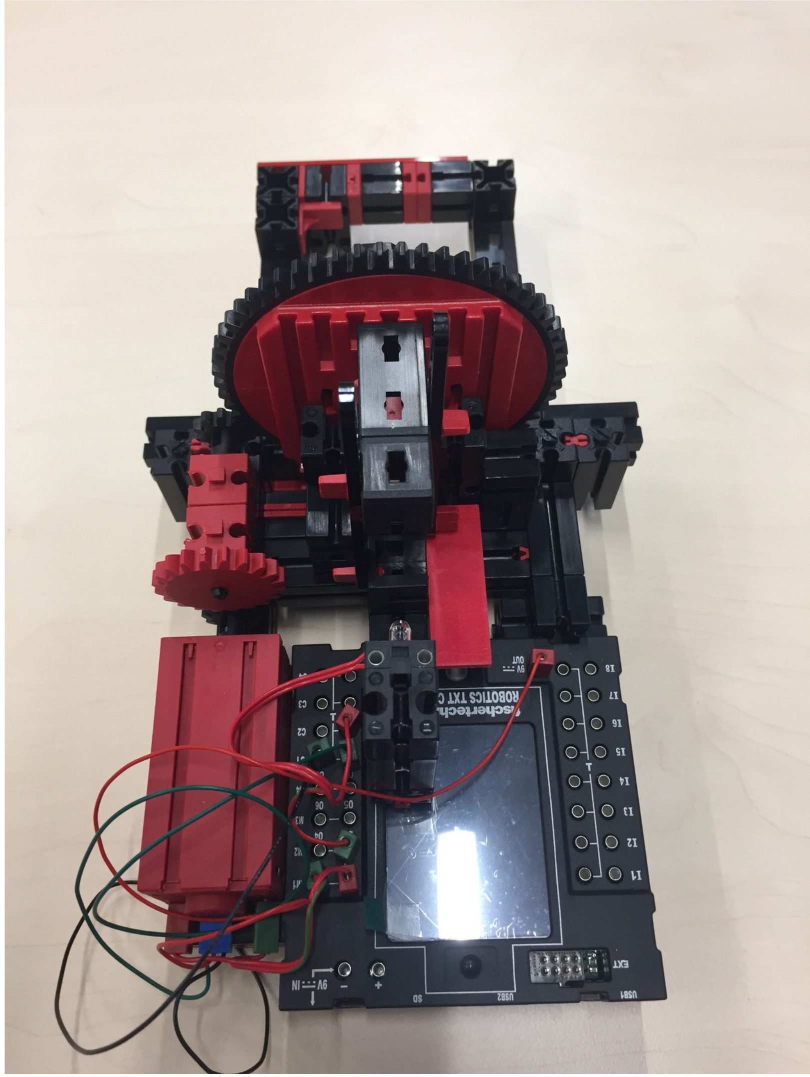

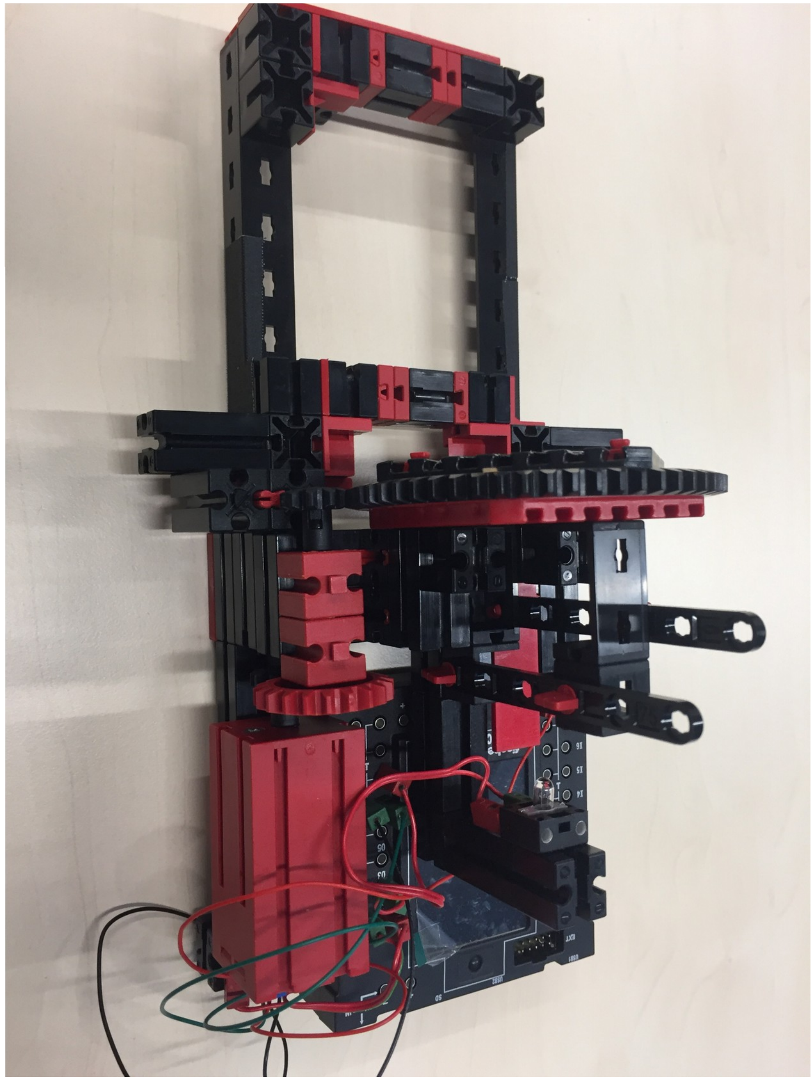

### Pictures from different angles

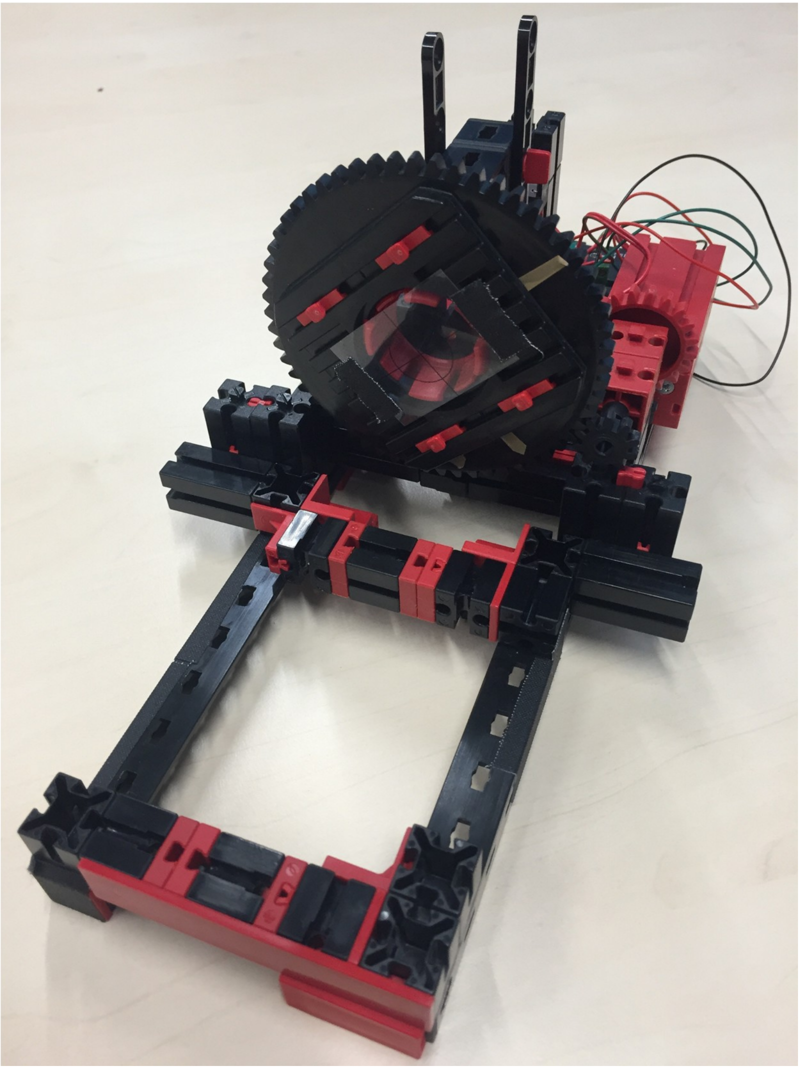

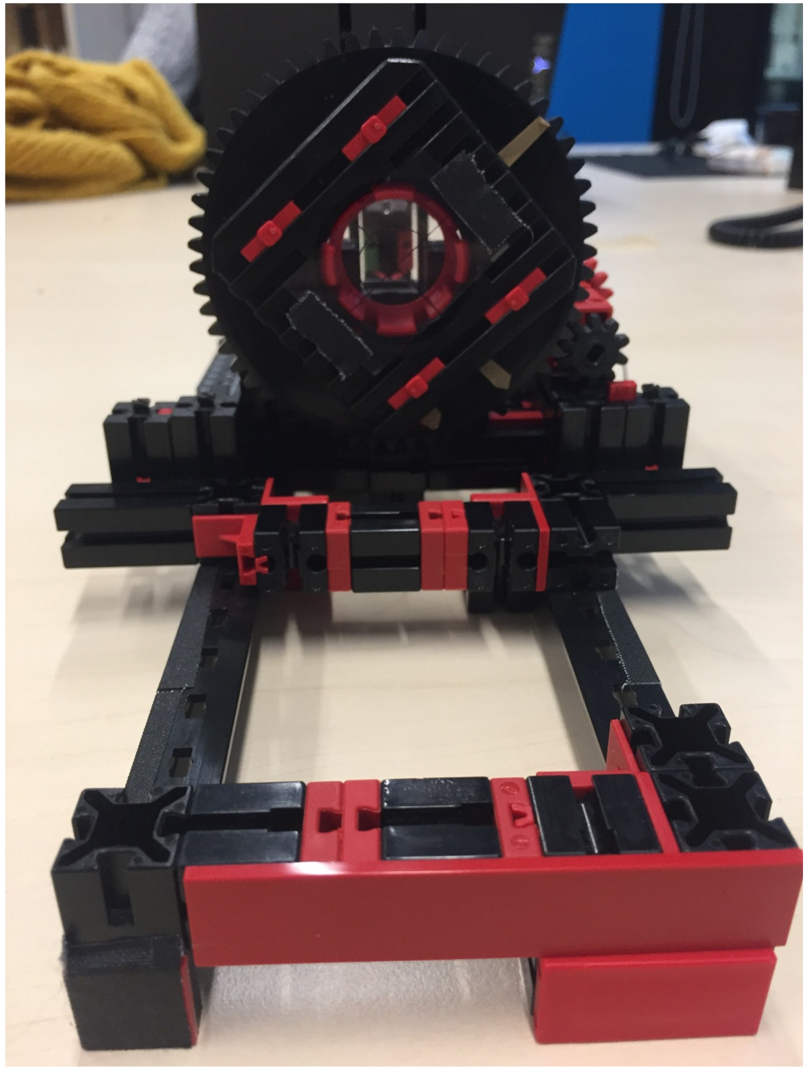

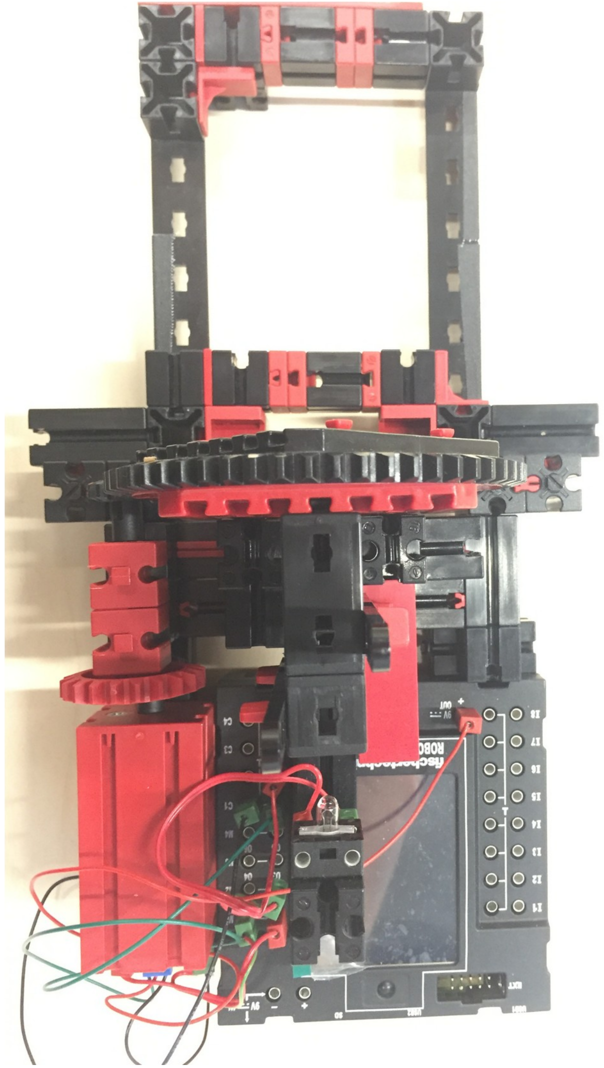

### Pictures from different angles

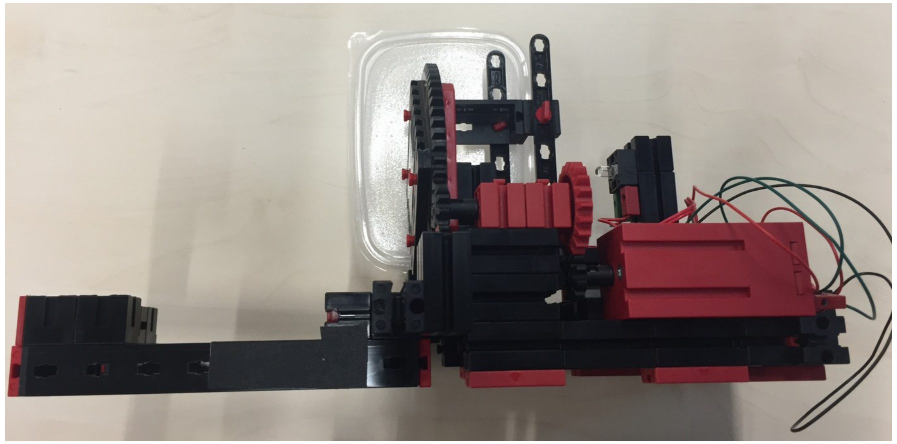

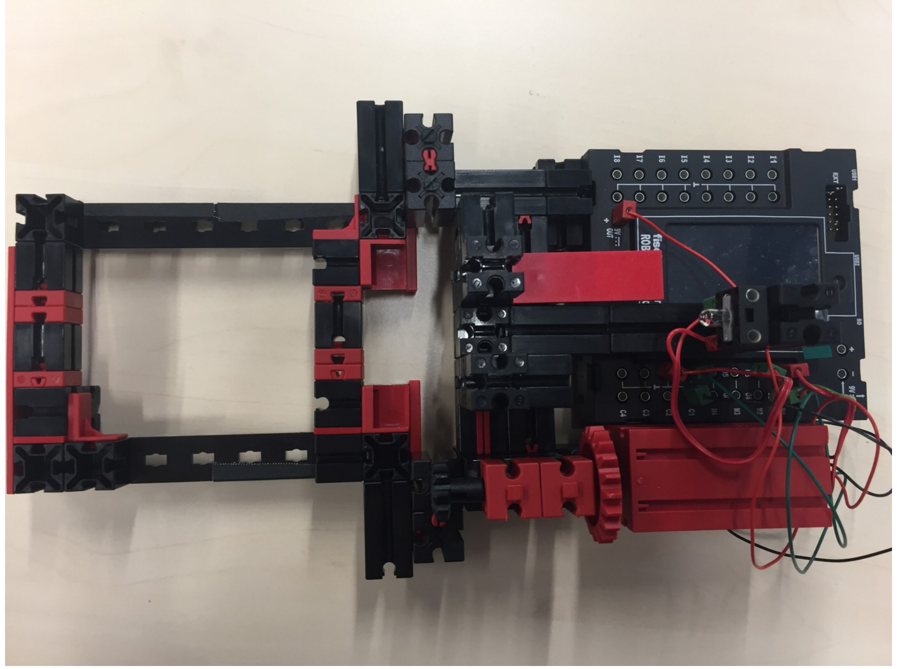

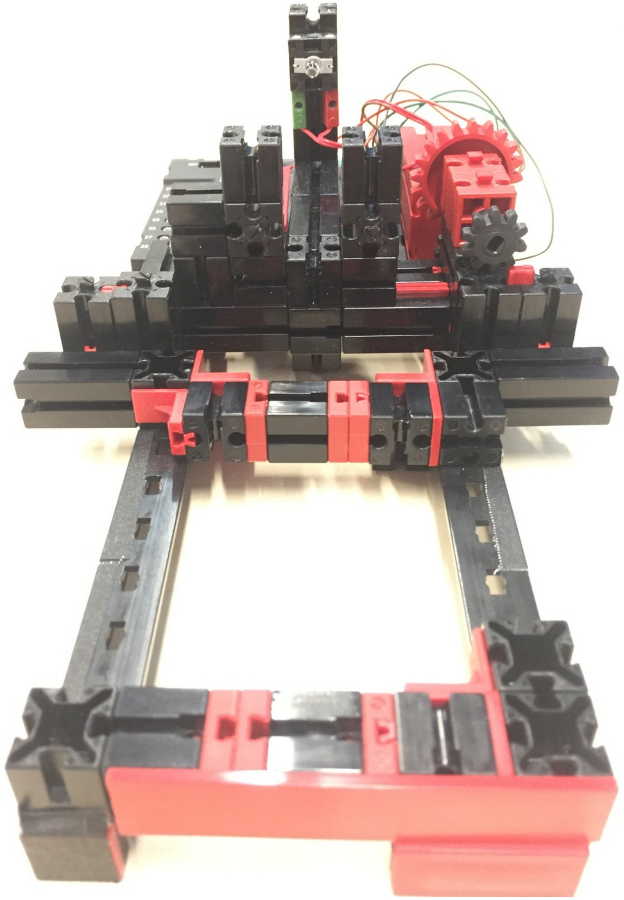

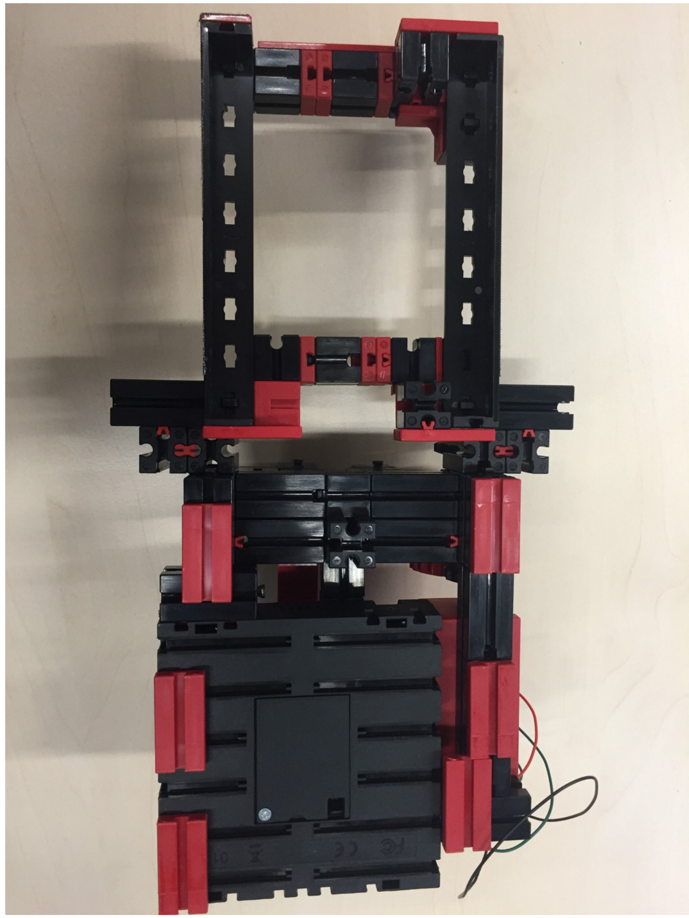

### All Fischertechnik parts used

| Baustein | Art. number | Anzahl |
|---|---|---|
| Baustein 30 | 32879 | 23 |
| Baustein 15 | 32881 | 27 |
| Verbindungsstück 30, rot | 31061 | 8 |
| Baustein 5, rot | 37237 | 6 |
| Baustein V15 Eck, rot | 38240 | 2 |
| Winkelstein 10x15x15, rot | 38423 | 2 |
| Bauplatte 15x30x3,75 mit Nut, rot | 32330 | 3 |
| Baustein 15x30x5 mit Nut und Zapfen, rot | 35049 | 5 |
| Federnocken, rot | 31982 | 22 |
| Riegelstein 15x15, schwarz | 32850 | 4 |
| Bauplatte 15x15, rot | 32850 | 3 |
| Bauplatte 15x45, rot | 38242 | 1 |
| Bauplatte 15x75, rot | 38244 | 1 |
| Baustein 15 mit Bohrung rot | 32064 | 2 |
| Pin, grün | 31336 | 6 |
| Pin, rot | 31337 | 6 |
| Kabel, grün/rot | 31360 | 1 |
| Winkelträger 120, schwarz | 36293 | 2 |
| I-Strebe mit Loch 75, schwarz | 36923 | 2 |
| Rastachse 45, schwarz | 35064 | 1 |
| S-Riegel 4, rot | 36323 | 4 |
| Winkelträger 15, schwarz | 36323 | 2 |
| Winkelträger 30, schwarz | 36920 | 2 |
| Rast-Ritzel Z10 m=1,5, schwarz | 35945 | 2 |
| Rastkettenrad Z20, rot | 137677 | 1 |
| Encodermotor, rot | 153422 | 1 |
| TXT Controller | 153513 | 1 |
| Drehkranz-Unterteil, rot | 31391 | 1 |
| Drehkranz-Oberteil, schwarz | 31390 | 1 |
| Power Set | 505283 | 1 |

### Construction manual

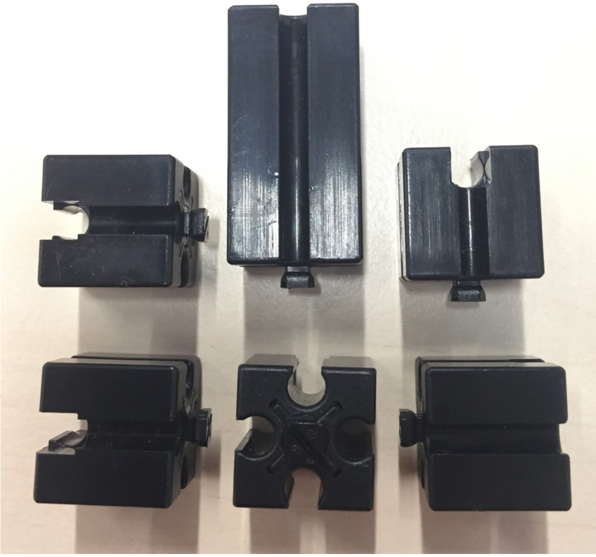

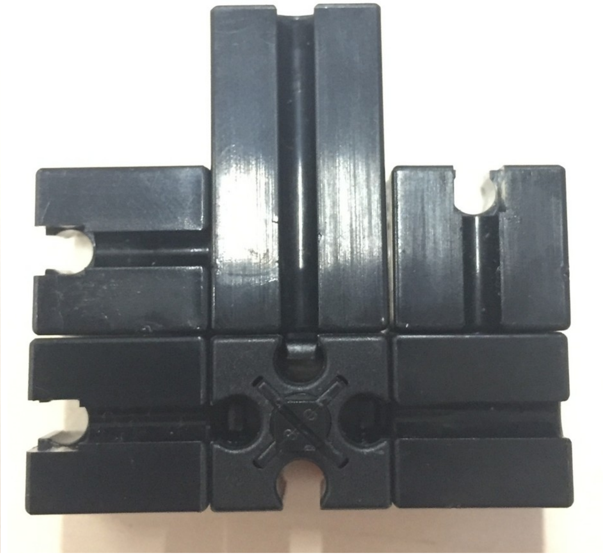

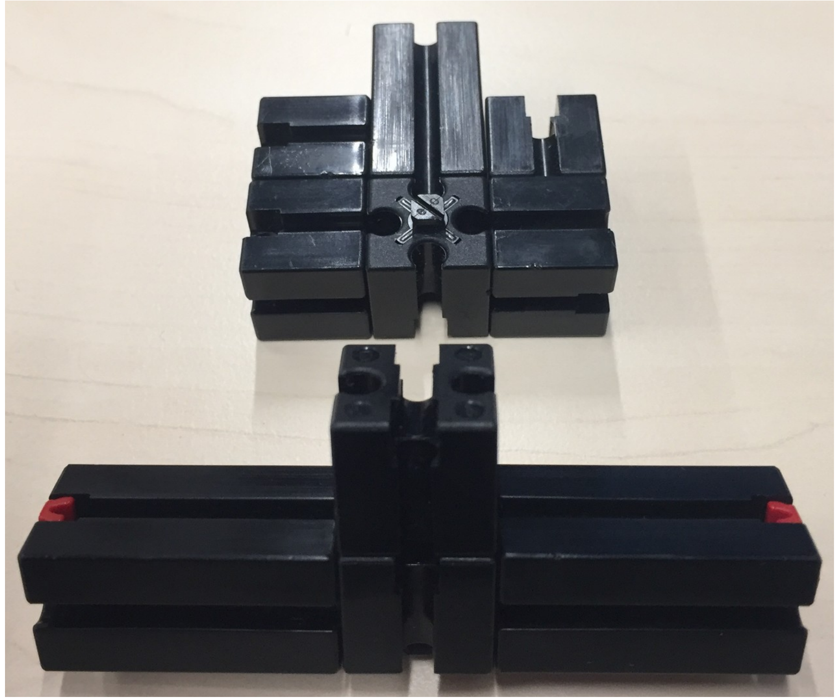

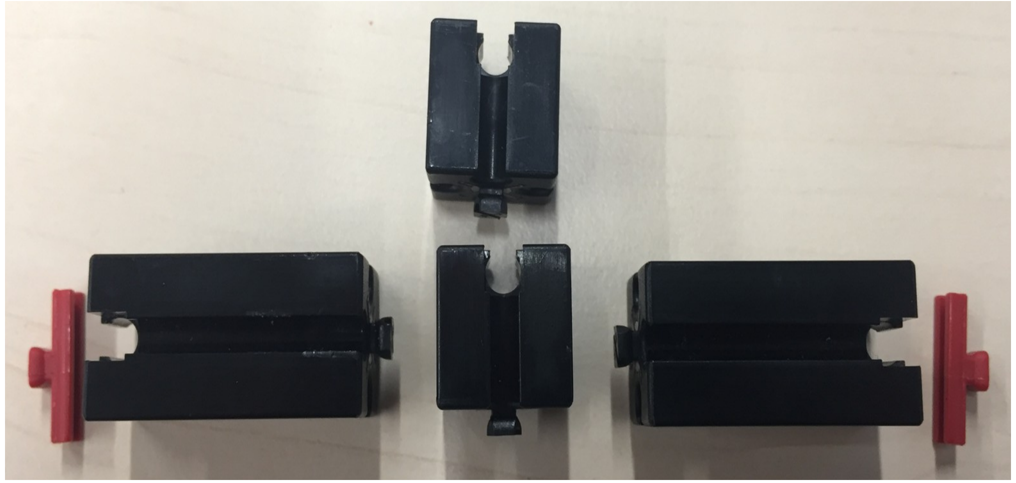

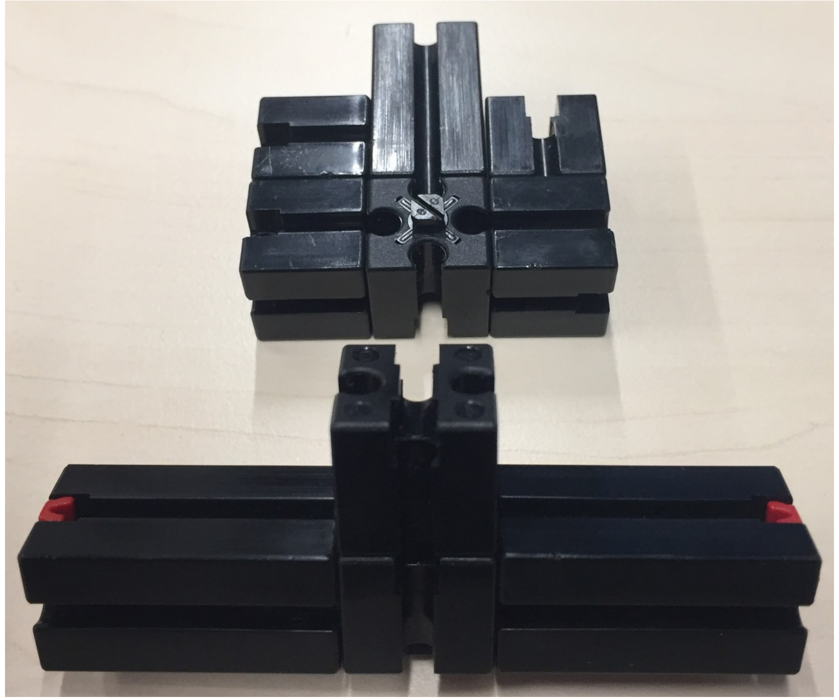

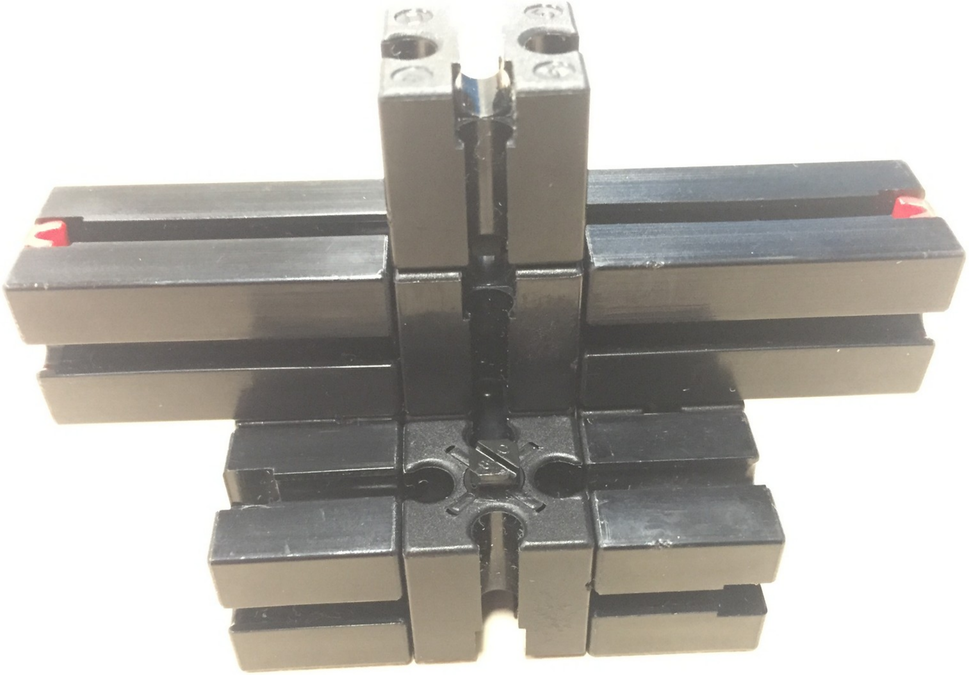

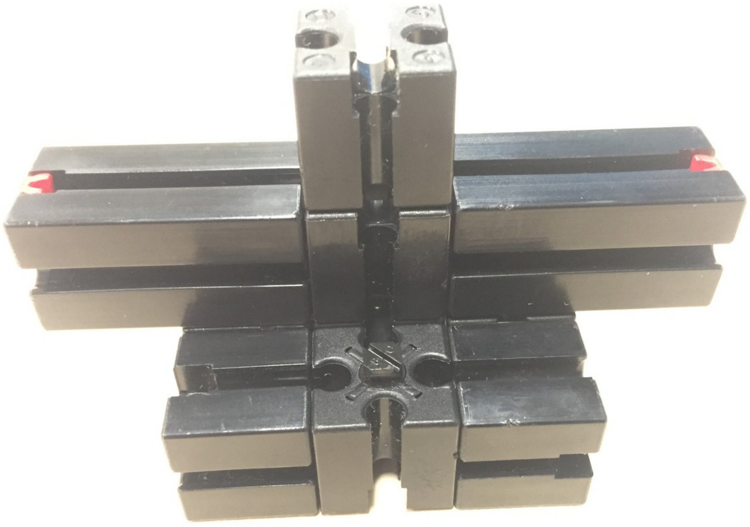
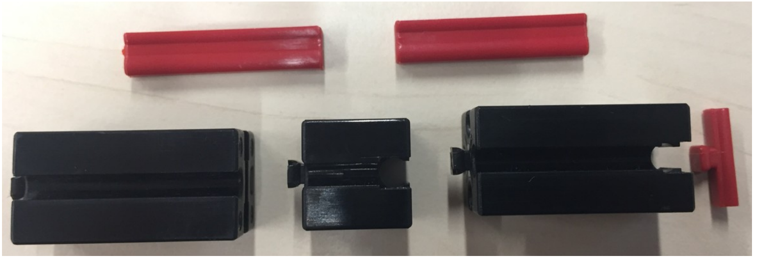
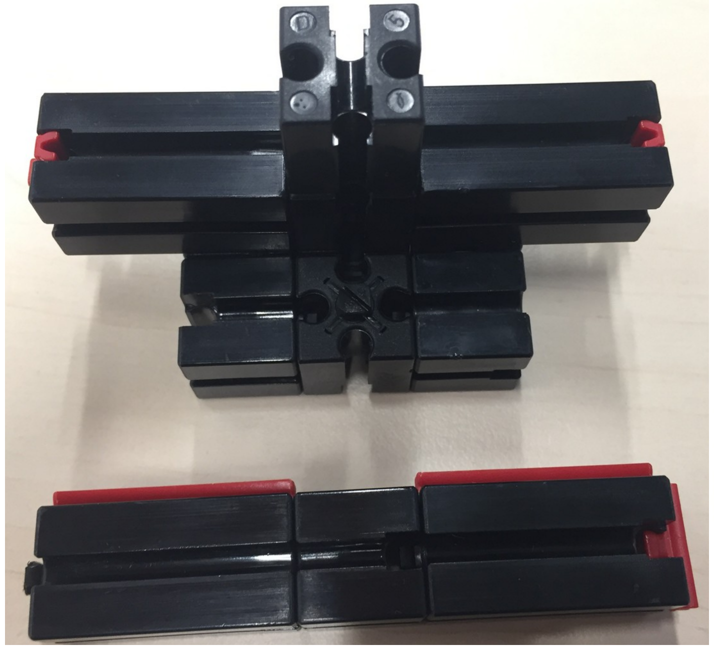
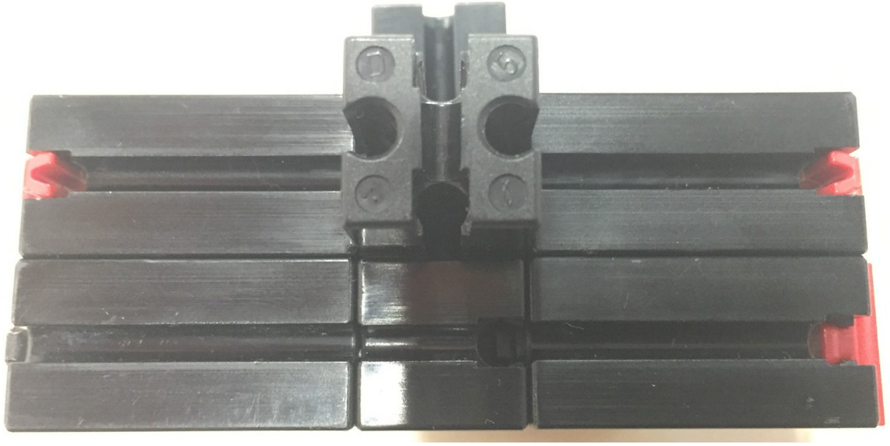

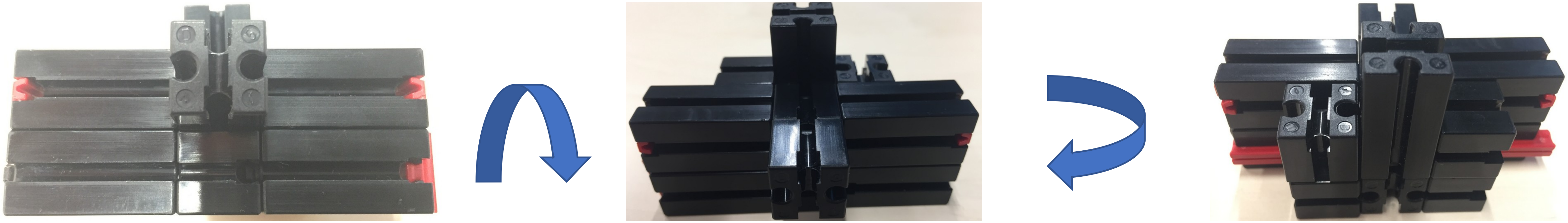

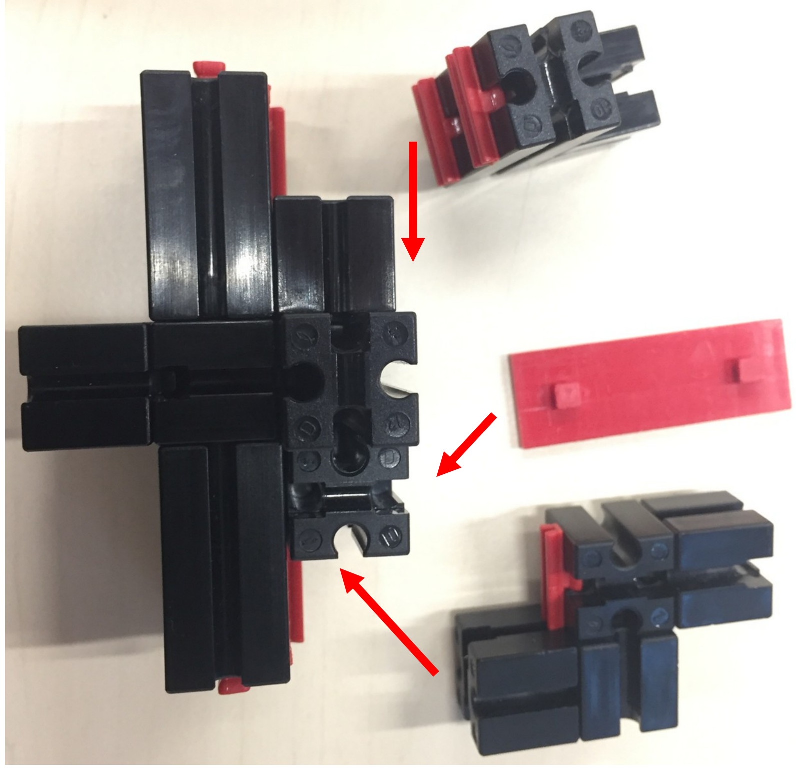
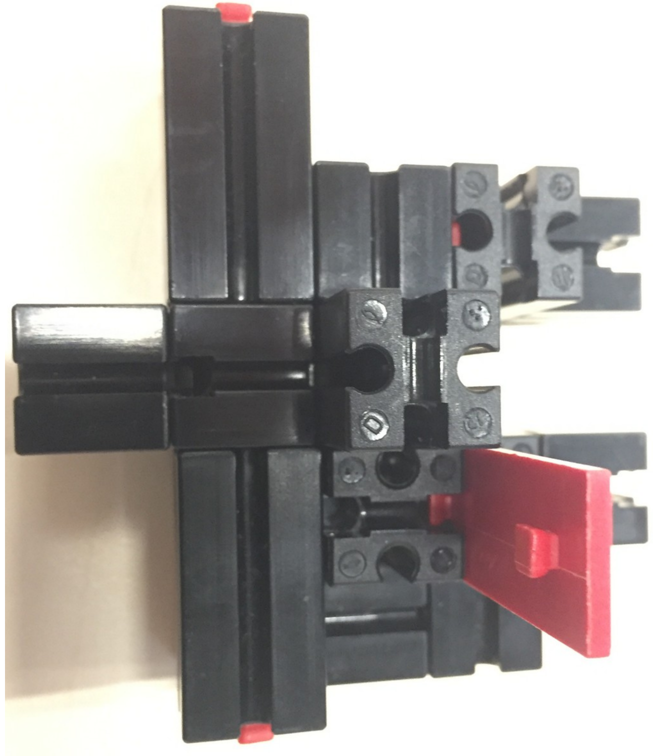

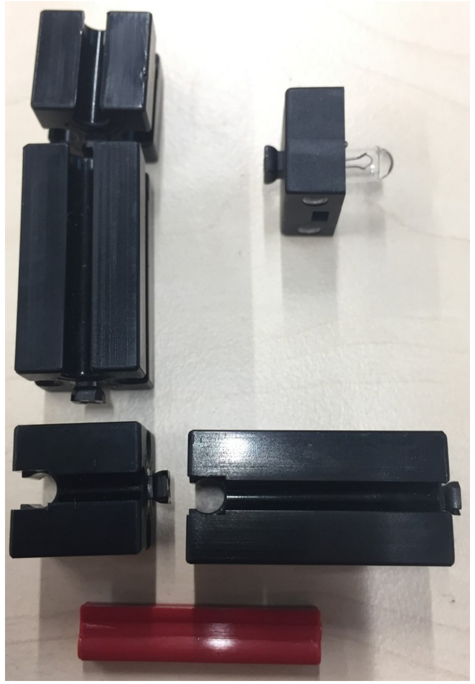

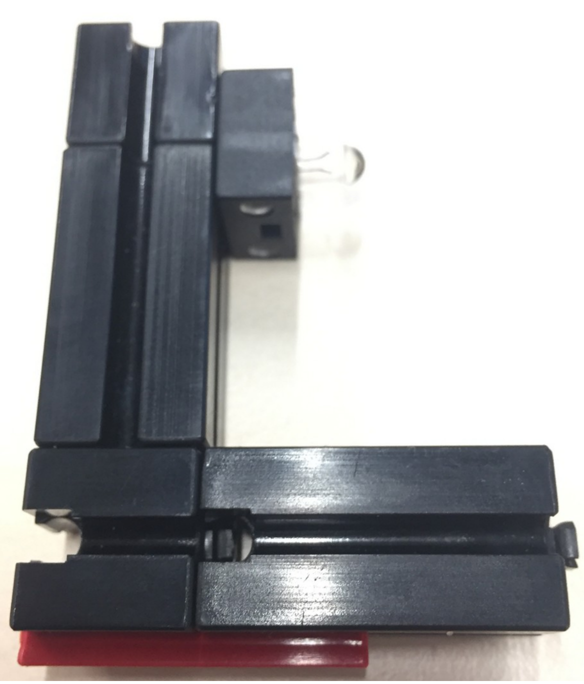

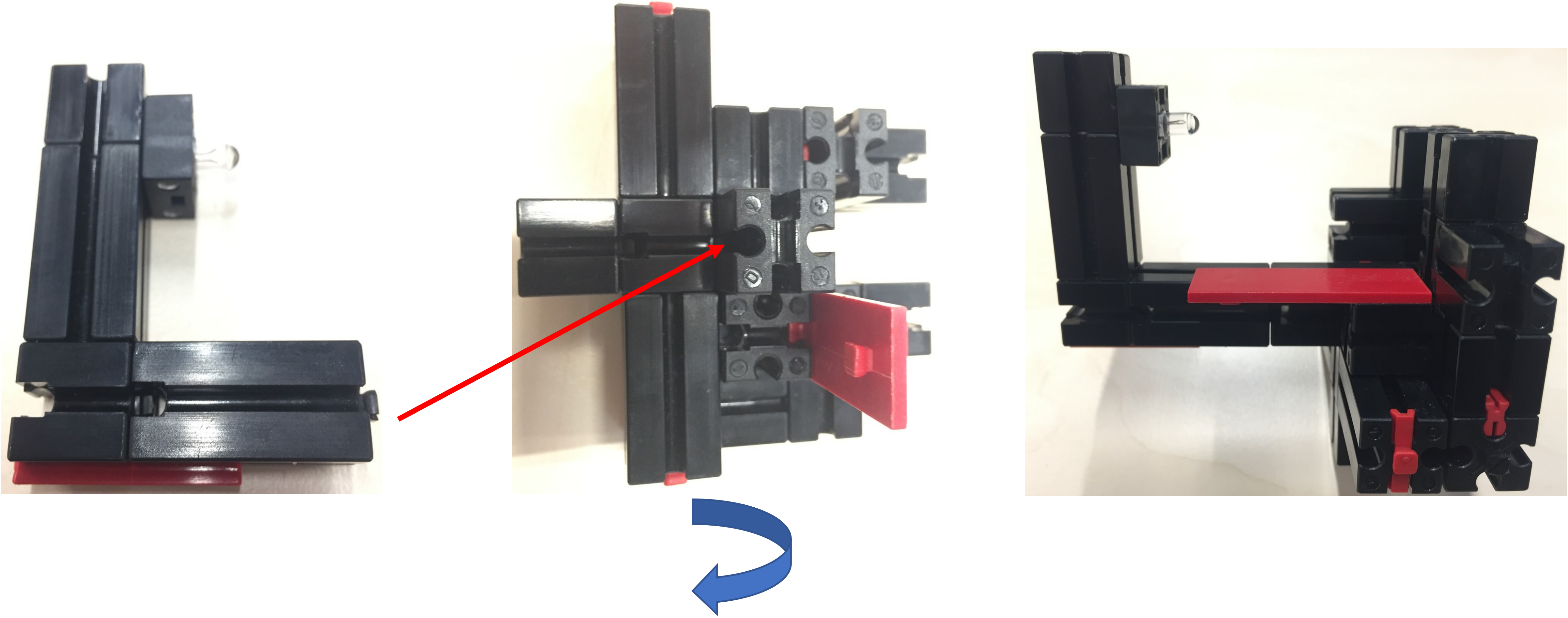

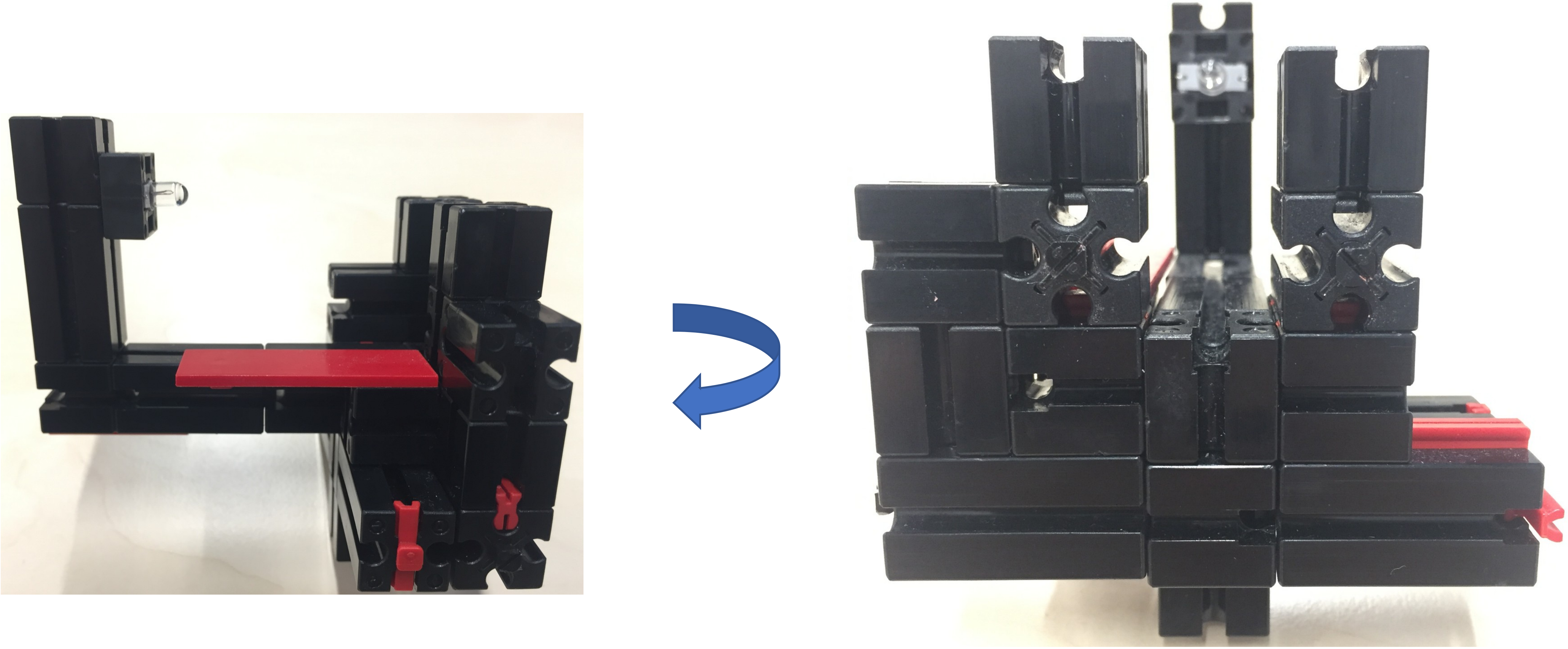

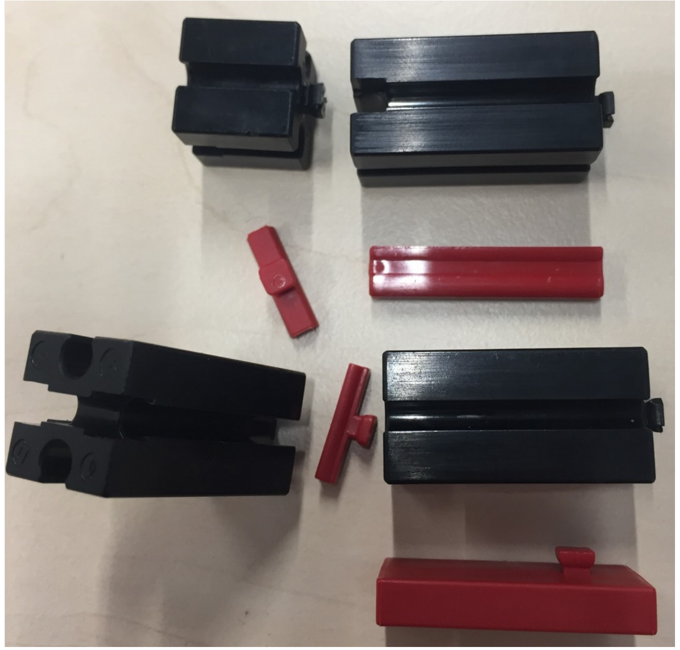

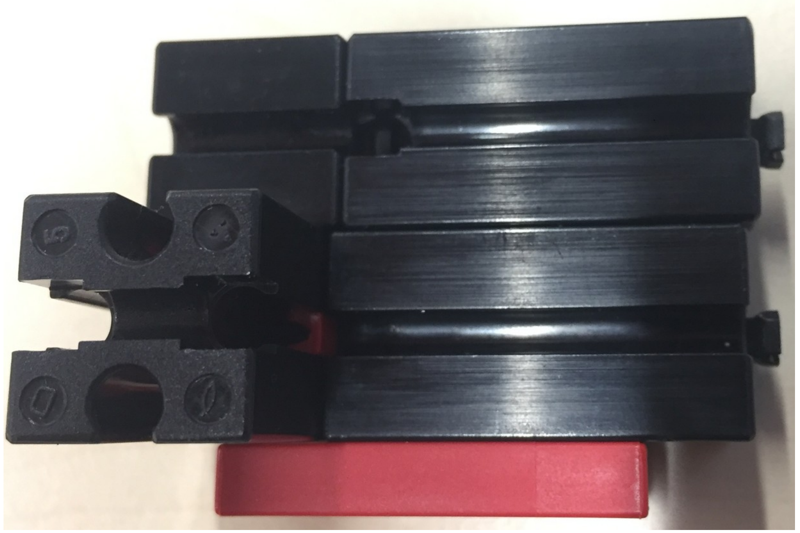

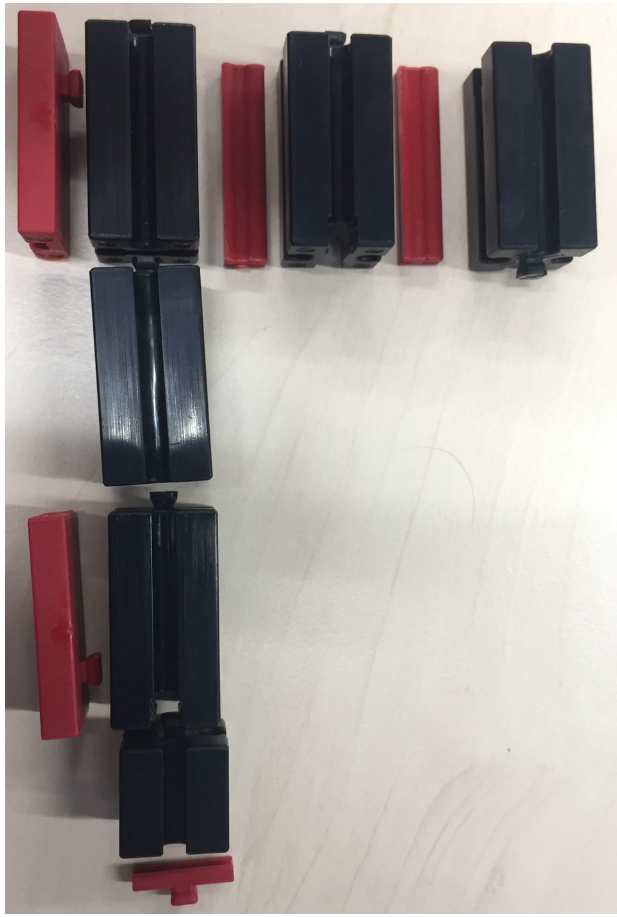
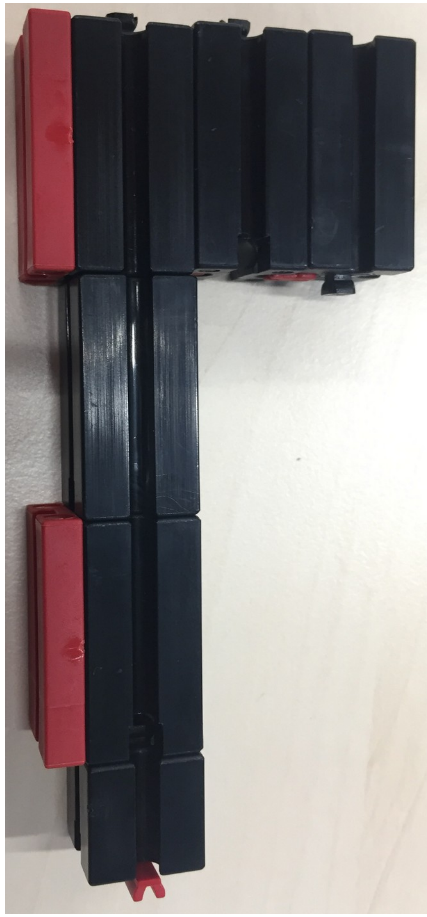

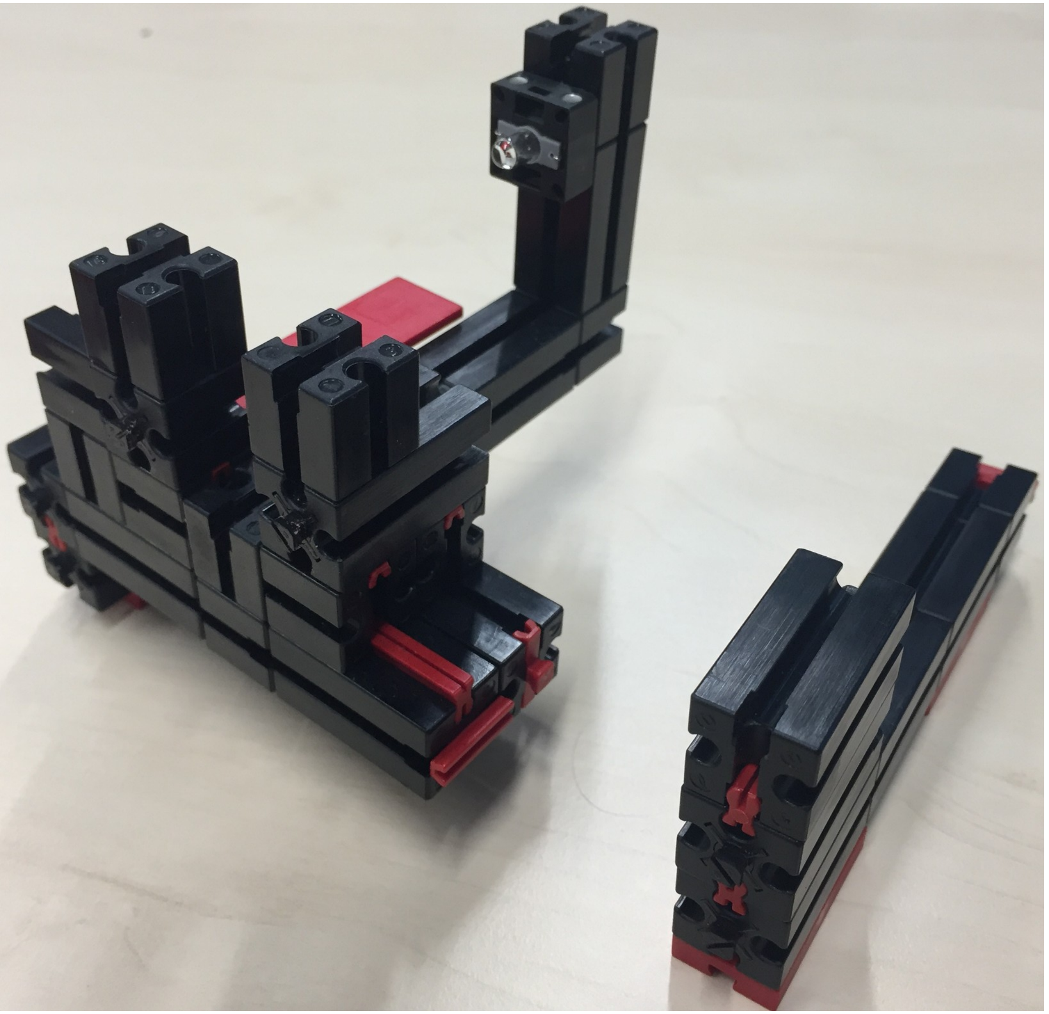
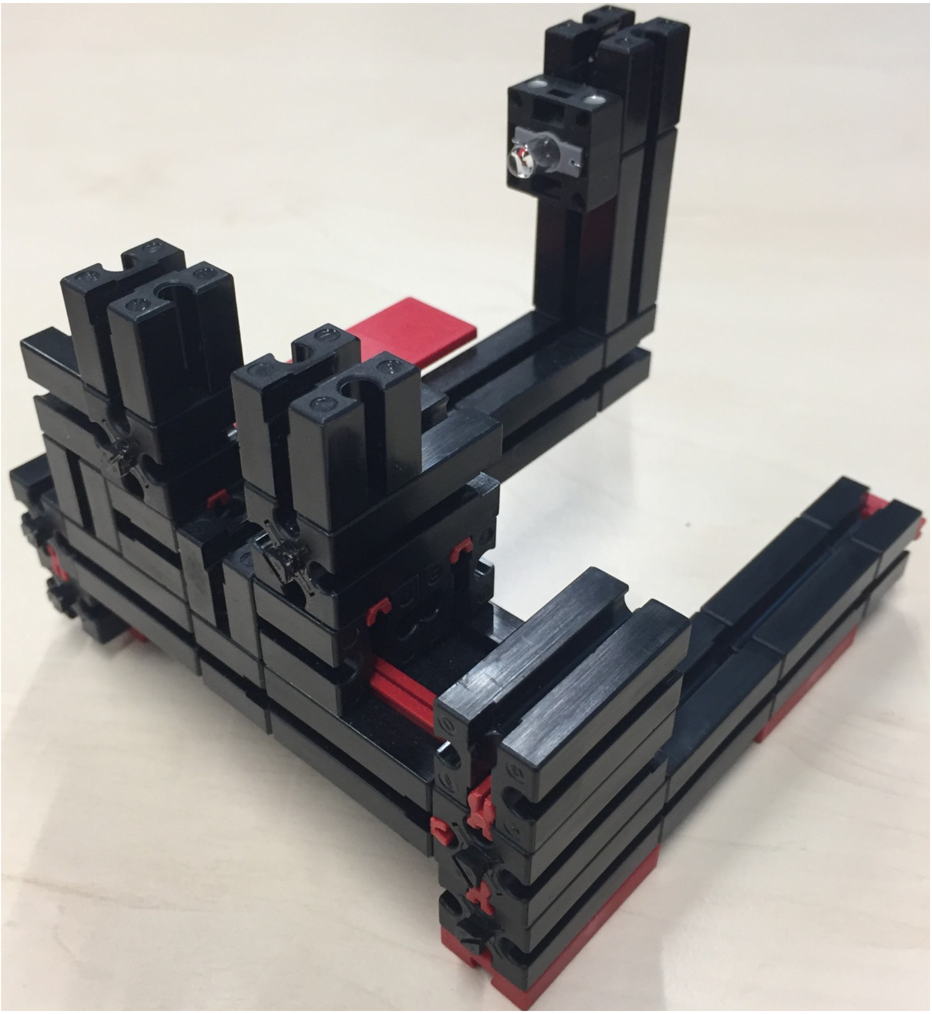

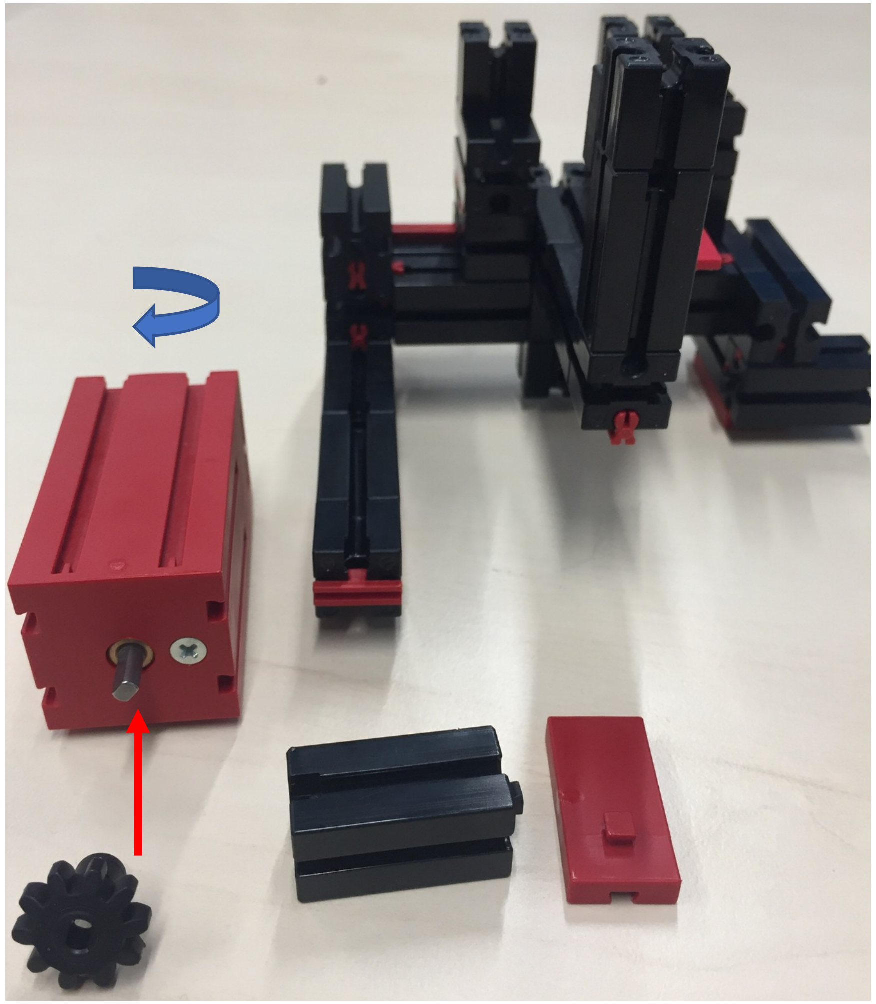

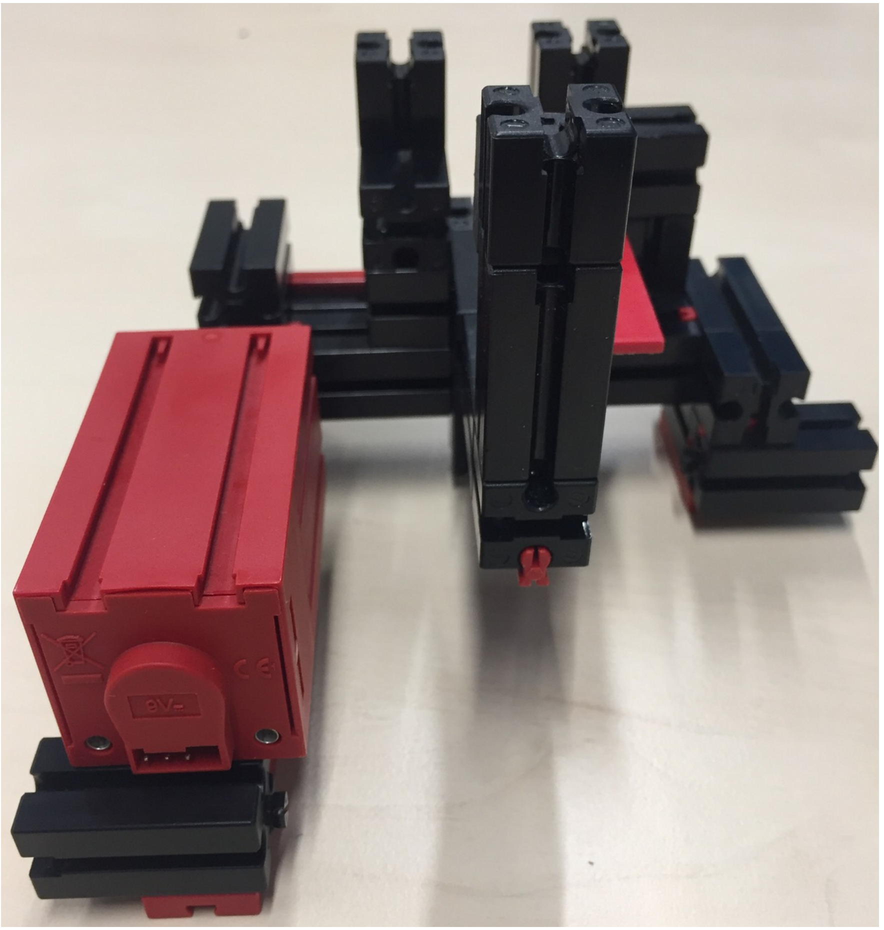

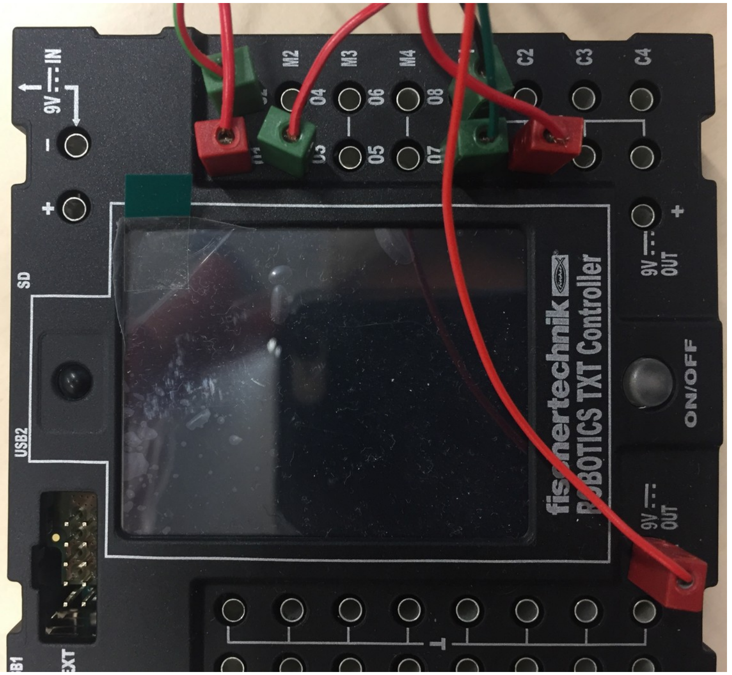
TXT Controller
ON/OFF
9V OUT
USB2
SD

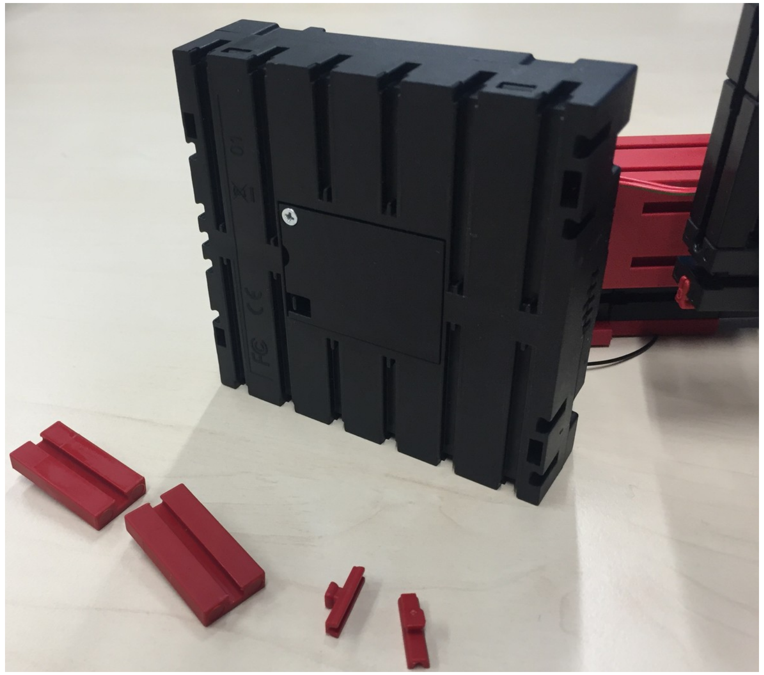

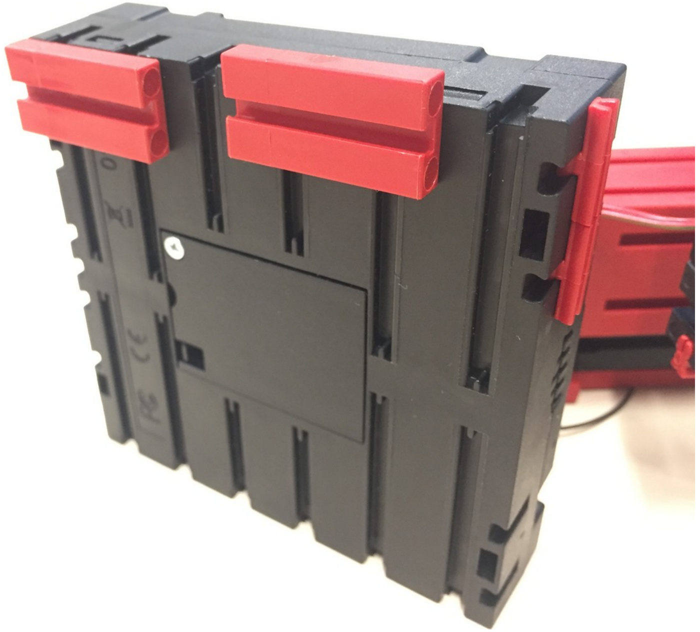

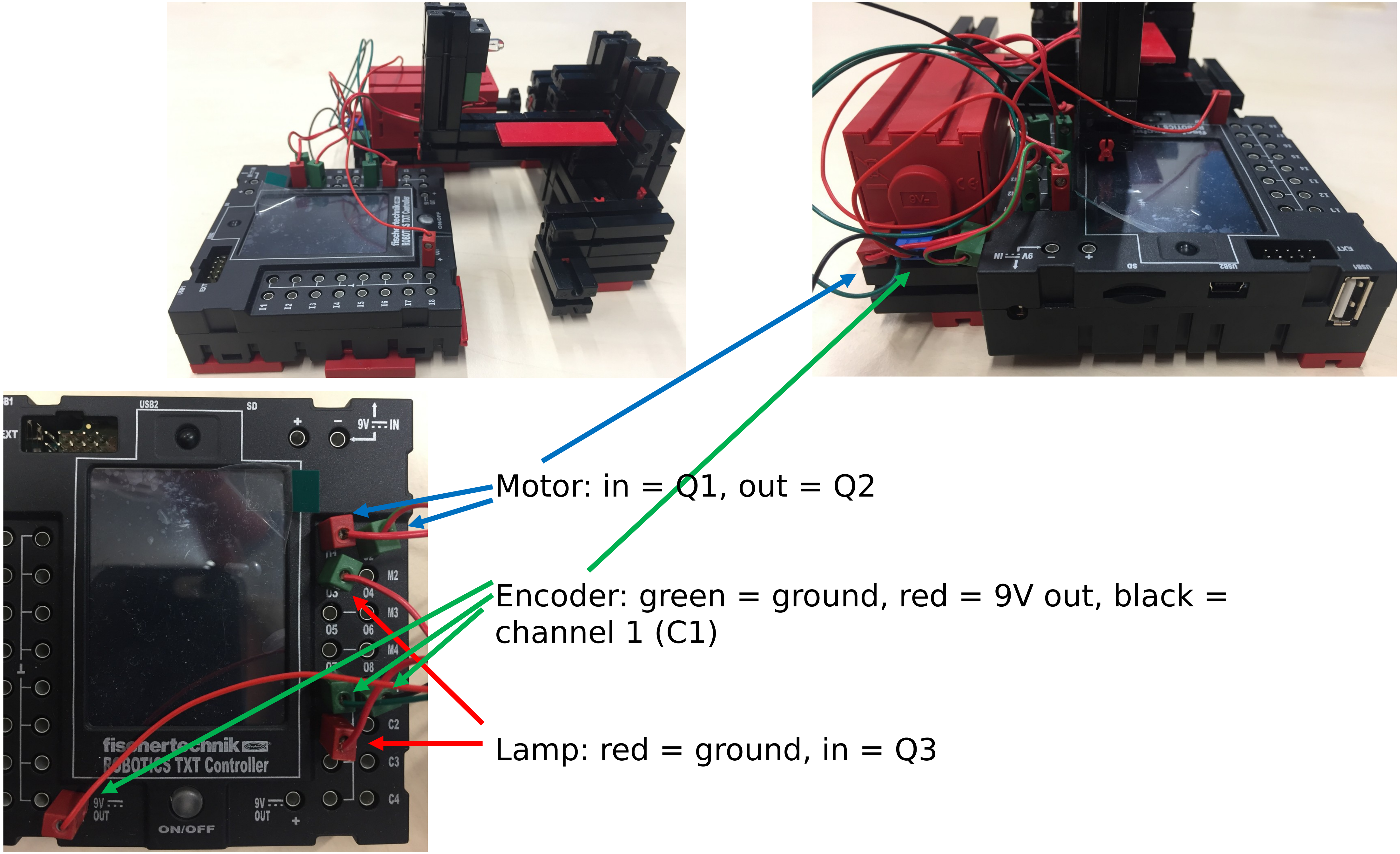
Motor: in = Q1, out = Q2
Encoder: green = ground, red = 9V out, black = channel 1 (C1)
Lamp: red = ground, in = Q3

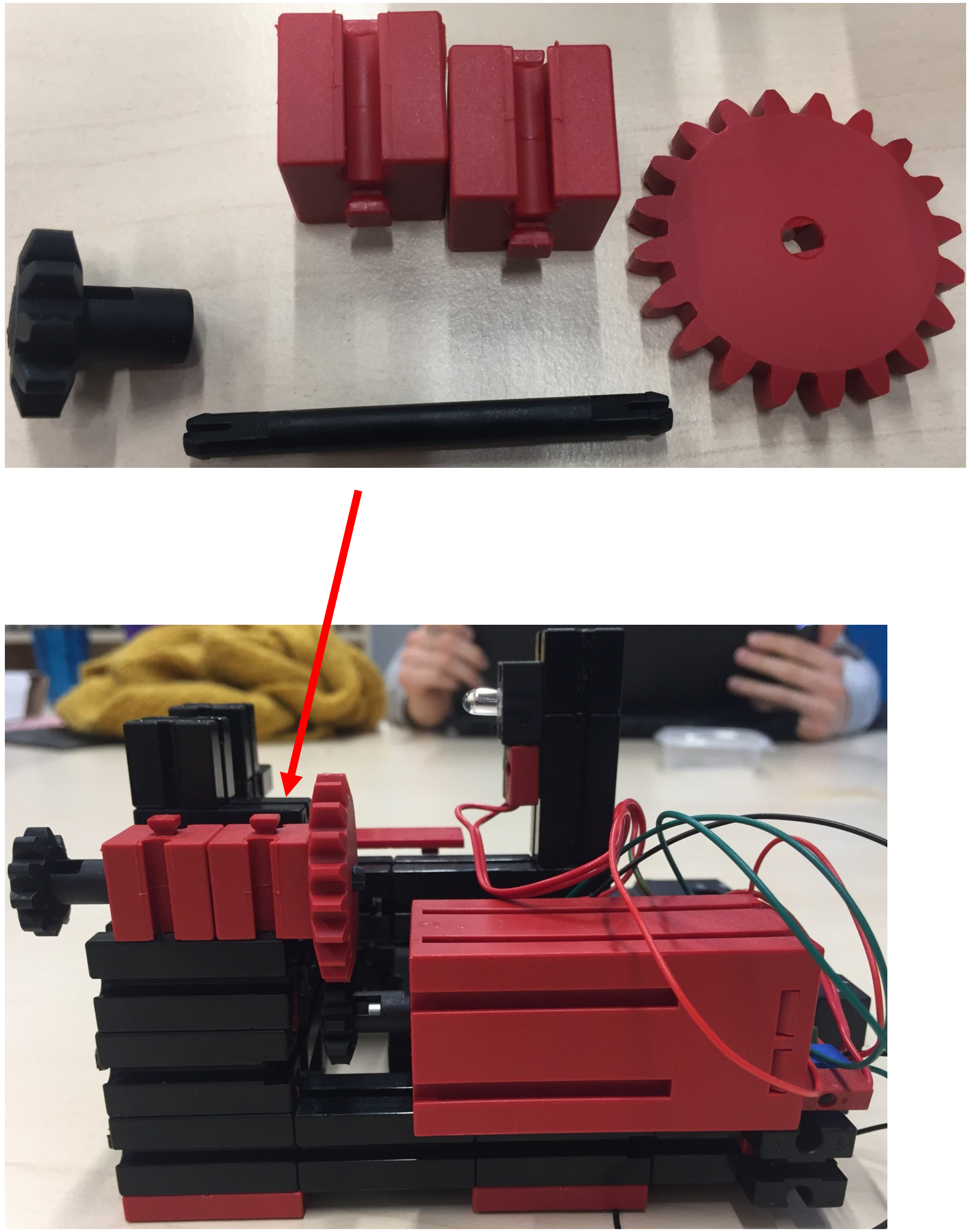

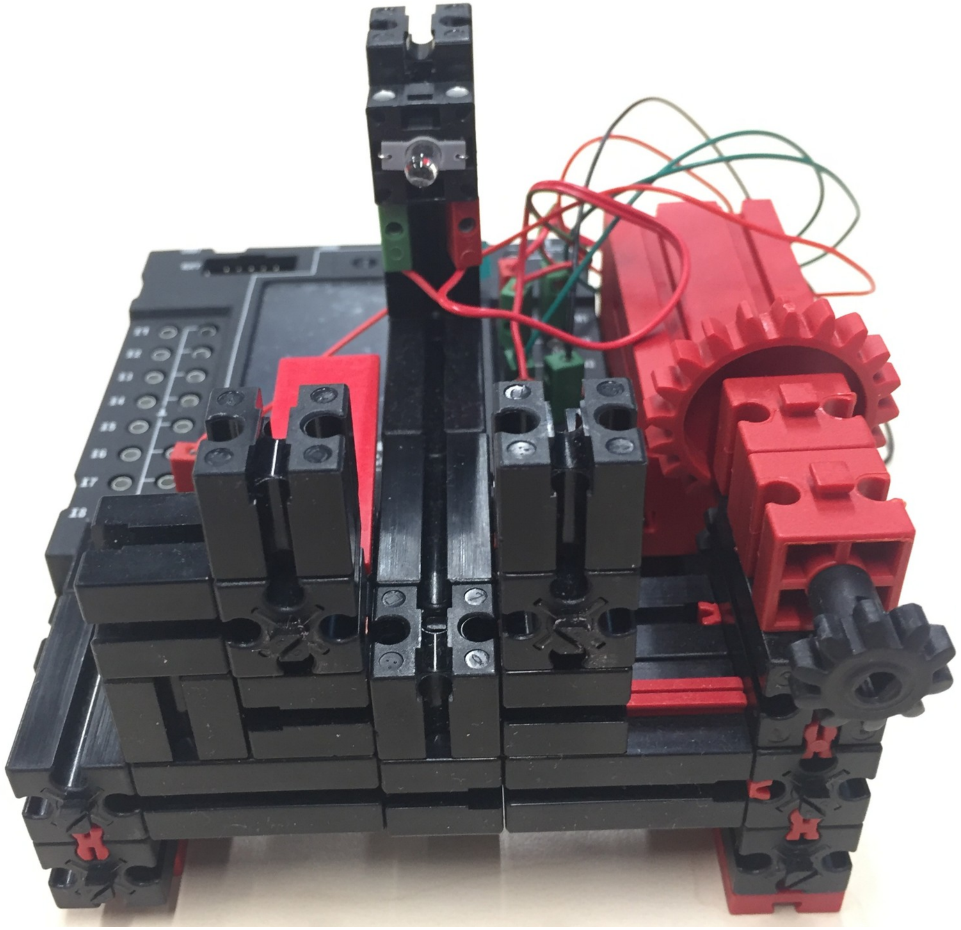

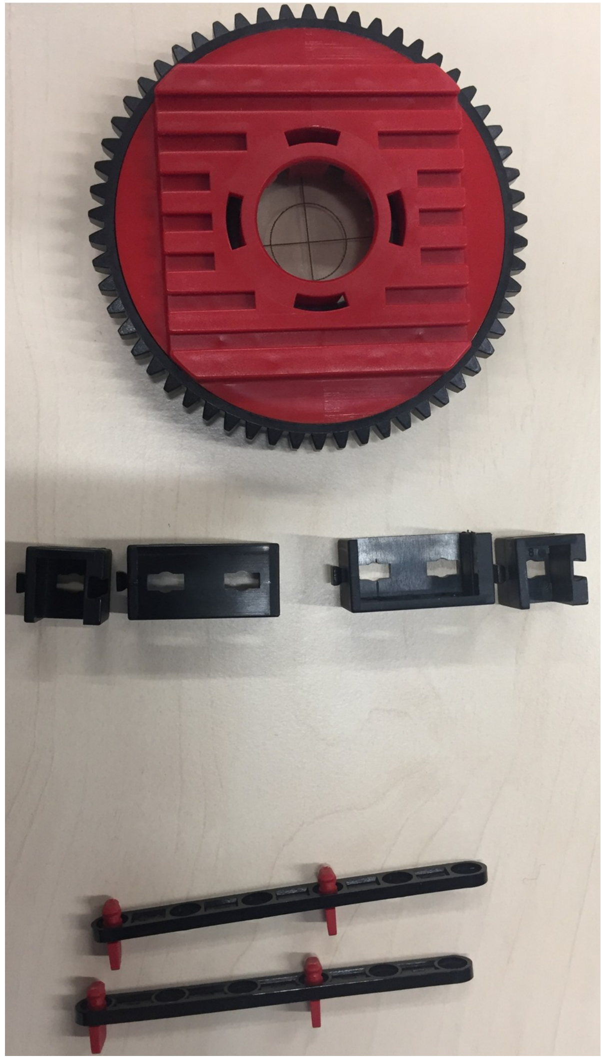

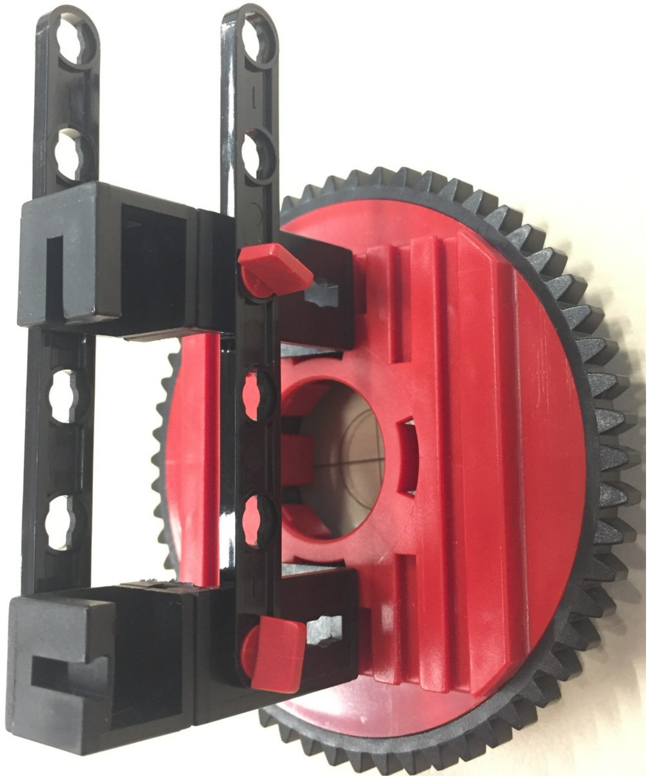

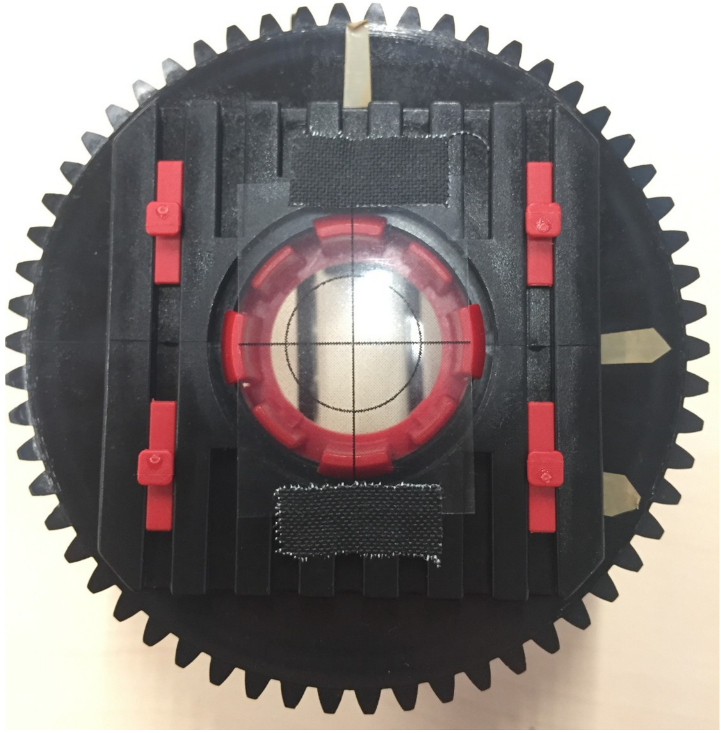

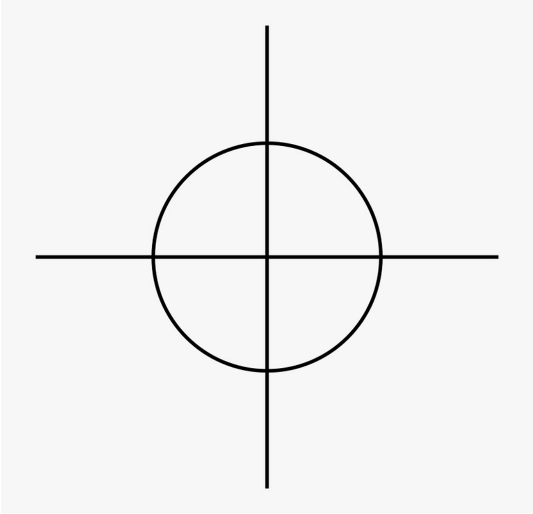

- Crosshairs can be printed on monochrome transparents via laser printer
- Roughly centered and fixed with black tape

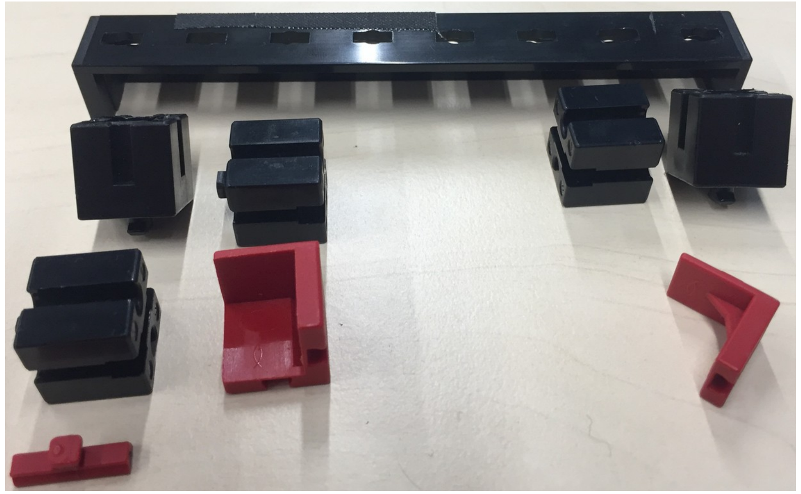

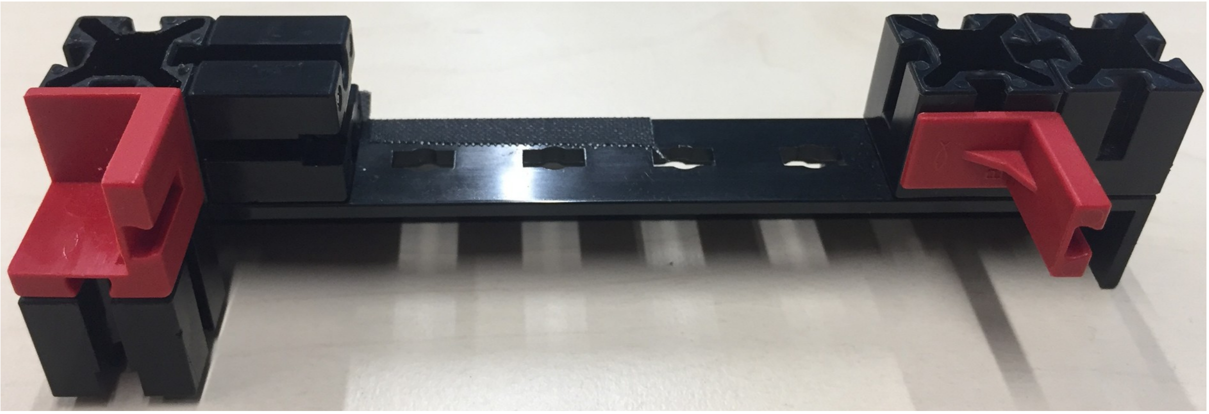

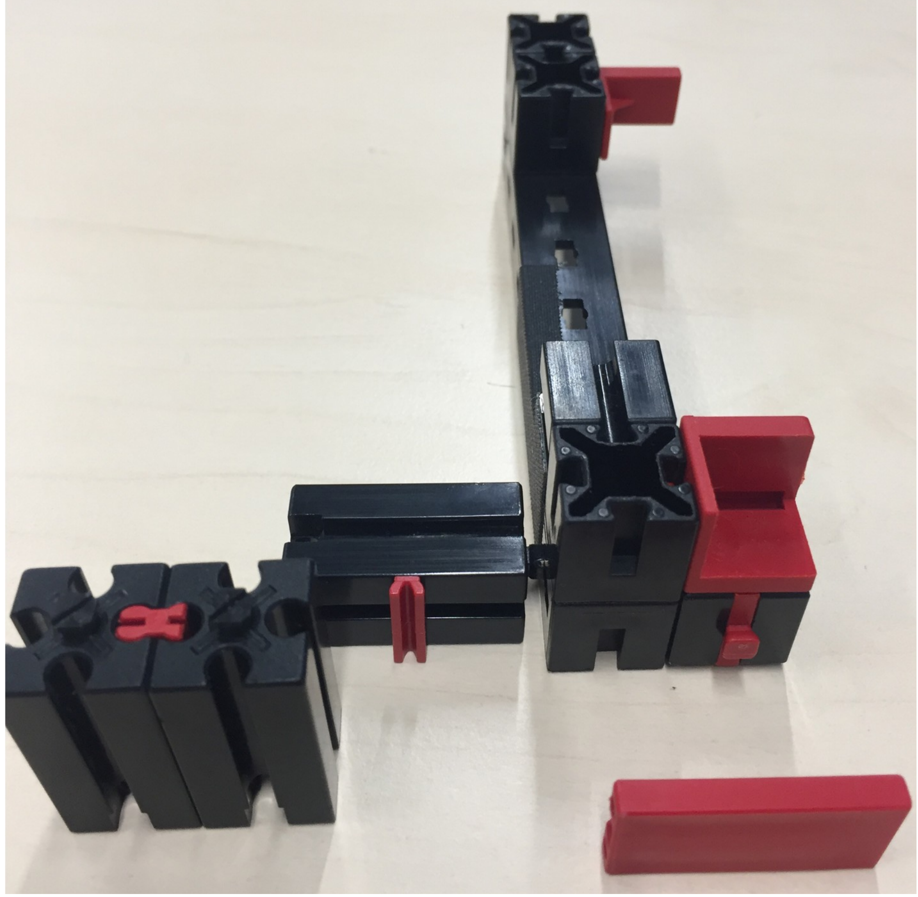

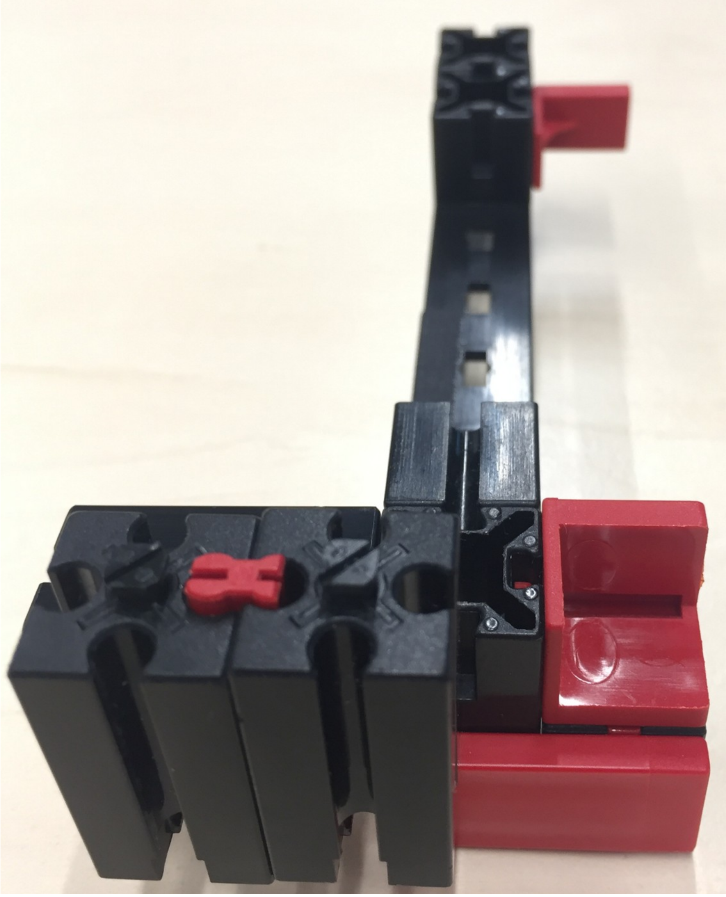

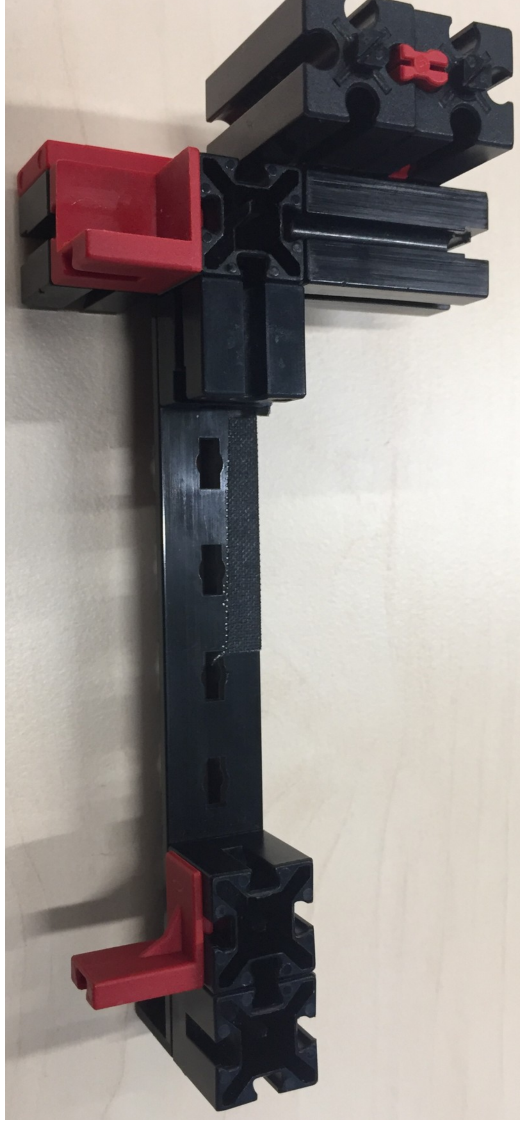

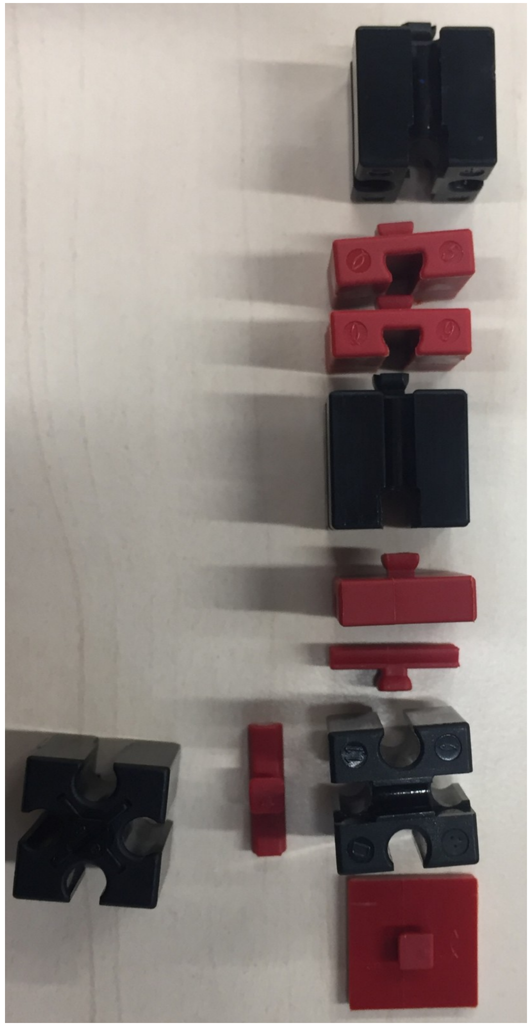
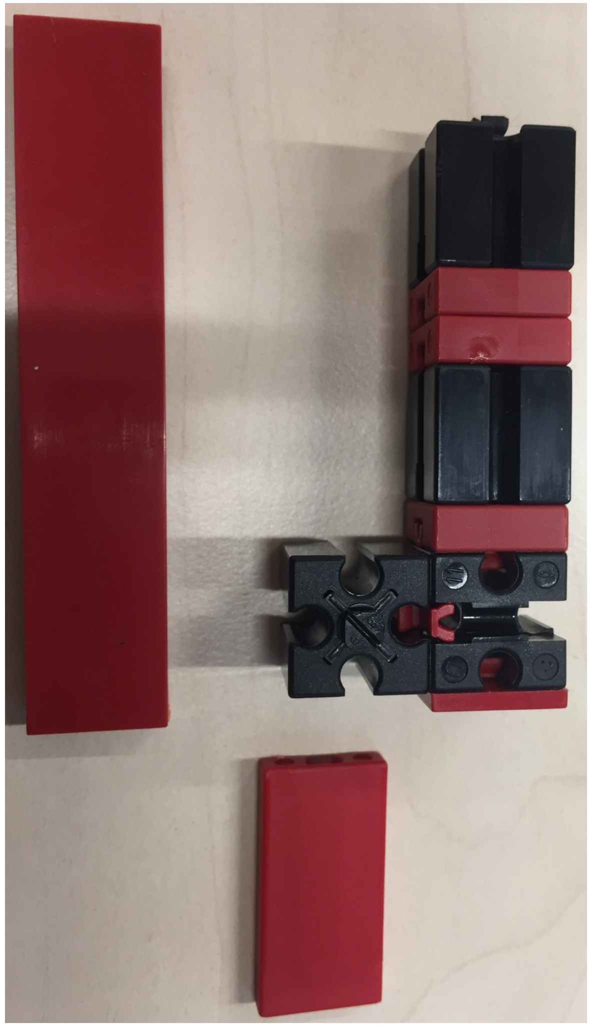
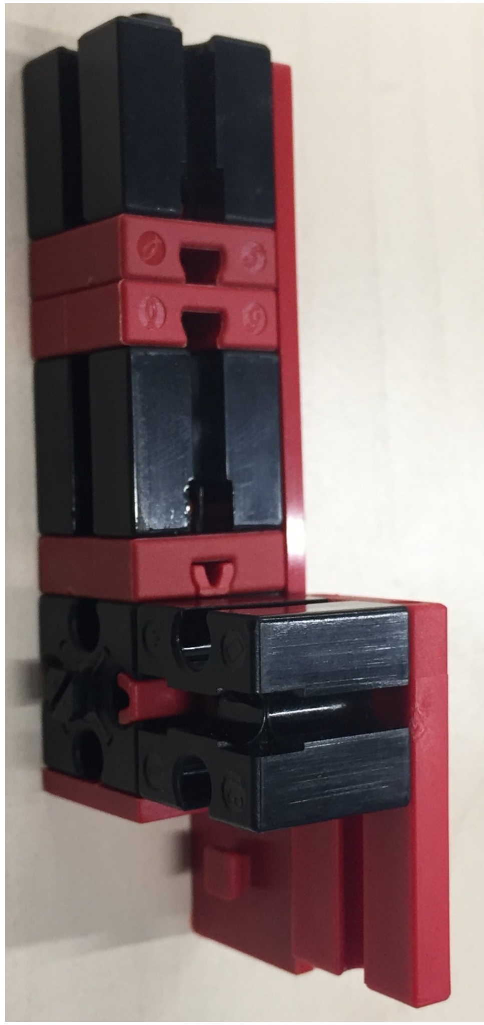

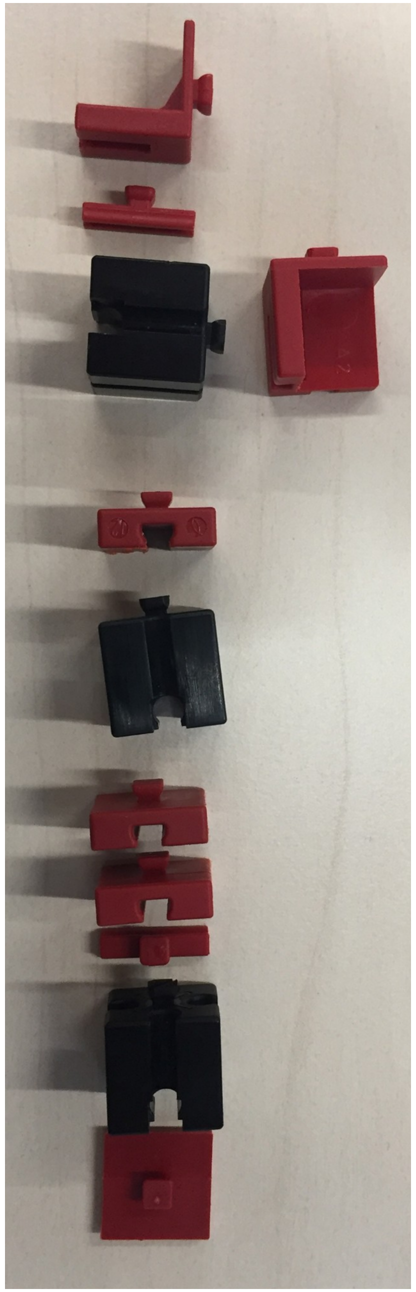

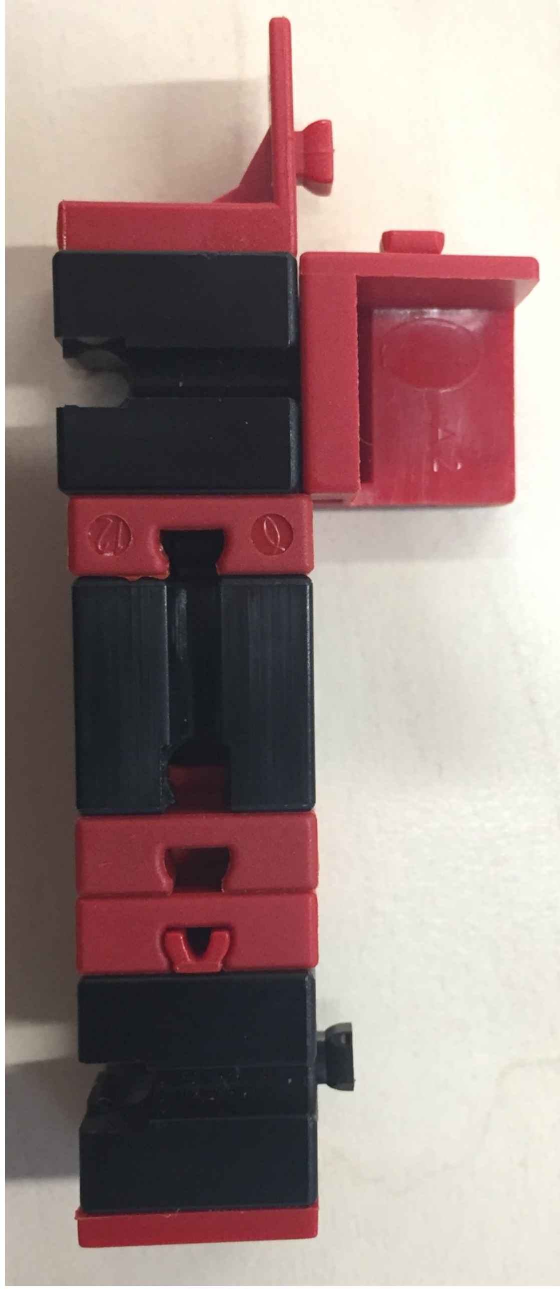

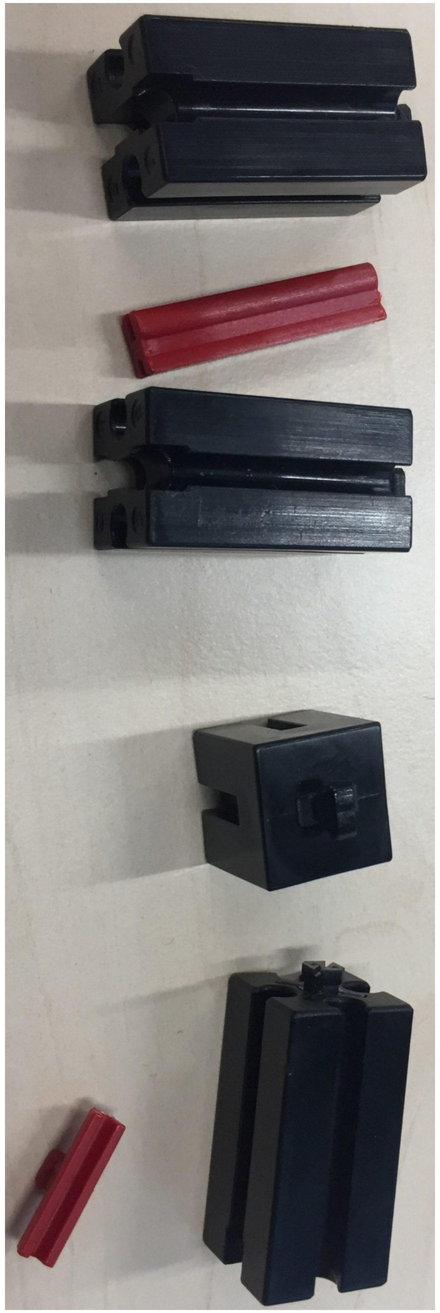
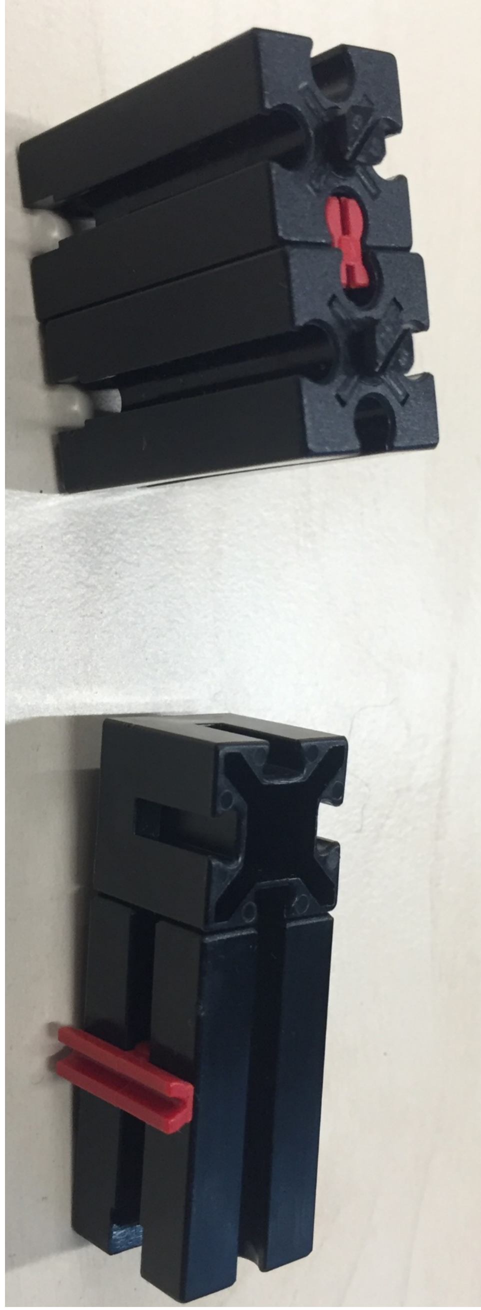
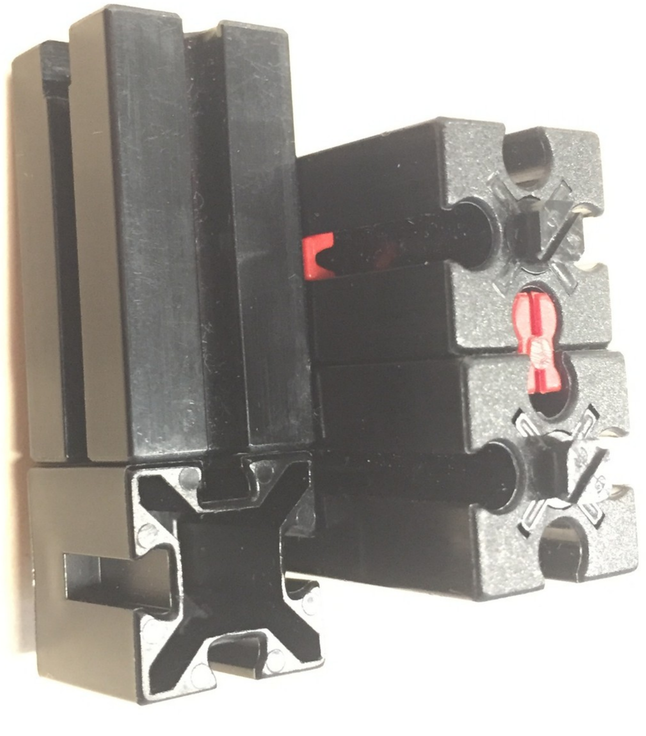

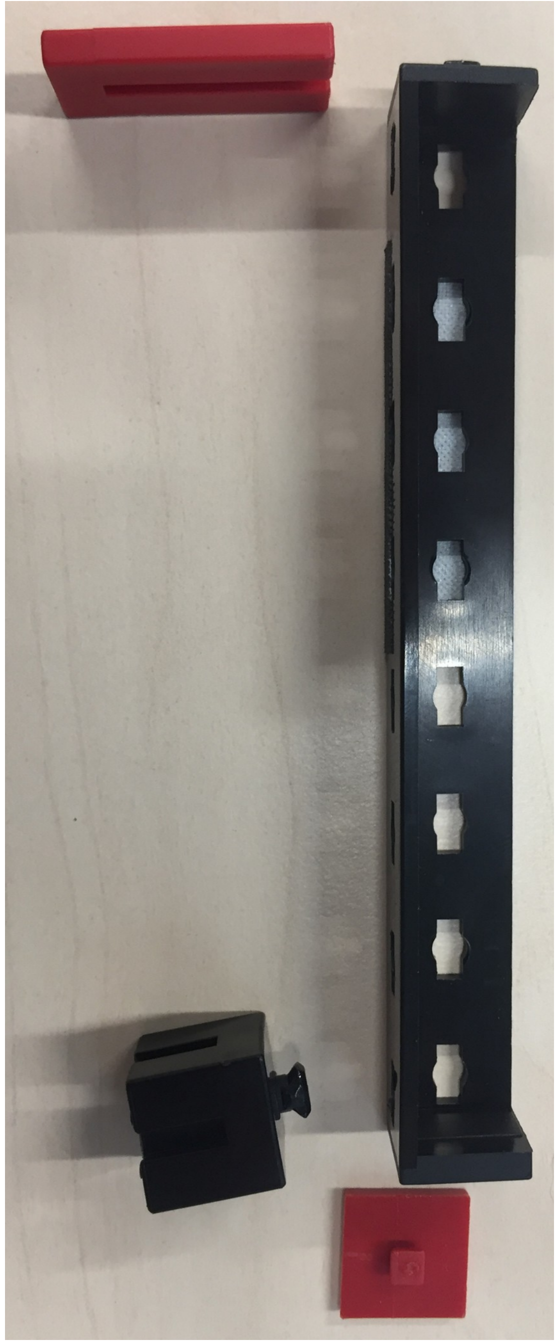
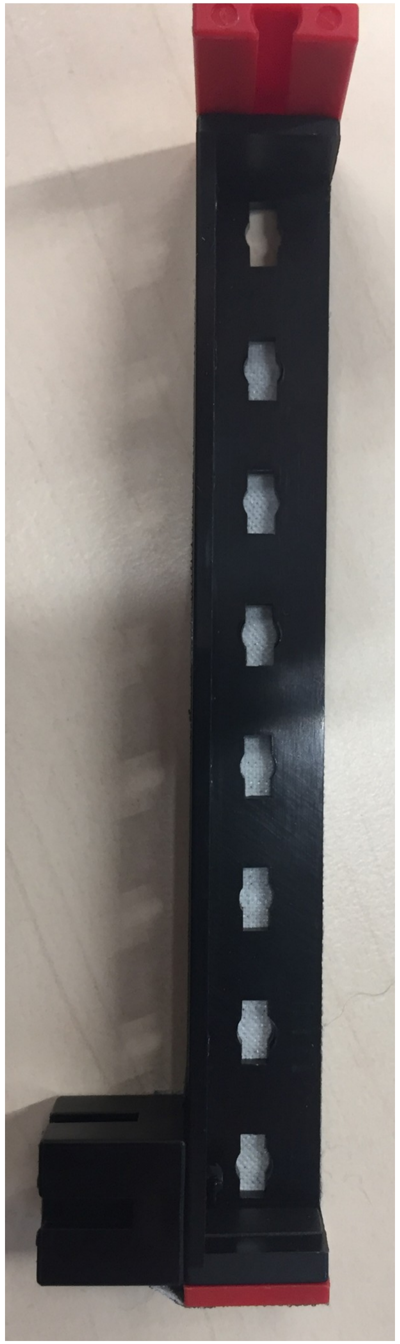
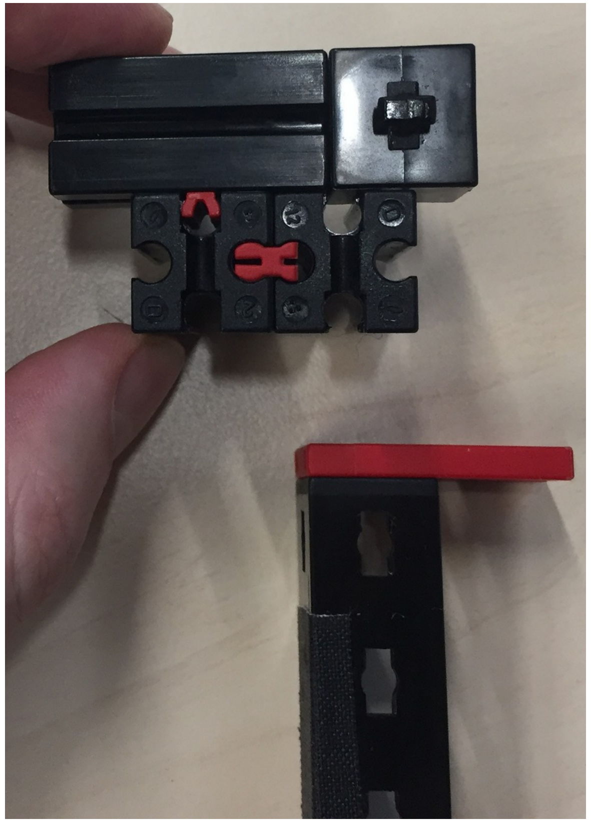
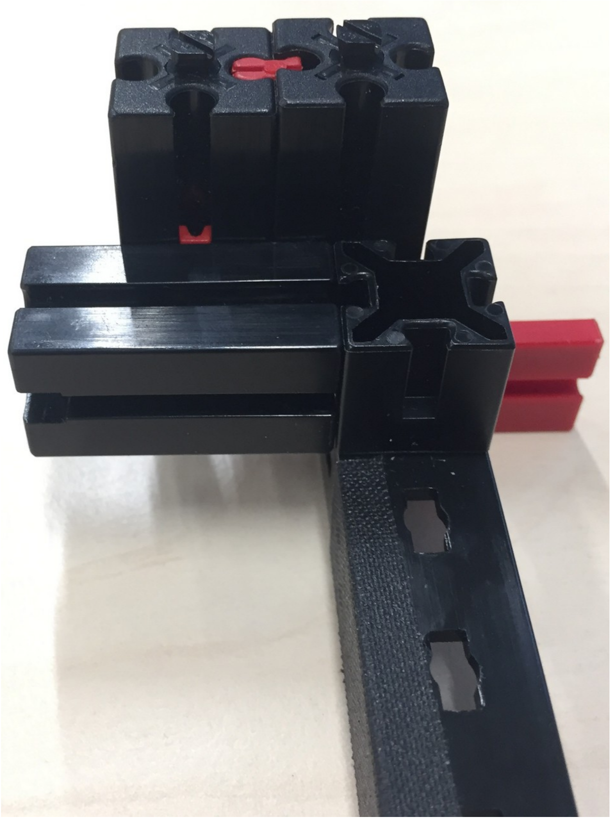

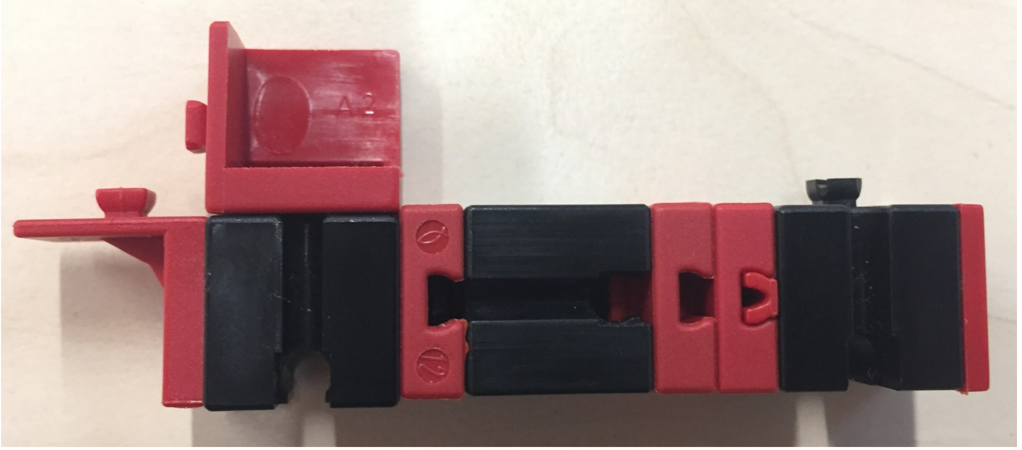
A2

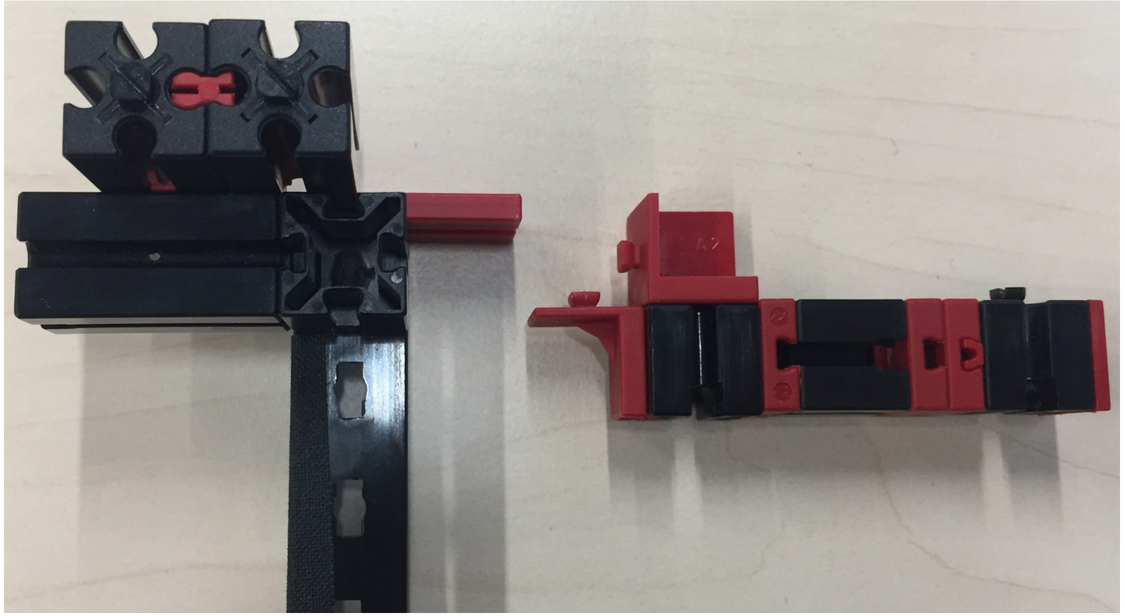

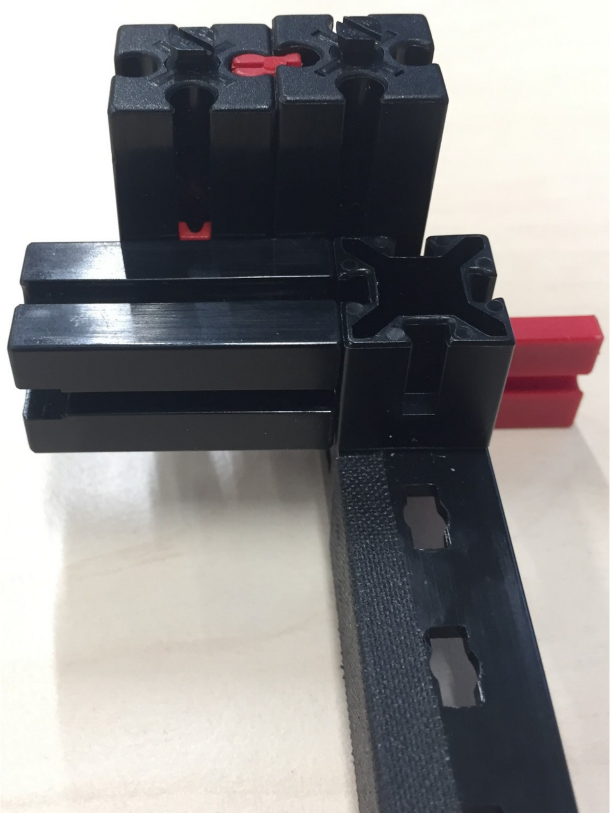

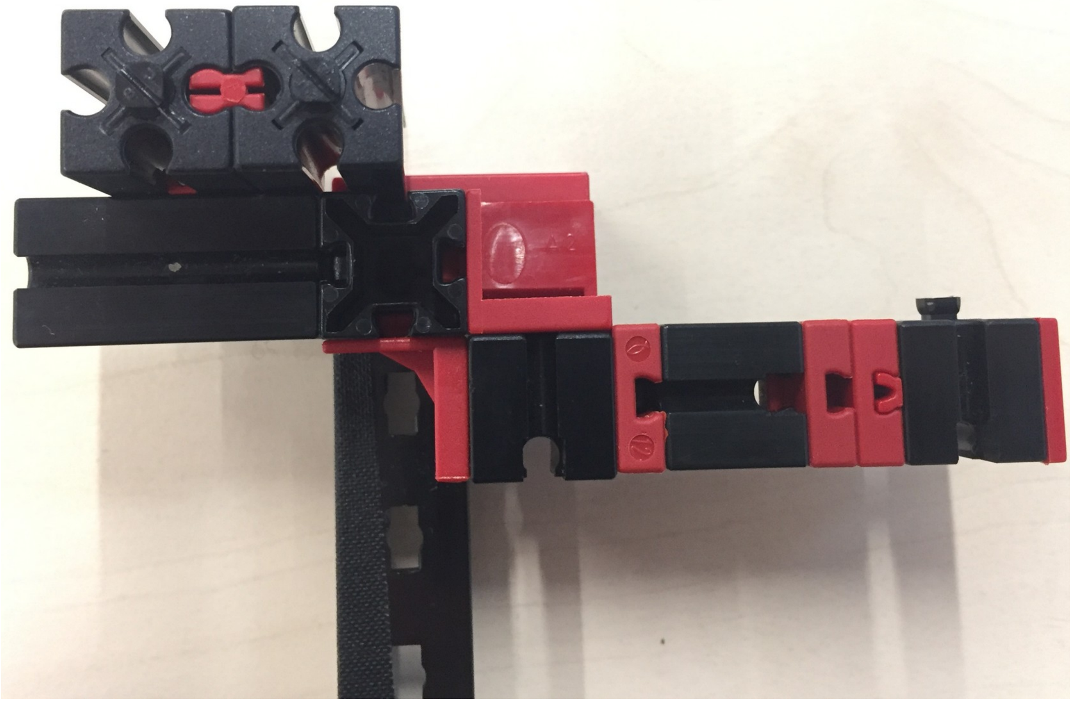

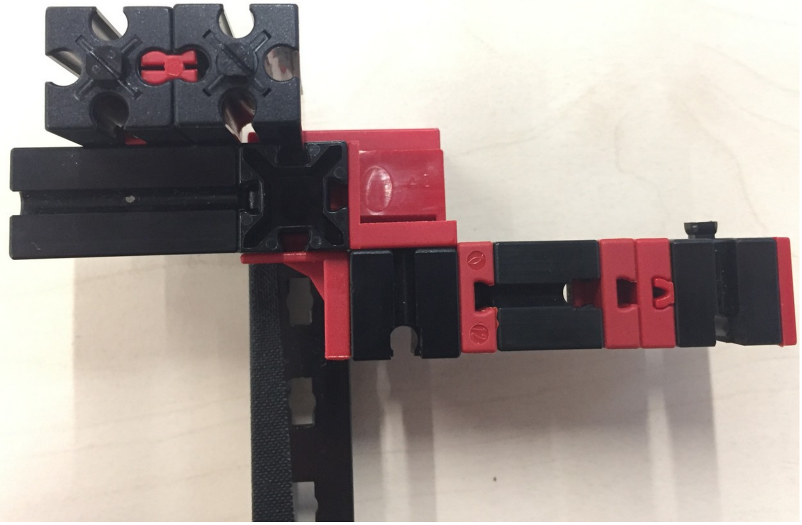

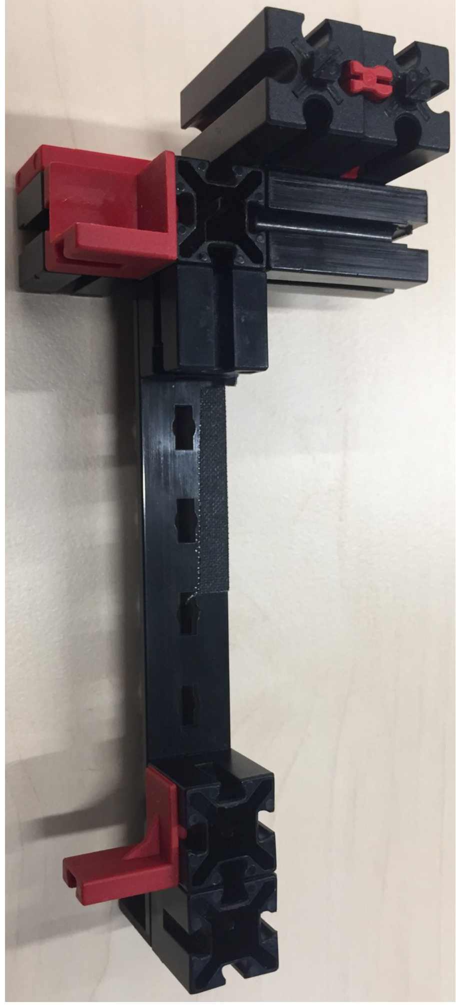

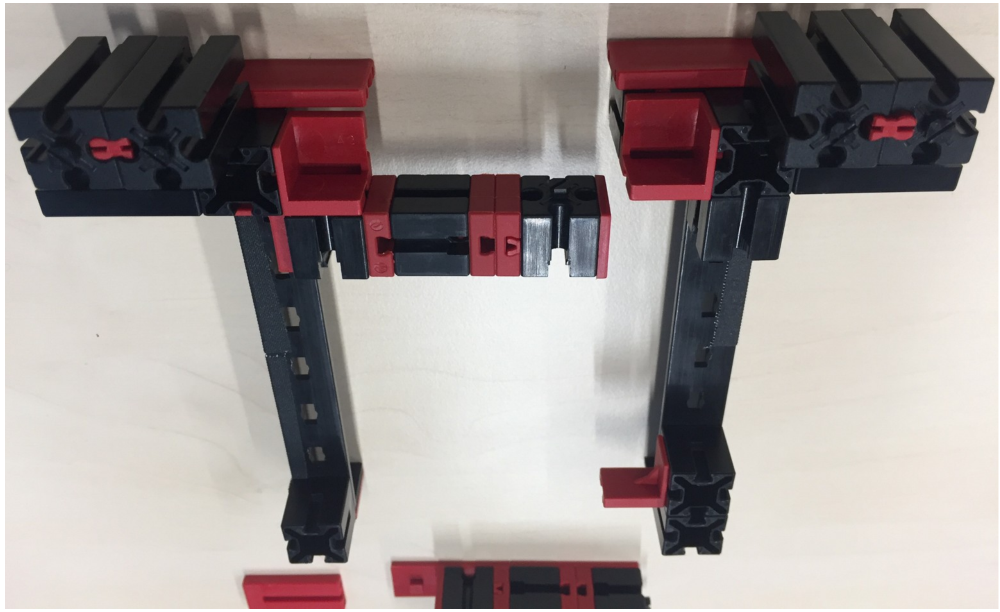

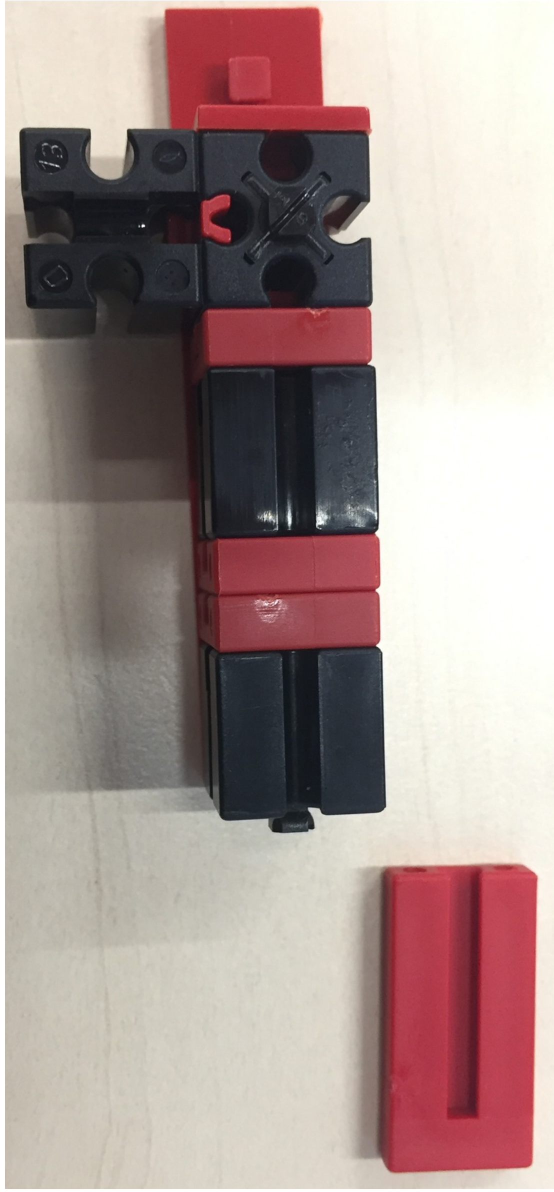

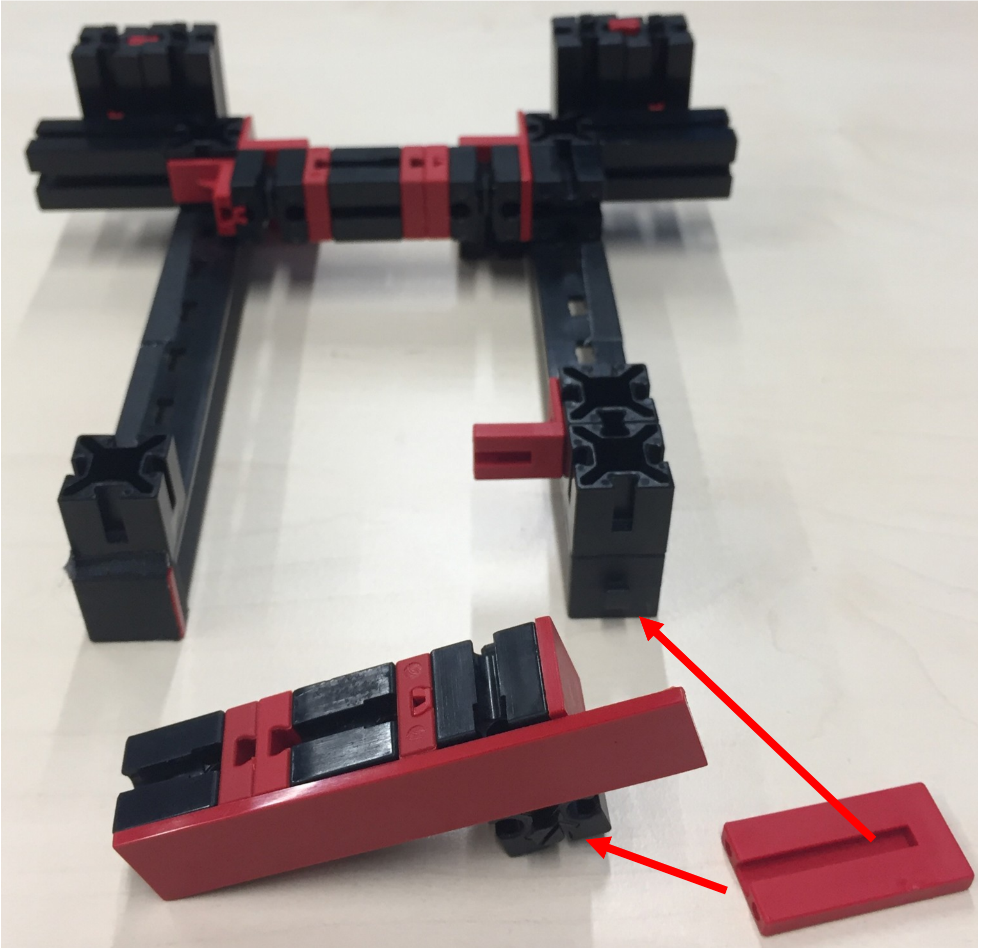

Multiwellplate-Adapter Electronic-Setup

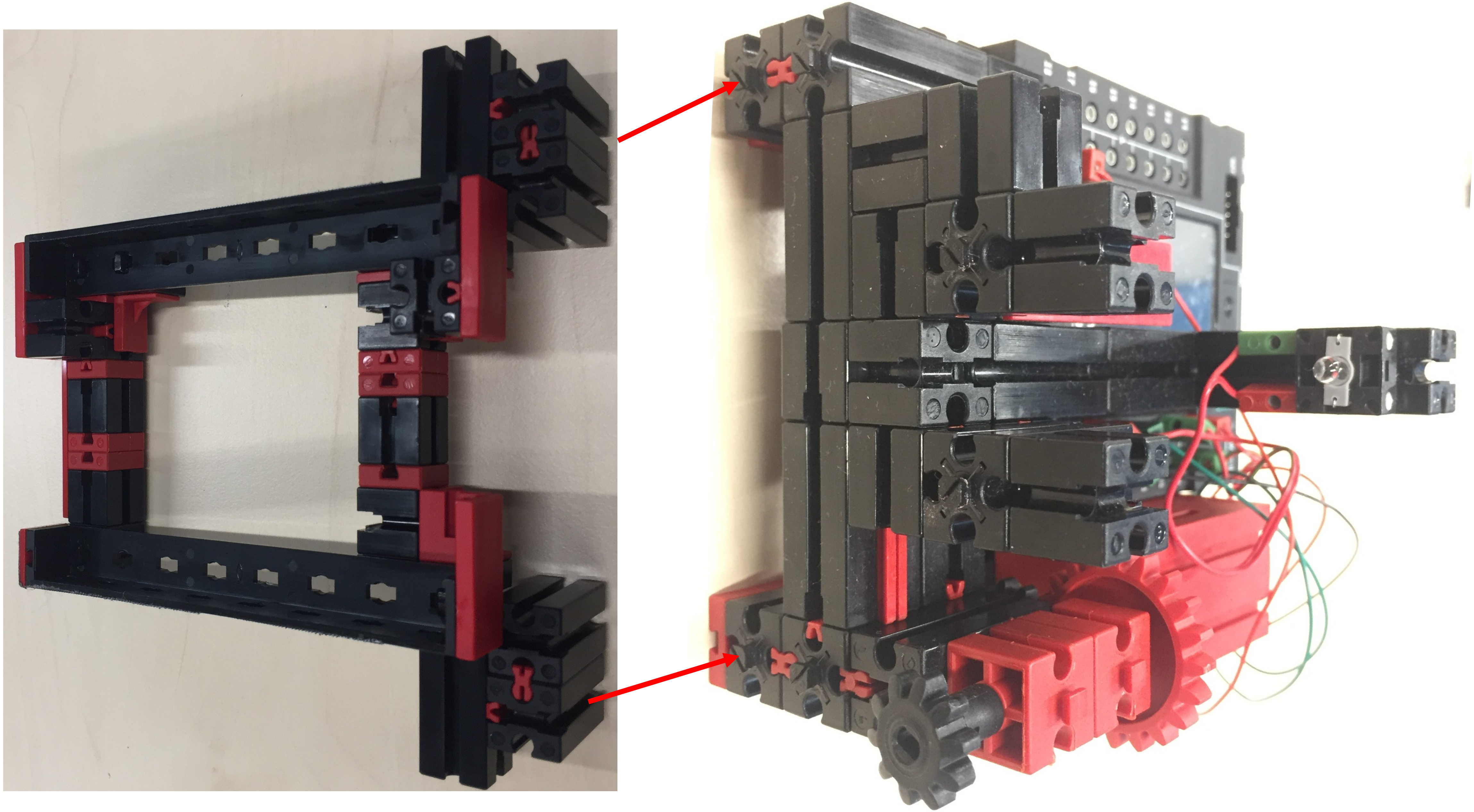
