## Supplementary material for "GraviKit: an easy-to-implement microscope add-on for observation of gravitation dependent processes": ftConstruction_manualStage (construction manual)

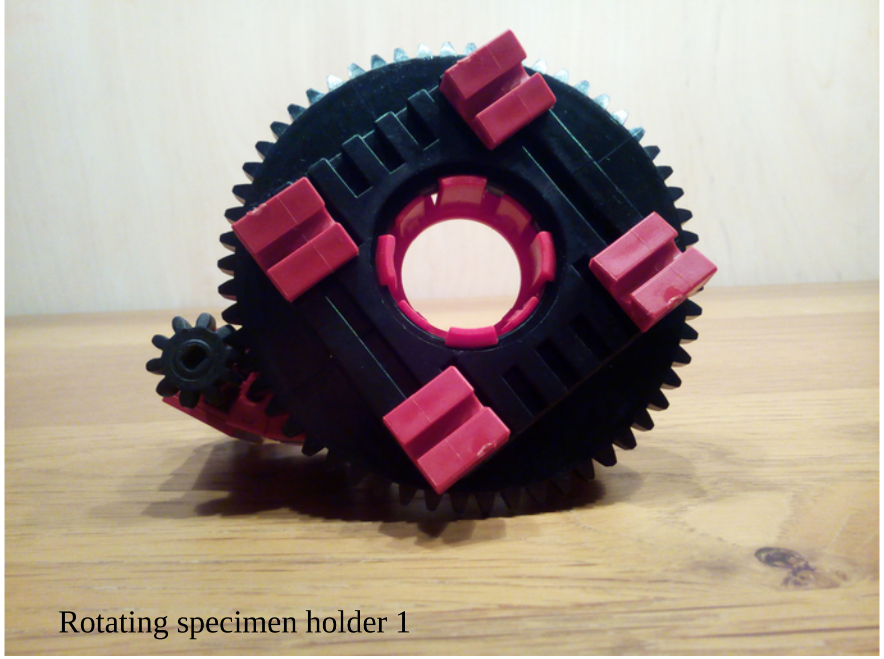

Rotating specimen holder 1

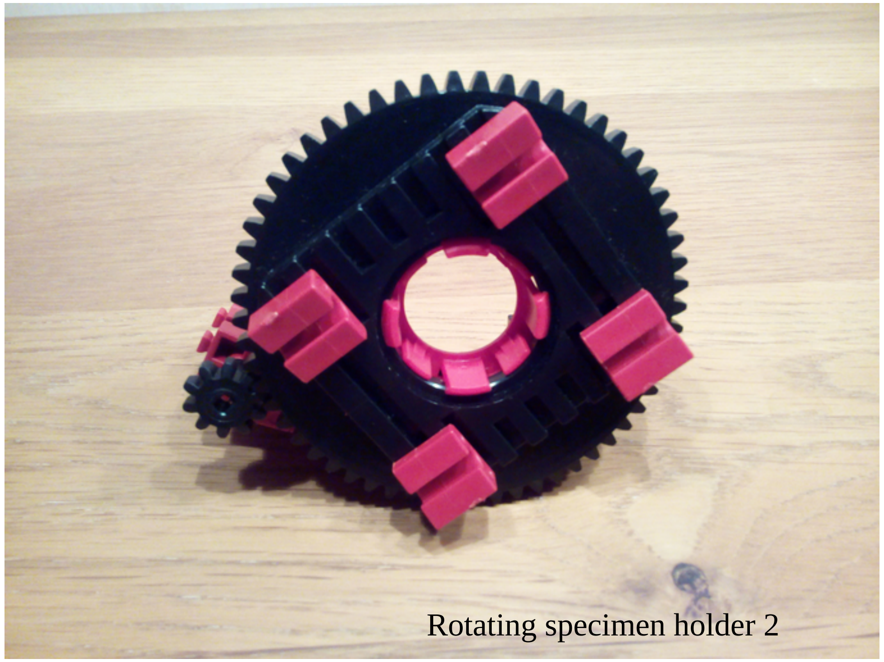

Rotating specimen holder 2

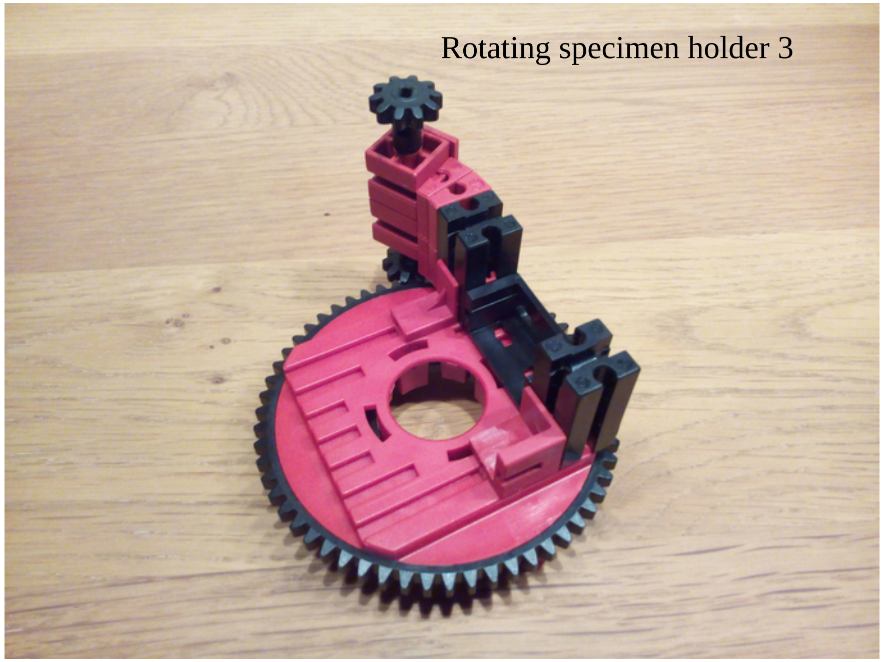

Rotating specimen holder 3

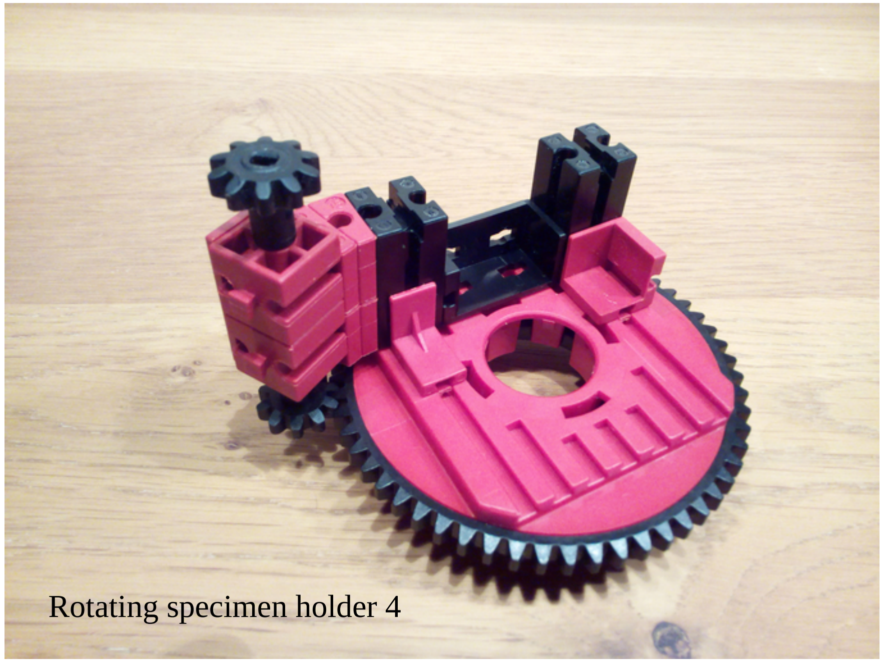

Rotating specimen holder 4

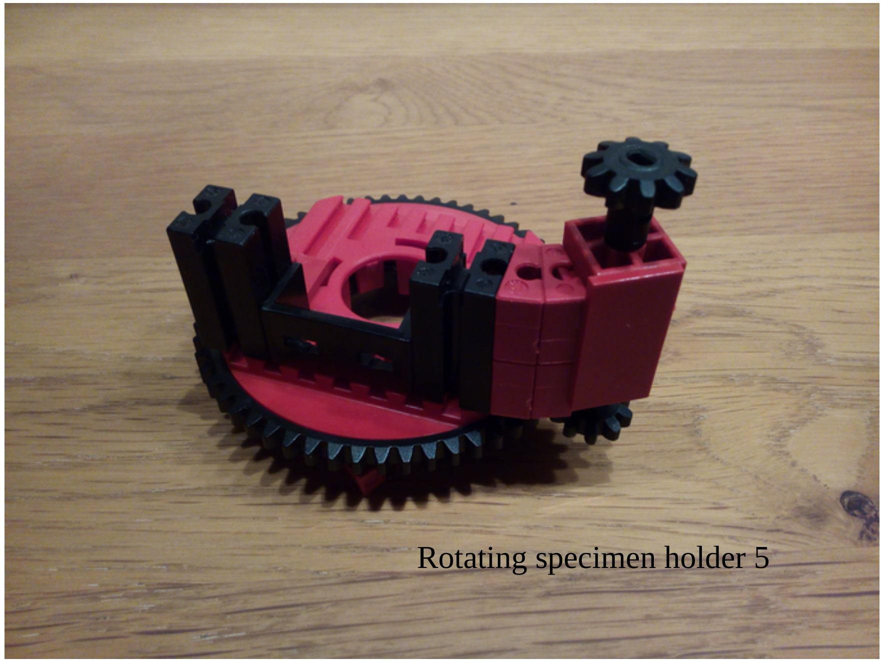
Rotating specimen holder 5

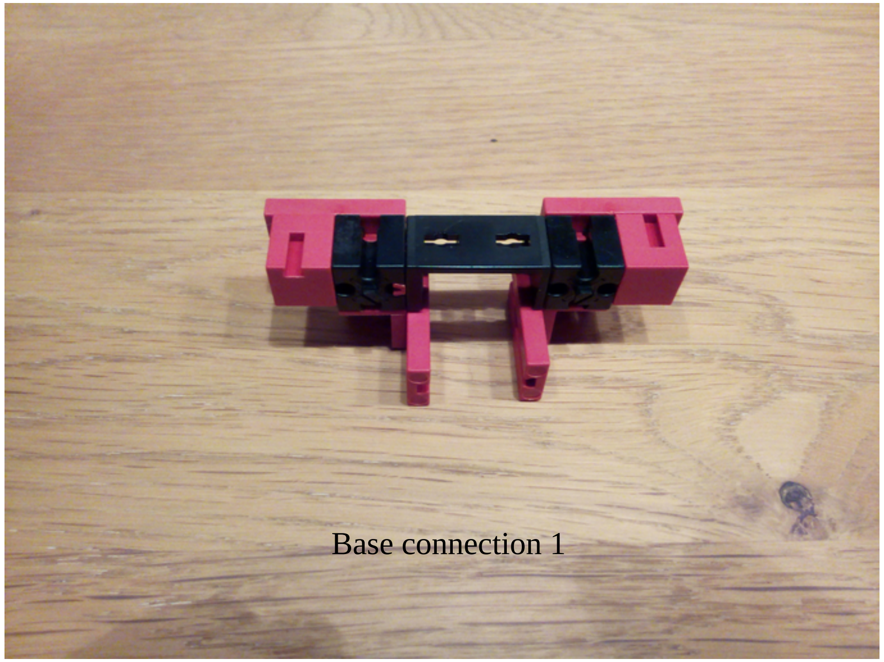
Base connection 1

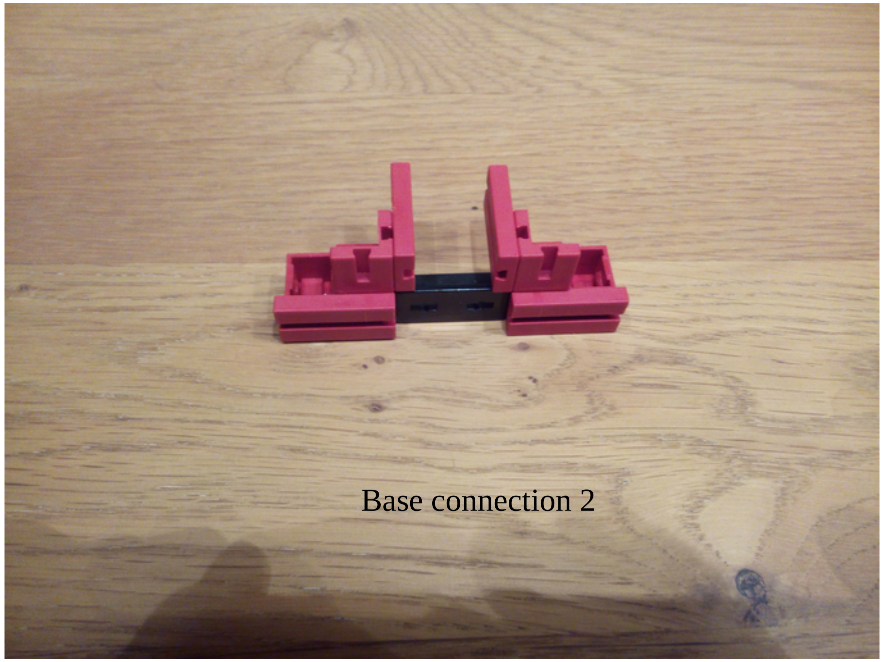
Base connection 2

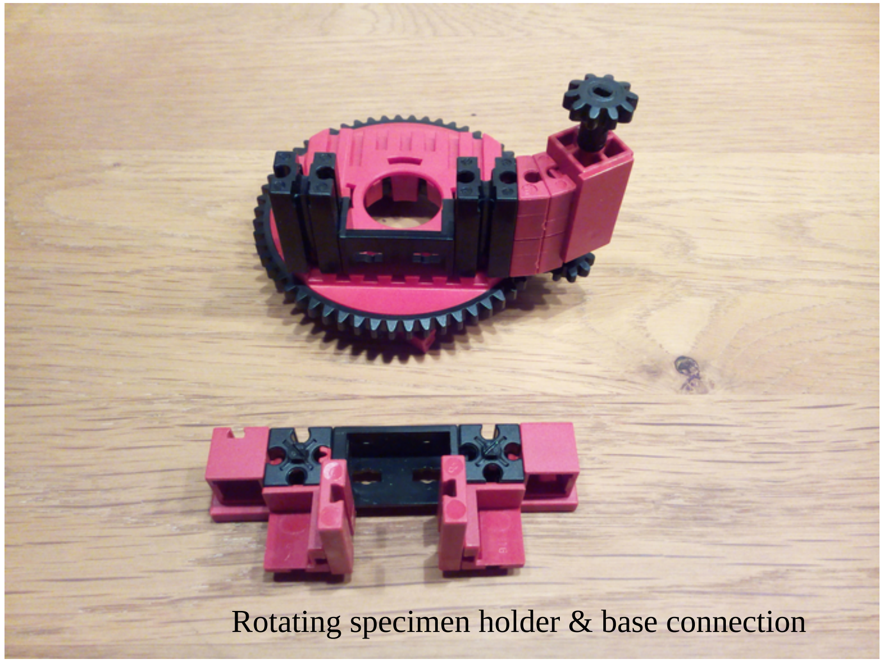
Rotating specimen holder & base connection

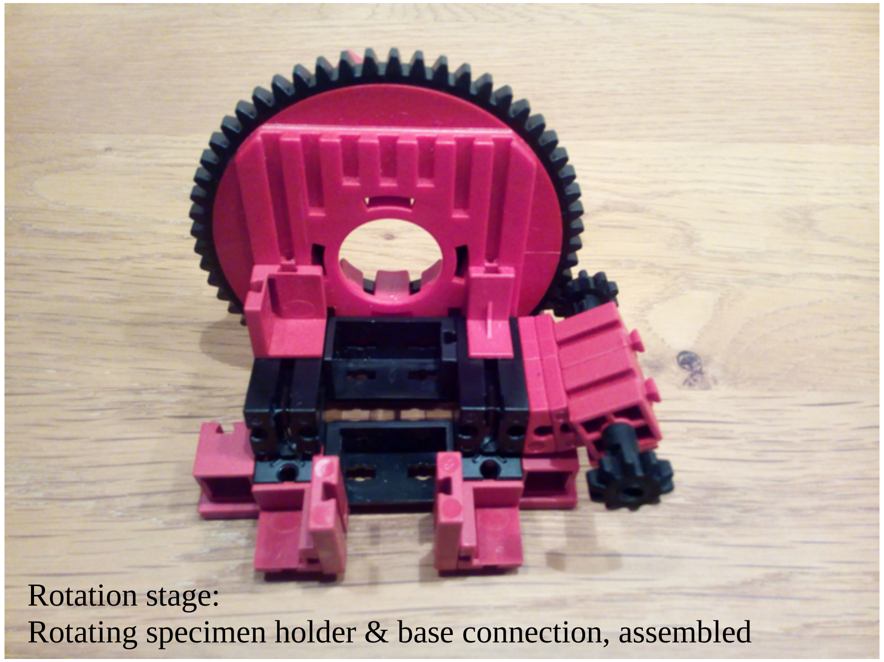

Rotation stage:
Rotating specimen holder & base connection, assembled

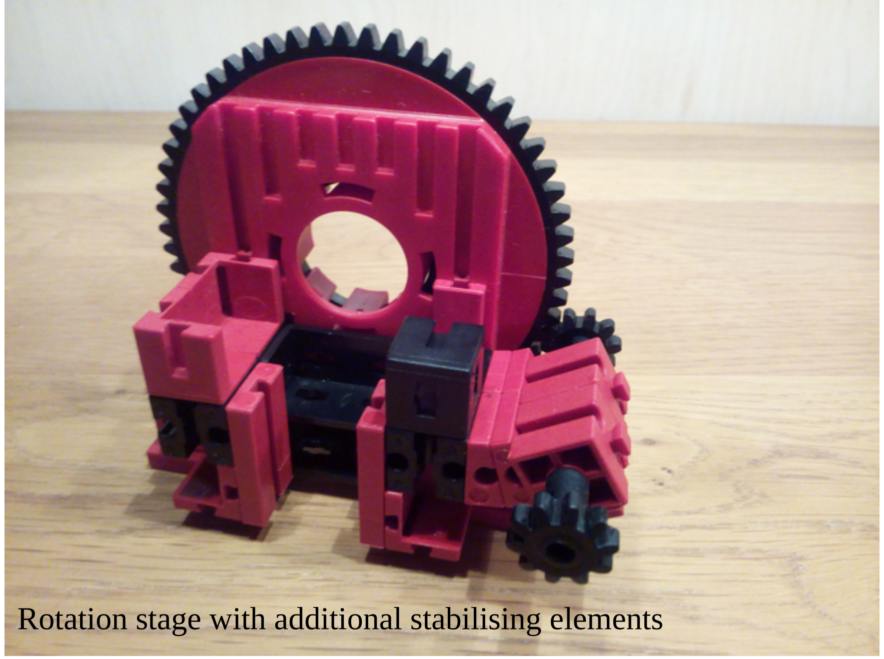

Rotation stage with additional stabilising elements

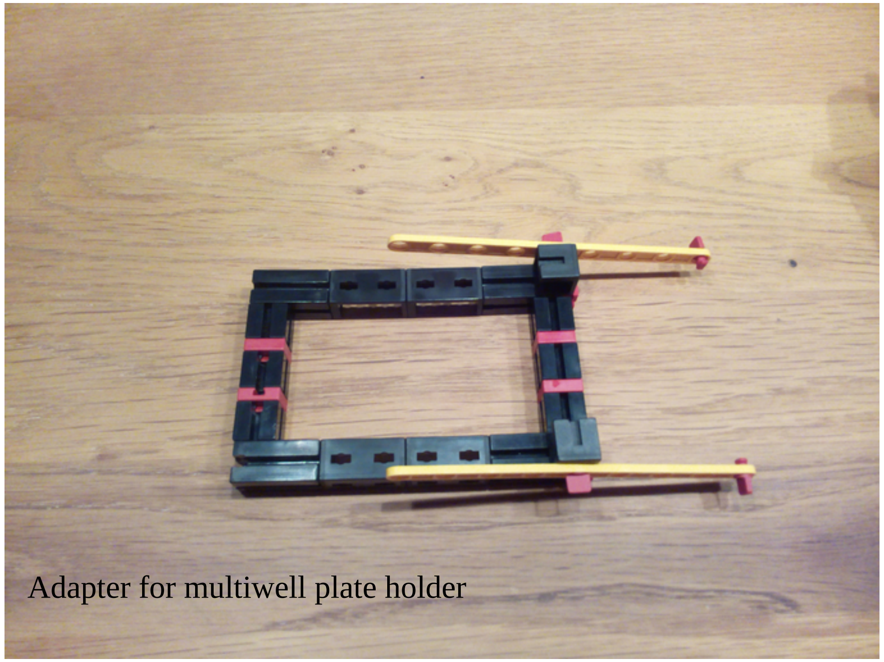

Adapter for multiwell plate holder

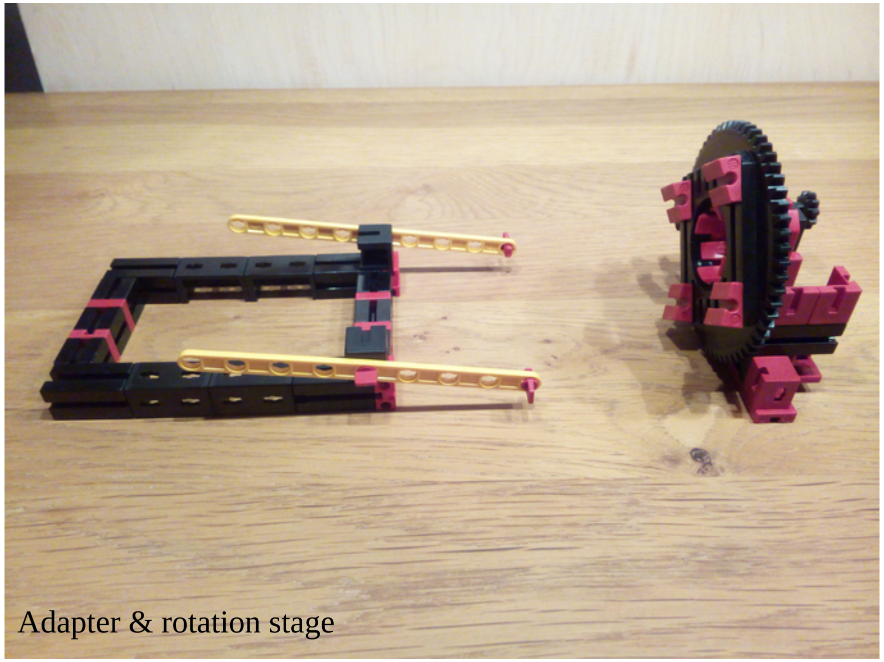

Adapter & rotation stage

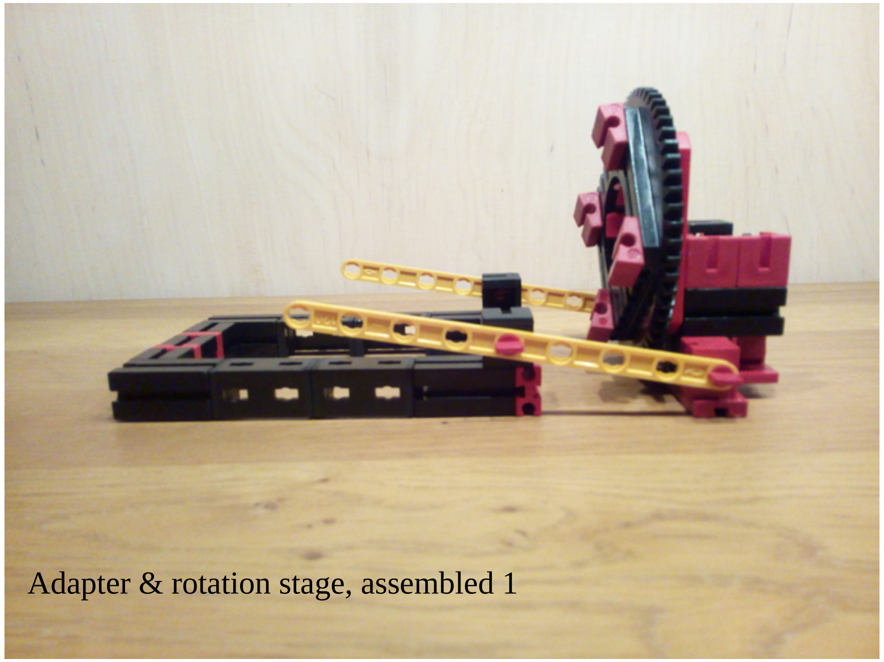

Adapter & rotation stage, assembled 1

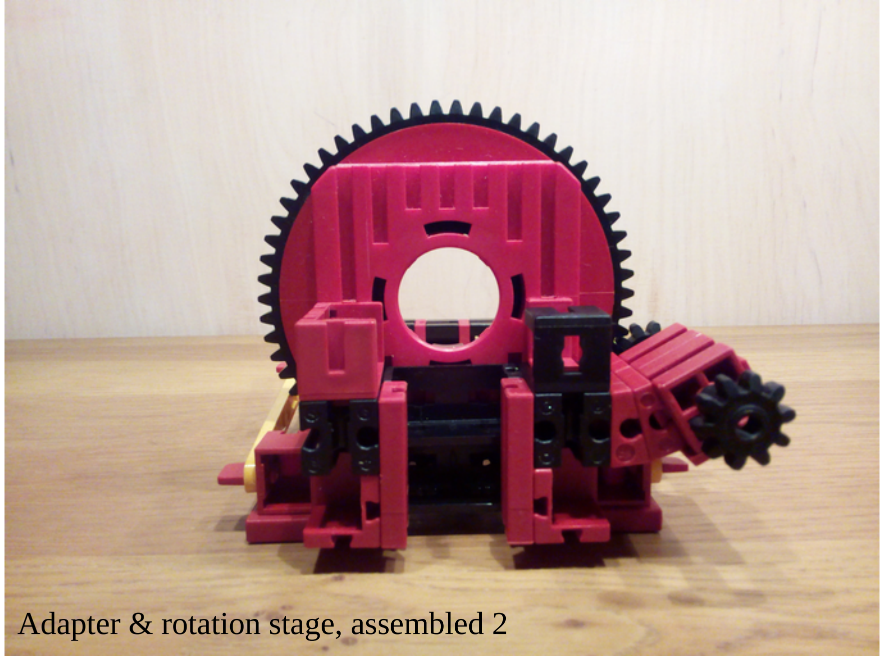

Adapter & rotation stage, assembled 2

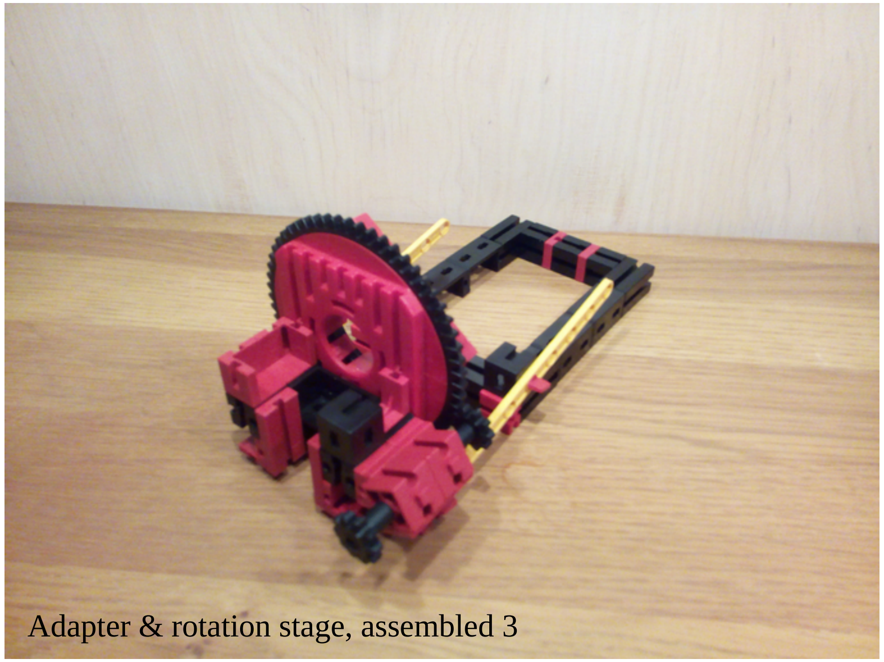

Adapter & rotation stage, assembled 3

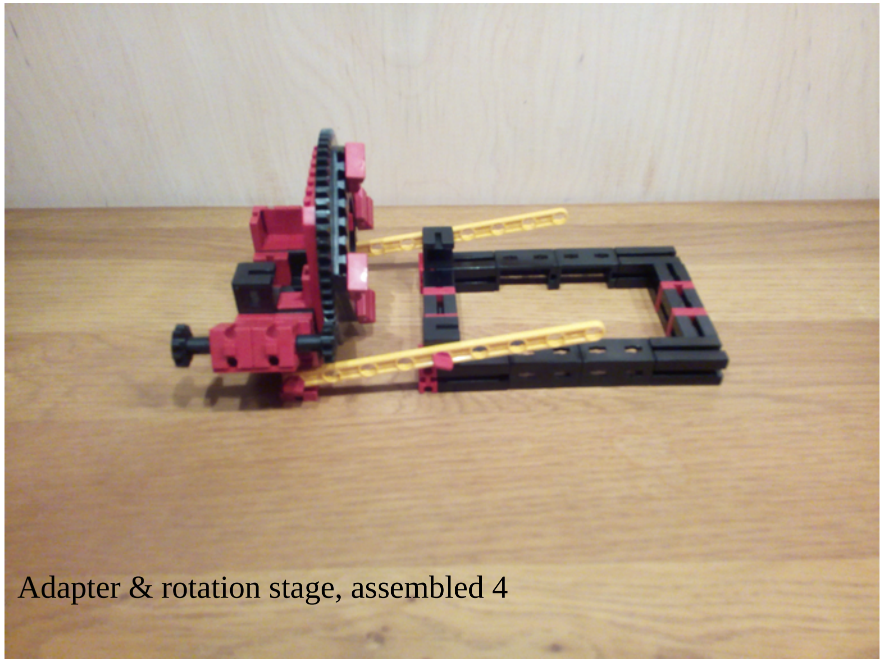

Adapter & rotation stage, assembled 4

Adapter & rotation stage, assembled 5

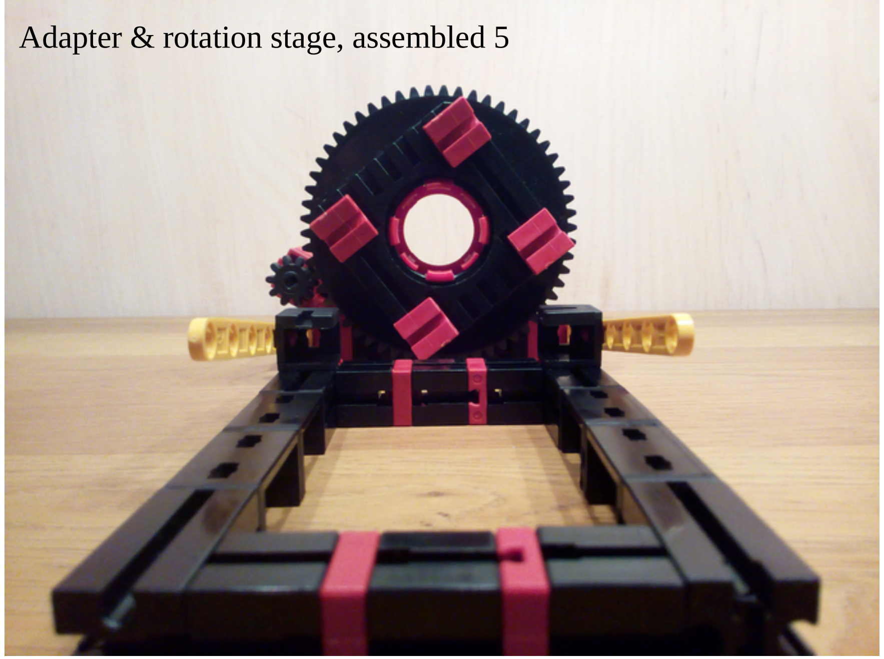
