## Supplementary material for "GraviKit: an easy-to-implement microscope add-on for observation of gravitation dependent processes": chamberPrepVerticalImaging (sample preparation)

**Creating a liquid chamber for vertical microscopy:**

Here‘s what you need (apart from your sample):

- a slide
- a coverslip (ideally 22mm wide)
- scissors
- sharp knife/ scalpel/ razor blade
- forceps
- tape (3M™ Montageband wiederlösbar, Transparent, 19 mm x 5 m, 0,8 mm, 3M-ID 7000073335), this is a double-sided transparent tape with a protective foil on 1 side
- some paper

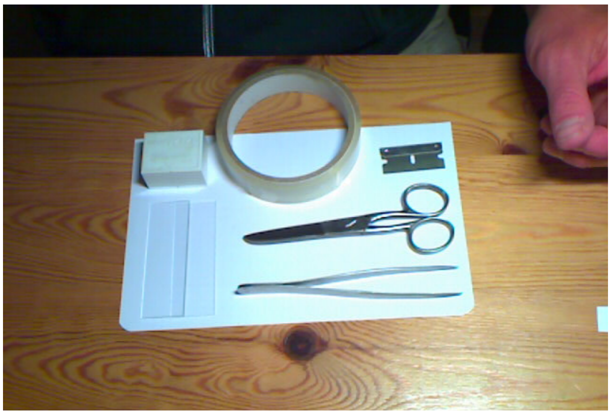

First you cut a piece of paper to the size of the desired chamber. Then you put that piece of paper on the tape (on the side without the protective foil) to make it not stick on the area where your sample chamber will be. Do not remove the protective foil.

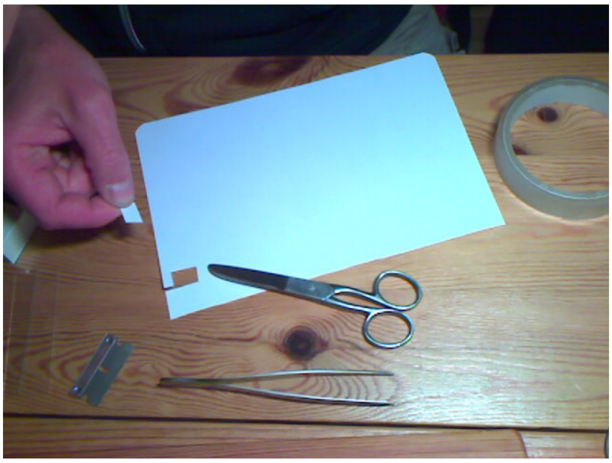

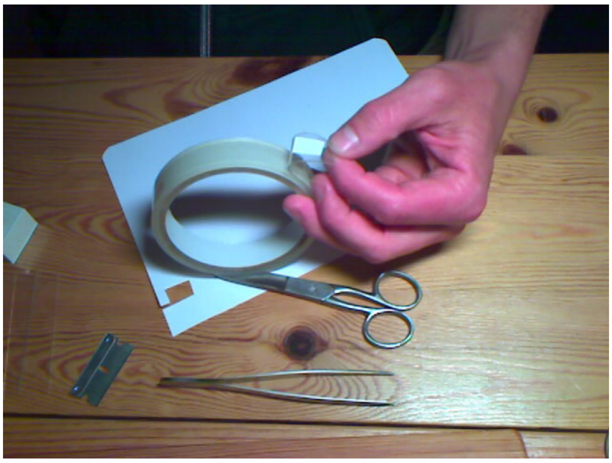

You cut off the piece of tape carrying the paper leaving enough edge behind the paper for sticking to the slide. Then you put the piece of tape onto the slide (paper facing the slide) and press firmly onto the protective foil.

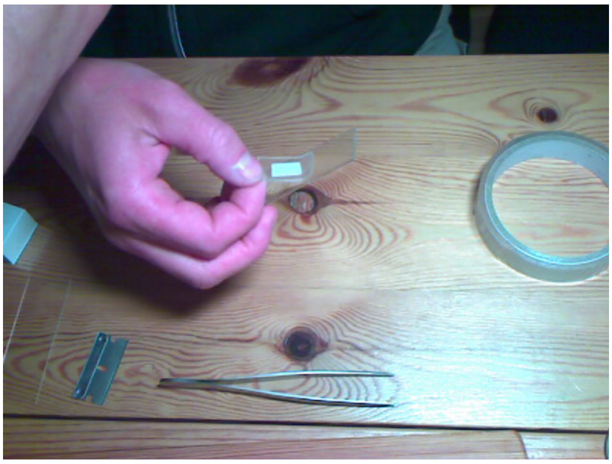

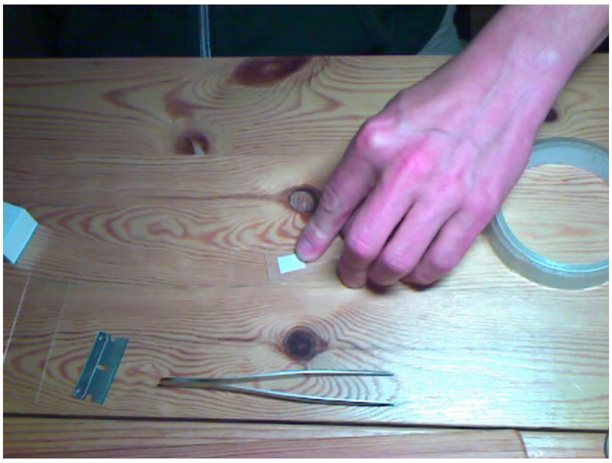

Christian Feldhaus, MPI for Developmental Biology

cc BY SA

The next step is to cut out the part where the paper is with a sharp knife. It is tricky but make sure you cut down to the glass all around the paper. This makes it easier to remove the part tape with the paper in the following step using the forceps.

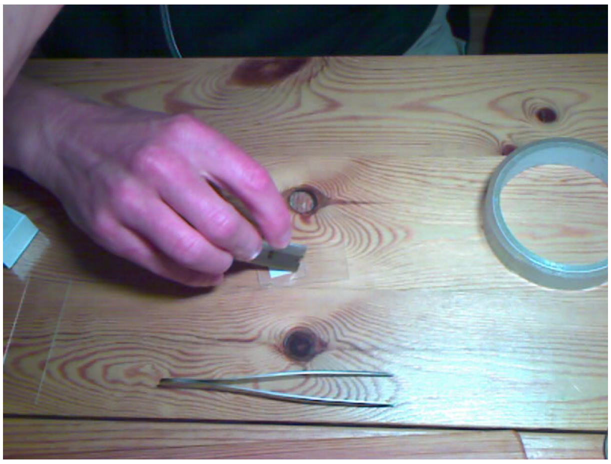
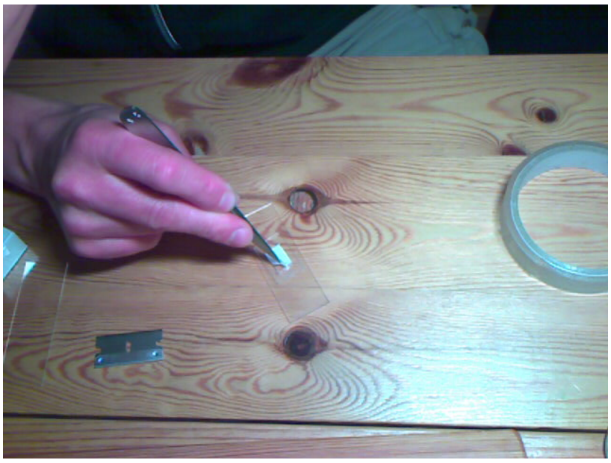

Now you have your chamber ready and you can remove the protective foil before …

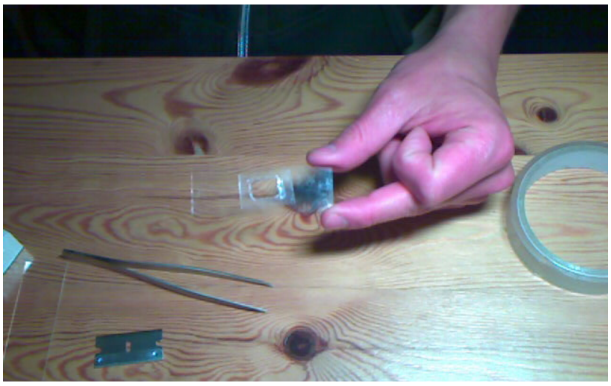
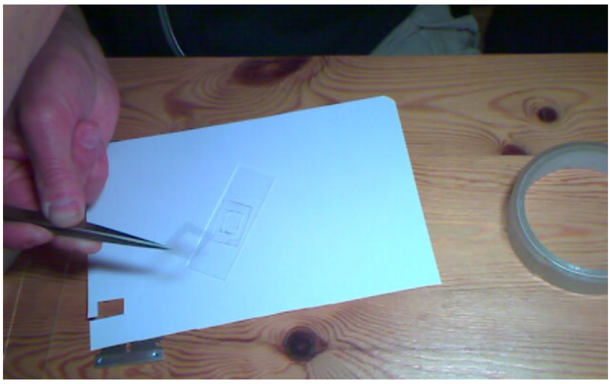

you can fill in your sample and put the coverslip on top.

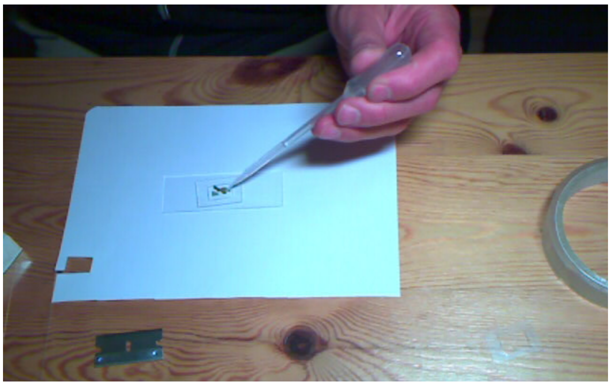
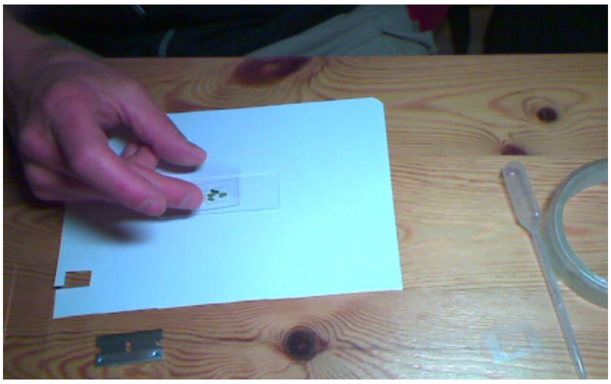

This last step is also quite tricky, especially for a closed chamber. What we do for e.g. plants is that we have an opening on one side (during preparation the paper goes over the edge of the tape), we put the sample on a drop of agarose, place the coverslip and then fill up with liquid. You could also after filling the liquid close the opening with e.g. another small piece of tape or wax.

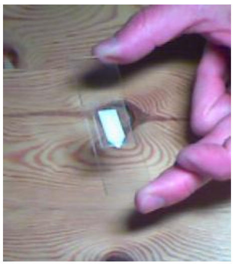

Christian Feldhaus, MPI for Developmental Biology
