## Supplementary material for "GraviKit: an easy-to-implement microscope add-on for observation of gravitation dependent processes": LiteratureRootImaging (suppl. table)

| Articles in "Plant Methods" | date | doi | original | review | vert. beampath | horiz. beampath |
|---|---|---|---|---|---|---|
| Fröschel, C. In-depth evaluation of root infection systems using the vascular fungus *Verticillium longisporum* as soil-borne model pathogen. *Plant Methods* **17,** 57 (2021). | 05 June 2021 | https://doi.org/1( | 1 | 0 | 1 | 0 |
| Cheng, Y., Wang, X., Cao, L. *et al.* Highly efficient *Agrobacterium rhizogenes*-mediated hairy root transformation for gene functional and gene editing analysis in soybean. *Plant Methods* **17,** 73 (2021). | 10 July 2021 | https://doi.org/1( | 1 | 0 | 1 | 0 |
| Wang, Y., Yang, F., Zhu, PF. *et al.* Use of the rhizobial type III effector gene *nopP* to improve *Agrobacterium rhizogenes*-mediated transformation of *Lotus japonicus*. *Plant Methods* **17,** 66 (2021). | 23 June 2021 | https://doi.org/1( | 1 | 0 | 1 | 0 |
| Xu, H., Zhao, Y., Suo, Y. *et al.* A label-free, fast and high-specificity technique for plant cell wall imaging and composition analysis. *Plant Methods* **17,** 29 (2021). | 19 March 2021 | https://doi.org/1( | 1 | 0 | 1 | 0 |
| Amarteifio, S., Fallesen, T., Pruessner, G. *et al.* A random-sampling approach to track cell divisions in time-lapse fluorescence microscopy. *Plant Methods* **17,** 25 (2021). | 08 March 2021 | https://doi.org/1( | 1 | 0 | 0 | 1 |
| Harrington, S.A., Backhaus, A.E., Fox, S. *et al.* A heat-shock inducible system for flexible gene expression in cereals. *Plant Methods* **16,** 137 (2020). | 14 October 2020 | https://doi.org/1( | 1 | 0 | 1 | 0 |
| Zhang, L., Zhao, Y., Liang, H. *et al.* Gateway-compatible vectors for functional analysis of proteins in cell type specific manner. *Plant Methods* **16,** 93 (2020). | 06 July 2020 | https://doi.org/1( | 1 | 0 | 1 | 0 |
| Zhang, K., He, J., Liu, L. *et al.* A convenient, rapid and efficient method for establishing transgenic lines of *Brassica napus*. *Plant Methods* **16,** 43 (2020). | 30 March 2020 | https://doi.org/1( | 1 | 0 | 1 | 0 |
| Osman, M., Stigloher, C., Mueller, M.J. *et al.* An improved growth medium for enhanced inoculum production of the plant growth-promoting fungus *Serendipita indica*. *Plant Methods* **16,** 39 (2020). | 16 March 2020 | https://doi.org/1( | 1 | 0 | 1 | 0 |
| Tofanelli, R., Vijayan, A., Scholz, S. *et al.* Protocol for rapid clearing and staining of fixed Arabidopsis ovules for improved imaging by confocal laser scanning microscopy. *Plant Methods* **15,** 120 (2019). | 25 October 2019 | https://doi.org/1( | 1 | 0 | 1 | 0 |

| Articles in "Nature Plants" | date | doi | original | review | vert. beampath | horiz. beampath |
|---|---|---|---|---|---|---|
| Serre, N.B.C., Kralík, D., Yun, P. et al. AFB1 controls rapid auxin signalling through membrane depolarization in Arabidopsis thaliana root. Nat. Plants (2021). | 19 July 2021 | https://dc | 1 | 0 | 0 | 1 |
| Kumari, P., Dahiya, P., Livanos, P. *et al.* IQ67 DOMAIN proteins facilitate preprophase band formation and division-plane orientation. *Nat. Plants* **7,** 739–747 (2021). | 24 May 2021 | https://dc | 1 | 0 | 1 | 0 |
| Zhang, Y., Mitsuda, N., Yoshizumi, T. *et al.* Two types of bHLH transcription factor determine the competence of the pericycle for lateral root initiation. *Nat. Plants* **7,** 633–643 (2021). | 18 May 2021 | https://dc | 1 | 0 | 1 | 0 |
| Vukašinović, N., Wang, Y., Vanhoutte, I. *et al.* Local brassinosteroid biosynthesis enables optimal root growth. *Nat. Plants* **7,** 619–632 (2021). | 17 May 2021 | https://dc | 1 | 0 | 1 | 1 |
| Doumane, M., Lebecq, A., Colin, L. *et al.* Inducible depletion of PI(4,5)$P_2$ by the synthetic iDePP system in *Arabidopsis*. *Nat. Plants* **7,** 587–597 (2021). | 17 May 2021 | https://dc | 1 | 0 | 1 | 0 |
| Ursache, R., De Jesus Vieira Teixeira, C., Dénervaud Tendon, V. *et al.* GDSL-domain proteins have key roles in suberin polymerization and degradation. *Nat. Plants* **7,** 353–364 (2021). | 08 March 2021 | https://dc | 1 | 0 | 1 | 0 |
| Jiang, J., Liu, J., Sanders, D. *et al.* UVR8 interacts with de novo DNA methyltransferase and suppresses DNA methylation in *Arabidopsis*. *Nat. Plants* **7,** 184–197 (2021). | 25 January 2021 | https://dc | 1 | 0 | 1 | 0 |
| Kulich, I., Vogler, F., Bleckmann, A. *et al.* ARMADILLO REPEAT ONLY proteins confine Rho GTPase signalling to polar growth sites. *Nat. Plants* **6,** 1275–1288 (2020). | 05 October 2020 | https://dc | 1 | 0 | 1 | 0 |
| Meier, M., Liu, Y., Lay-Pruitt, K.S. *et al.* Auxin-mediated root branching is determined by the form of available nitrogen. *Nat. Plants* **6,** 1136–1145 (2020). | 11 September 202( | https://dc | 1 | 0 | 1 | 0 |
| Matosevich, R., Cohen, I., Gil-Yarom, N. *et al.* Local auxin biosynthesis is required for root regeneration after wounding. *Nat. Plants* **6,** 1020–1030 (2020). | 03 August 2020 | https://dc | 1 | 0 | 1 | 0 |

| Articles in "Plant Cell" | date | doi | original | review | vert. beampath | horiz. beampath |
|---|---|---|---|---|---|---|
| Lise C Noack, Vincent Bayle, Laia Armengot, Frédérique Rozier, Adiilah Mamode-Cassim, Floris D Stevens, Marie-Cécile Caillaud, Teun Munnik, Sébastien Mongrand, Roman Pleskot, Yvon Jaillais, A nanodomain-anchored scaffolding complex is required for the function and localization of phosphatidylinositol 4-kinase alpha in plants, *The Plant Cell*, 2021 | 19 May 2021 | https://do | 1 | 0 | 1 | 0 |
| Sabrina Chin, Taegun Kwon, Bibi Rafeiza Khan, J Alan Sparks, Eileen L Mallery, Daniel B Szymanski, Elison B Blancaflor, Spatial and temporal localization of SPIRRIG and WAVE/SCAR reveal roles for these proteins in actin-mediated root hair development, *The Plant Cell*, 2021 | 20 April 2021 | https://do | 1 | 0 | 1 | 0 |
| Samuel W H Koh, Petra Marhava, Surbhi Rana, Alina Graf, Bernard Moret, Alkistis E L Bassukas, Melina Zourelidou, Martina Kolb, Ulrich Z Hammes, Claus Schwechheimer, Christian S Hardtke, Mapping and engineering of auxin-induced plasma membrane dissociation in BRX family proteins, *The Plant Cell*, Volume 33, Issue 6, June 2021, Pages 1945–1960 | 05 March 2021 | https://do | 1 | 0 | 1 | 0 |
| Kaija Goodman, Julio Paez-Valencia, Janice Pennington, Annika Sonntag, Xinxin Ding, Han Nim Lee, Paul G Ahlquist, Isabel Molina, Marisa S Otegui, ESCRT components ISTL1 andLIP5 are required for tapetal function and pollen viability, *The Plant Cell*, 2021 | 10 May 2021 | https://do | 1 | 0 | 1 | 0 |
| Noemi Ruiz-Lopez, Jessica Pérez-Sancho, Alicia Esteban del Valle, Richard P Haslam, Steffen Vanneste, Rafael Catalá, Carlos Perea-Resa, Daniël Van Damme, Selene García-Hernández, Armando Albert, José Vallarino, Jinxing Lin, Jiří Friml, Alberto P Macho, Julio Salinas, Abel Rosado, Johnathan A Napier, Vitor Amorim-Silva, Miguel A Botella, Synaptotagmins at theendoplasmic reticulum–plasma membrane contact sites maintain diacylglycerol homeostasis during abiotic stress, *The Plant Cell*, 2021 | 04 May 2021 | https://do | 1 | 0 | 1 | 0 |
| Xi Yang, Weiguo Dong, Wenqing Ren, Qiuxia Zhao, Feijie Wu, Yuke He, Cytoplasmic HYL1 modulates miRNA-mediated translational repression, *The Plant Cell*, Volume 33, Issue 6, June 2021, Pages 1980–1996 | 25 March 2021 | https://do | 1 | 0 | 1 | 0 |
| Akira Yoshinari, Takuya Hosokawa, Marcel Pascal Beier, Keishi Oshima, Yuka Ogino, Chiaki Hori, Taichi E Takasuka, Yoichiro Fukao, Toru Fujiwara, Junpei Takano, Transport-coupled ubiquitination of the borate transporter BOR1 for its boron-dependent degradation, *The Plant Cell*, Volume 33, Issue 2, February 2021, Pages 420–438 | 03 December 20 | https://do | 1 | 0 | 1 | 0 |
| Carla Brillada, Ooi-Kock Teh, Franck Anicet Ditengou, Chil-Woo Lee, Till Klecker, Bushra Saeed, Giulia Furlan, Marco Zietz, Gerd Hause, Lennart Eschen-Lippold, Wolfgang Hoehenwarter, Justin Lee, Thomas Ott, Marco Trujillo, Exocyst subunit Exo70B2 is linked to immune signaling and autophagy, *The Plant Cell*, Volume 33, Issue 2, February 2021, Pages 404–419 | 03 December 20 | https://do | 1 | 0 | 1 | 0 |
| Xiu-Li Hou, Wen-Qiang Chen, Yifeng Hou, Hua-Qin Gong, Jing Sun, Zhen Wang, Heng Zhao, Xiaofeng Cao, Xiu-Fen Song, Chun-Ming Liu, DEAD-BOX RNA HELICASE 27 regulates microRNA biogenesis, zygote division, and stem cell homeostasis, *The Plant Cell*, Volume 33, Issue 1, January 2021, Pages 66–84 | 17 November 2( | https://do | 1 | 0 | 1 | 0 |
| Jing Liu, Miao Xia Liu, Li Ping Qiu, Fang Xie, SPIKE1 Activates the GTPase ROP6 to Guide the Polarized Growth of Infection Threads in *Lotus japonicus* , *The Plant Cell*, Volume 32, Issue 12, December 2020, Pages 3774–3791 | 06 October 202( | https://do | 1 | 0 | 1 | 0 |

| Articles in "Plant Physiology" | date | doi | original | review | vert. beampath | horiz. beampath |
|---|---|---|---|---|---|---|
| José Manuel Ugalde, Michelle Schlößer, Armelle Dongois, Alexandre Martinière, Andreas J Meyer, The latest HyPe(r) in plant $H_2O_2$ biosensing, *Plant Physiology*, 2021 | 02 July 2021 | https://do | 1 | 0 | 1 | 0 |
| Taro Kimura, Ken Haga, Yuko Nomura, Takumi Higaki, Hirofumi Nakagami, Tatsuya Sakai, Phosphorylation of NONPHOTOTROPIC HYPOCOTYL3 affects photosensory adaptation during the phototropic response, *Plant Physiology*, 2021 | 17 June 2021 | https://do | 1 | 0 | 1 | 0 |
| Francesca Resentini, Matteo Grenzi, Daniele Ancora, Mara Cademartori, Laura Luoni, Marianna Franco, Andrea Bassi, Maria Cristina Bonza, Alex Costa, Simultaneous imaging of ER and cytosolic $Ca^{2+}$ dynamics reveals long-distance ER $Ca^{2+}$ waves in plants, *Plant Physiology*, 2021 | 27 May 2021 | https://do | 1 | 0 | 1 | 0 |
| Sandra E Zimmermann, Ruben M Benstein, María Flores-Tornero, Samira Blau, Armand D Anoman, Sara Rosa-Téllez, Silke C Gerlich, Mohamed A Salem, Saleh Alseekh, Stanislav Kopriva, Vera Wewer, Ulf-Ingo Flügge, Richard P Jacoby, Alisdair R Fernie, Patrick Giavalisco, Roc Ros, Stephan Krueger, The phosphorylated pathway of serine biosynthesis links plant growth with nitrogen metabolism, *Plant Physiology*, Volume 186, Issue 3, July 2021, Pages 1487–1506 | 26 May 2021 | https://do | 1 | 0 | 1 | 0 |
| Zachary G Beamer, Pratyush Routray, Won-Gyu Choi, Margaret K Spangler, Ansul Lokdarshi, Daniel M Roberts, Aquaporin family lactic acid channel NIP2;1 promotes plant survival under low oxygen stress in Arabidopsis, *Plant Physiology*, 2021 | 07 May 2021 | https://do | 1 | 0 | 1 | 0 |
| Yeling Zhou, Yuzhu Wang, Jingwen Li, Jiansheng Liang, *In vivo* FRET–FLIM reveals ER-specific increases in the ABA level upon environmental stresses, *Plant Physiology*, Volume 186, Issue 3, July 2021, Pages 1545–1561 | 12 April 2021 | https://do | 1 | 0 | 1 | 0 |
| Thomas D Alcock, Catherine L Thomas, Seosamh Ó Lochlainn, Paula Pongrac, Michael Wilson, Christopher Moore, Guilhem Reyt, Katarina Vogel-Mikuš, Mitja Kelemen, Rory Hayden, Lolita Wilson, Pauline Stephenson, Lars Østergaard, Judith A Irwin, John P Hammond, Graham J King, David E Salt, Neil S Graham, Philip J White, Martin R Broadley, Magnesium and calcium overaccumulate in the leaves of a *schengen3* mutant of *Brassica rapa*, *Plant Physiology*, Volume 186, Issue 3, July 2021, Pages 1616–1631 | 08 April 2021 | https://do | 1 | 0 | 1 | 0 |
| Guichen Li, Zitong Li, Zeyun Yang, Yehoram Leshem, Yuequan Shen, Shuzhen Men, Mitochondrial heat-shock cognate protein 70 contributes to auxin-mediated embryo development, *Plant Physiology*, Volume 186, Issue 2, June 2021, Pages 1101–1121 | 21 March 2021 | https://do | 1 | 0 | 1 | 0 |
| Madhumitha Narasimhan, Michelle Gallei, Shutang Tan, Alexander Johnson, Inge Verstraeten, Lanxin Li, Lesia Rodriguez, Huibin Han, Ellie Himschoot, Ren Wang, Steffen Vanneste, Judit Sánchez-Simarro, Fernando Aniento, Maciek Adamowski, Jiří Friml, Systematic analysis of specific and nonspecific auxin effects on endocytosis and trafficking, *Plant Physiology*, Volume 186, Issue 2, June 2021, Pages 1122–1142 | 18 March 2021 | https://do | 1 | 0 | 0 | 1 |
| Laining Zhang, Tetyana Smertenko, Deirdre Fahy, Nuria Koteyeva, Natalia Moroz, Anna Kuchařová, Dominik Novák, Eduard Manoilov, Petro Smertenko, Charitha Galva, Jozef Šamaj, Alla S. Kostyukova, John C. Sedbrook, Andrei Smertenko, Analysis of formin functions during cytokinesis using specific inhibitor SMIFH2, *Plant Physiology*, Volume 186, Issue 2, June 2021, Pages 945–963 | 23 February 202 | https://do | 1 | 0 | 1 | 0 |

| Articles in "The Plant Journal" | date | doi | original | review | vert. beampath | horiz. beampath |
|---|---|---|---|---|---|---|
| Li, C., Liu, G., Geng, X., He, C., Quan, T., Hayashi, K.-I., De Smet, I., Robert, H.S., Ding, Z. and Yang, Z.-B. (2021), Local regulation of auxin transport in root-apex transition zone mediates aluminium-induced Arabidopsis root-growth inhibition. Plant J | 17 July 2021 | https://do | 1 | 0 | 1 | 0 |
| Brasileiro, A.C.M., Lacorte, C., Pereira, B.M., Oliveira, T.N., Ferreira, D.S., Mota, A.P.Z., Saraiva, M.A.P., Araujo, A.C.G., Silva, L.P. and Guimaraes, P.M. (2021), Ectopic expression of an *expansin-like B* gene from wild *Arachis* enhances tolerance to both abiotic and biotic stresses. Plant J | 07 July 2021 | https://do | 1 | 0 | 1 | 0 |
| Fu, J., Zhang, X., Liu, J., Gao, X., Bai, J., Hao, Y. and Cui, H. (2021), A mechanism coordinating root elongation, endodermal differentiation, redox homeostasis and stress response. Plant J | 31 May 2021 | https://do | 1 | 0 | 1 | 0 |
| Li, J., Wang, Y., Zou, W., Jian, L., Fu, Y. and Zhao, J. (2021), *AtNUF2* modulates spindle microtubule organization and chromosome segregation during mitosis. Plant J | 16 May 2021 | https://do | 1 | 0 | 1 | 0 |
| Nisa, M., Bergis, C., Pedroza-Garcia, J.-A., Drouin-Wahbi, J., Mazubert, C., Bergounioux, C., Benhamed, M. and Raynaud, C. (2021), The plant DNA polymerase theta is essential for the repair of replication-associated DNA damage. Plant J, 106: 1197-1207 | 14 May 2021 | https://do | 1 | 0 | 1 | 0 |
| Chen, K., Su, C., Tang, W., Zhou, Y., Xu, Z., Chen, J., Li, H., Chen, M. and Ma, Y. (2021), Nuclear transport factor GmNTF2B-1 enhances soybean drought tolerance by interacting with oxidoreductase GmOXR17 to reduce reactive oxygen species content. Plant J. | 12 May 2021 | https://do | 1 | 0 | 1 | 0 |
| Zhang, C., He, M., Wang, S., Chu, L., Wang, C., Yang, N., Ding, G., Cai, H., Shi, L. and Xu, F. (2021), Boron deficiency-induced root growth inhibition is mediated by brassinosteroid signalling regulation in Arabidopsis. Plant J. | 08 May 2021 | https://do | 1 | 0 | 1 | 0 |
| Kim, L.J., Tsuyuki, K.M., Hu, F., Park, E.Y., Zhang, J., Iraheta, J.G., Chia, J.-C., Huang, R., Tucker, A.E., Clyne, M., Castellano, C., Kim, A., Chung, D.D., DaVeiga, C.T., Parsons, E.M., Vatamaniuk, O.K. and Jeong, J. (2021), Ferroportin 3 is a dual-targeted mitochondrial/chloroplast iron exporter necessary for iron homeostasis in Arabidopsis. Plant J, 107: 215-236 | 22 April 2021 | https://do | 1 | 0 | 1 | 0 |
| Castaings, L., Alcon, C., Kosuth, T., Correia, D. and Curie, C. (2021), Manganese triggers phosphorylation-mediated endocytosis of the Arabidopsis metal transporter NRAMP1. Plant J, 106: 1328-1337. | 18 March 2021 | https://do | 1 | 0 | 1 | 0 |
| Ding, T., Zhang, F., Wang, J., Wang, F., Liu, J., Xie, C., Hu, Y., Shani, E., Kong, X., Ding, Z. and Tian, H. (2021), Cell-type action specificity of auxin on *Arabidopsis* root growth. Plant J, 106: 928-941 | 20 February 202 | https://do | 1 | 0 | 1 | 0 |
